# Single Cell Transcriptomics identifies a WNT7A-FZD5 Signaling Axis that maintains Fallopian Tube Stem Cells in Patient-derived Organoids

**DOI:** 10.1101/2022.08.02.502319

**Authors:** Abdulkhaliq Alsaadi, Mara Artibani, Zhiyuan Hu, Nina Wietek, Matteo Morotti, Laura Santana Gonzales, Moiad Alazzam, Jason Jiang, Beena Abdul, Hooman Soleymani majd, Levi L Blazer, Jarret Adams, Francesca Silvestri, Sachdev S Sidhu, Joan S. Brugge, Ahmed Ashour Ahmed

## Abstract

Despite its significance to reproduction, fertility, sexually transmitted infections and various pathologies, the fallopian tube (FT) is relatively understudied. Strong evidence points to the FT as the tissue-of-origin of high grade serous ovarian cancer (HGSOC), the most fatal gynaecological malignancy. HGSOC precursor lesions arise specifically in the distal FT (fimbria) which is reported to be enriched in stem-like cells. Investigation of the role of FT stem cells in health and disease has been hampered by a lack of characterization of FT stem cells and lack of models that recapitulate stem cell renewal and differentiation *in vitro*. Using optimized organoid culture conditions to address these limitations, we found that FT stem cell renewal is highly dependent on WNT/β-catenin signaling and engineered endogenous WNT/β-catenin signaling reporter organoids to biomark, isolate and characterize putative FT stem cells. Using functional approaches as well as bulk and single cell transcriptomic analyses, we show that an endogenous hormonally-regulated WNT7A-FZD5 signaling axis is critical for self-renewal of human FT stem cells, and that WNT/β-catenin pathway-activated FT cells form a distinct transcriptomic cluster of cells enriched in ECM remodelling and integrin signaling pathways. In addition, we find that the WNT7A-FZD5 signaling axis is dispensable for mouse oviduct regeneration. Overall, we provide a deep characterization of FT stem cells and their molecular requirements for self-renewal, paving the way for mechanistic work investigating the role of stem cells in FT health and disease.

**GRAPHICAL ABSTRACT:** 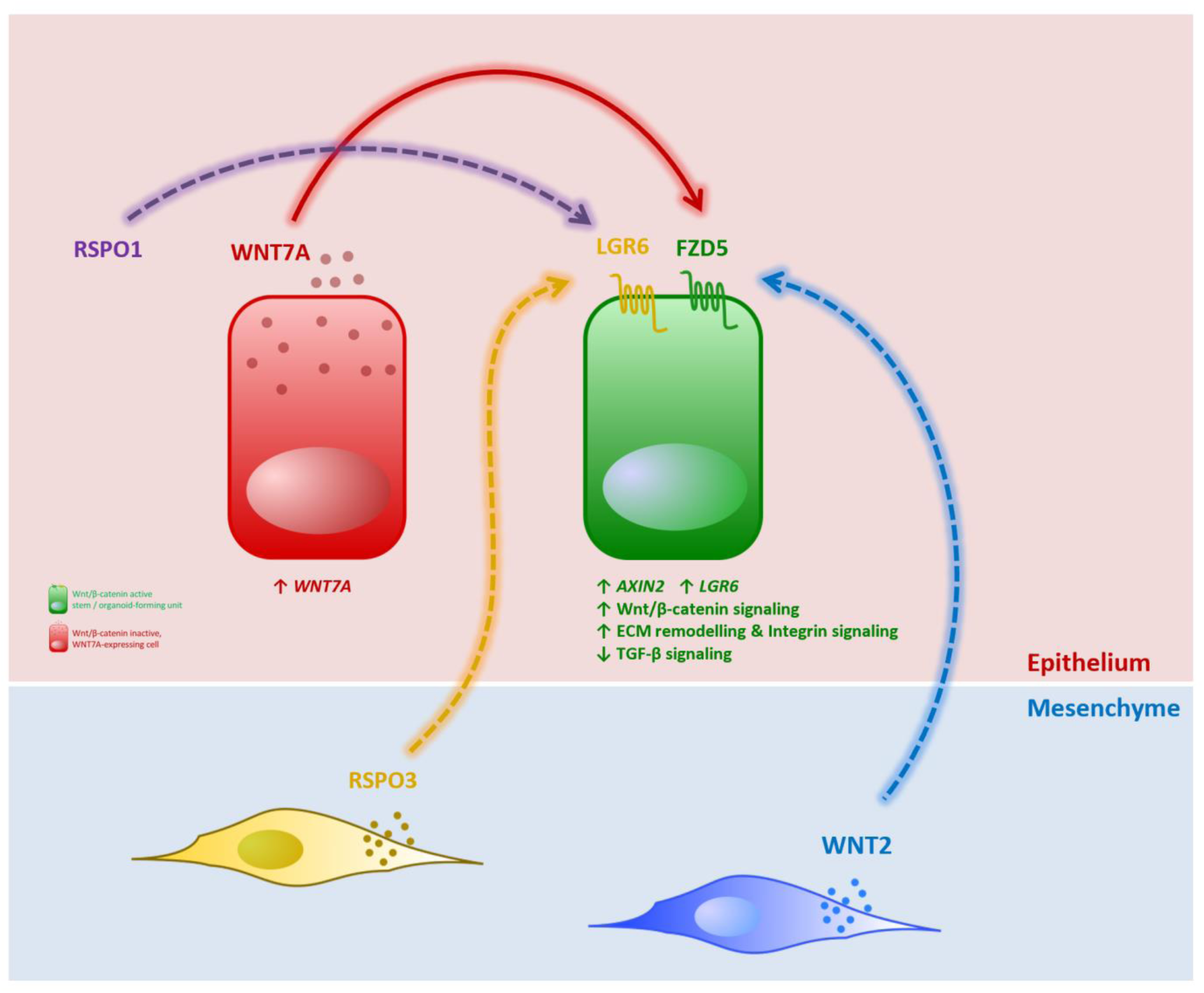

## INTRODUCTION

The human fallopian tube (hFT) is lined with pseudo-stratified columnar epithelium composed of PAX8+ MYC+ secretory cells that nourish released eggs / zygotes, as well as TUBB4+ FOXJ1+ ciliated cells that beat rhythmically to facilitate ovum/zygote transport to the uterine cavity. Despite its significance to fertility, reproduction and women’s health and disease, little is known about the FT’s biology, cellular hierarchy and homeostasis which are critical for understanding infertility disorders, ectopic pregnancy, sexually transmitted infections and FT-derived cancers. A number of studies have attempted to bridge this knowledge gap, showing that the distal hFT and the distal region of its murine equivalent, the mouse oviduct (mOV), are enriched in stem-like cells possessing longevity and multipotency (Erickson *et al*., 2013; Ghosh *et al*., 2017; Xie *et al*., 2018; Wang *et al*., 2012; Zhu *et al*., 2020). Studies in mice identified a population of label-retaining cells (LRCs) at the distal mOV (Snegovskikh *et al*., 2014) that are enhanced for differentiated spheroid formation (Wang *et al*., 2012; Xie *et al*., 2018).

However, strong evidence for the existence of mOV stem cells came from *in vivo* lineage tracing that employed doxycycline-inducible labelling of secretory cells using a Pax8^rtTA^ TetO^Cre^ YFP^fl/fl^ mouse model, demonstrating that ciliated cells emerge from secretory cells (Ghosh *et al*., 2017). Similarly in the hFT, putative stem cells were shown to be secretory in nature, using spheroid (Paik *et al*., 2012), air-liquid interface (Yamamoto *et al*., 2016) and organoid-based approaches (Kessler *et al*., 2015). Although this points to secretory cells as drivers of stem cell activity, we and others have recently uncovered a previously unappreciated heterogeneity within the FT secretory compartment (Hu *et al*., 2020; Dinh *et al*., 2021), using single cell transcriptomic (SCT) profiling of fresh hFT tissue. Therefore, although *in vivo* and other studies refined the search for hFT/mOV stem cells, no studies have successfully pinpointed the secretory cell type driving FT renewal, due to limitations in model tractability and difficulty in cell biomarking and isolation. Wnt/β-catenin signaling (WβS) is thought to be involved in hFT/mOV self-renewal (Kessler *et al*., 2015; Ghosh *et al*., 2017), but the precise molecular mechanisms remain unknown. Furthermore, humans and great apes possess FTs whereas the equivalent in mice is the oviduct. Due to major anatomical differences, biological differences are likely to exist, and no studies have scrutinized whether mOV biology is representative to the hFT.

In addition to being a site where stem-like cells concentrate as abovementioned, mounting evidence from several clinical and *in vivo* studies point to the distal FT fimbria as the site of origin of high grade serous ovarian cancer or HGSOC (Labidi-Galy *et al*., 2017; Kuhn *et al*., 2012; Kim *et al*., 2018). HGSOC is a fatal gynaecological malignancy with a dismal 5-year survival of 25% in late stage disease (Kroeger & Drapkin, 2017). Understanding FT renewal may shed light on HGSOC initiation mechanisms, which remain poorly understood, further reinforcing the urgency of this investigation. Here, we harness three-dimensional patient-derived FT organoids to model FT regeneration *in vitro*. We genetically label, isolate and characterize putative FT stem cells using functional approaches as well as bulk and SCT analyses, identifying a hormonally regulated WNT7A-FZD5 signaling axis that is critical for FT stem cell maintenance.

## RESULTS

### Optimizing robust regeneration of hFT organoids from single cells

Existing models of FT biology suffer from drawbacks that limit their utility in characterizing FT stem cells (Figure S1A). For example, conventional two-dimensional (2D) cultures of primary FT cells (Figure S1 B-D) lose epithelial markers within 5-6 weeks of culture (Figure S1E) and were previously shown to lack ciliated cells (Hu *et al*., 2020). In contrast, organoid cultures were shown to be robust hFT/mOV models incorporating PAX8+ secretory and TUBB4+ ciliated cells (Kessler *et al*., 2015; Xie *et al*., 2018) and we reproduced this using the reported culture conditions (Figure 1A). However, a major limitation of the human organoid culture is the lack of organoid regeneration after single cell dissociation of patient tissue or organoids; a pre-requisite for isolating and characterizing putative progenitor cells. The TGF-β pathway promotes stem cell differentiation (Sakaki-Yumoto *et al*., 2013) and its inhibition is critical for regeneration of organoids from various tissues (Sato *et al*., 2009; Bartfeld *et al*., 2015; Karthaus *et al*., 2014; Turco *et al*., 2017). We reasoned that TGF-β pathway inhibitors contained in previously published culture conditions, including NOGGIN which inhibits BMP-activated SMAD1/5/8-mediated signaling, and SB431542 which is a ALK4/5/7 inhibitor that blocks SMAD 2/3 TGFβ-mediated signaling, do not sufficiently inhibit TGF-β signaling to suppress differentiation and enable organoid regeneration. To address this, we employed another ALK4/5/7 inhibitor, A83.01, which has 10-fold higher potency compared to SB431542 (Tojo *et al*., 2005) and found it to promote organoid formation efficiency (OFE) from single cells (Figures 1 B-C and S2 A-B). Interestingly, we found that TGF-β suppression was not necessary for regeneration of mouse oviduct (mOV) organoids from single cells (Figure S2 C-E).

**Figure 1:**
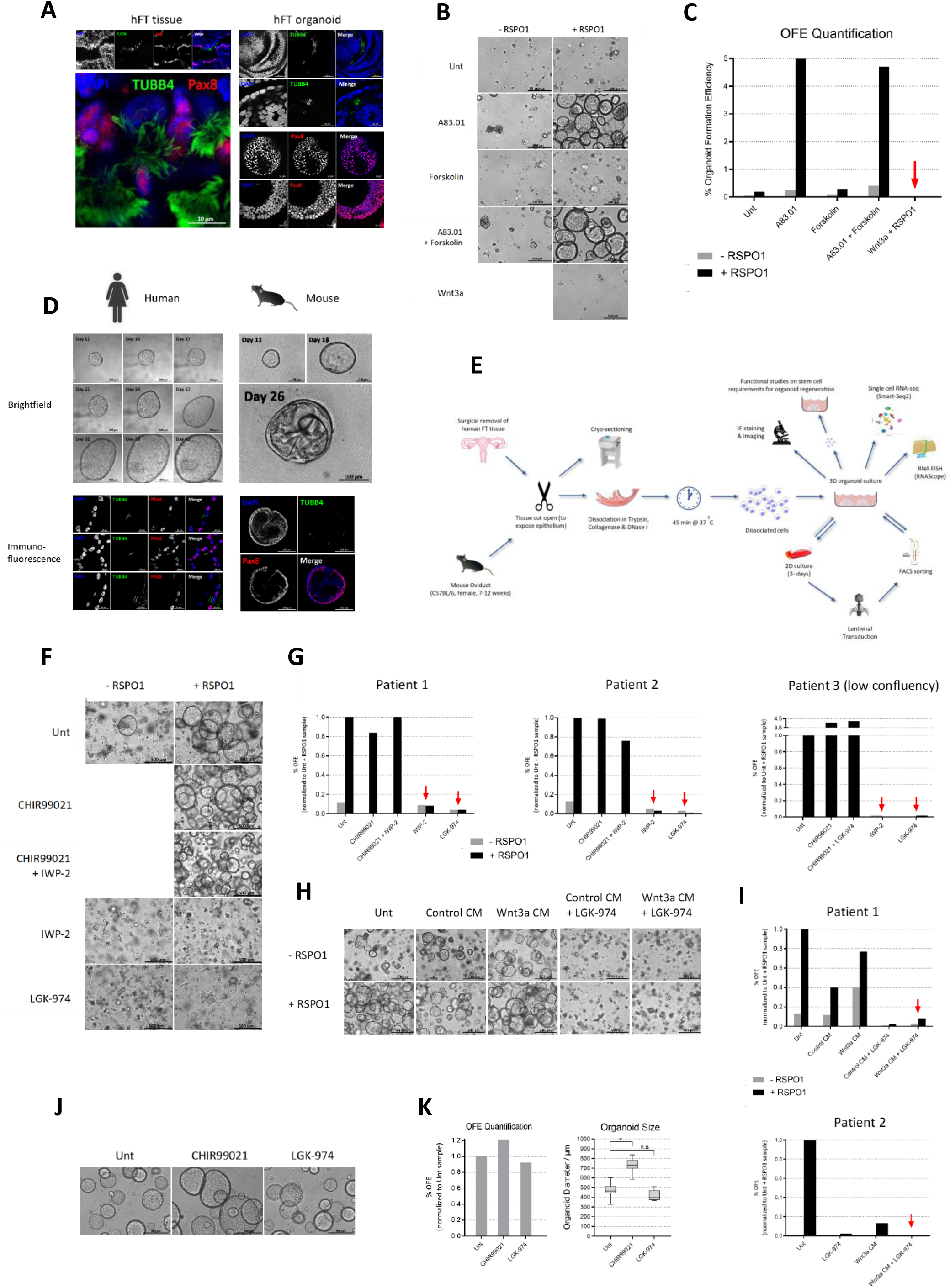
WNT/β-catenin signaling (WβS) is essential for organoid regeneration and is driven by unidentified endogenous WNT(s). **(A)** Confocal images of immunostaining for secretory cell marker PAX8 and ciliated cell marker TUBB4 in fresh frozen hFT sections or whole-mount immunostained hFT organoids. Organoids were grown for 10-13 days as described (Kessler et al., 2015). Representative of n = 3 patients. Scale bars as indicated. **(B-C)** TGF-β inhibitor A83.01 restores organoid formation efficiency (OFE) of single FT cells in an RSPO1-dependent manner **(B)** Representative brightfield images of EpCAM+ CD45-cells FACS-isolated from patient tissue and organoid-cultured for 12 days under the treatment conditions shown. 5,700 cells were plated in 50 μl Matrigel drops on 24-well plates. Scale bar, 500 μm. **(C)** Quantification of OFE for the samples shown in **(B)**. Red arrow points to conditions of the reported method for culturing FT organoids. Representative of 2 biological replicates (n=2 patients; second replicate, Figure S2 A-B). **(D)** Representative images of organoids cultured from single FACS-isolated EpCAM+ CD45-cells derived from established hFT or mOV organoid lines. Cells were cultured as single cells in 5 μl Matrigel drops in single wells of 96-well plates. Upper panel: brightfield images taken on the indicated days. Lower panel: representative confocal images of whole-mount immunostaining for differentiation markers. **(E)** Graphical summary of the experimental approaches employed to biomark and characterize putative hFT / mOV stem cells. **(F-G)** RSPO1 cooperates with an endogenous WNT(s) to drive organoid regeneration. **(F)** Representative brightfield images of p.1 hFT organoids after 8 days of the treatment with or without RSPO1, GSK-3β inhibitor CHIR99021 or/and Porcupine inhibitors IWP-2 or LGK-974. Scale bar, 500 μm. **(G)** Quantification of OFE for the samples shown in **(F)** plus 2 additional patient replicates. 19K, 12K or 7K cells were plated in 50 μl Matrigel drops on 24-well plates for patients 1, 2 or 3, respectively. Red arrows point to endogenous WNT secretion blockers abolishing OFE. P-values: (RSPO1) vs (RSPO1 + IWP-2) = 0.0006, (RSPO1) vs (RSPO1 + LGK-974) < 0.0001 (t-test, two-tailed, paired, n = 3 patient replicates). Four additional biological replicates (n=4 patients) for the effect of endogenous WNT secretion blockers on OFE are shown later in a different context (Figure 4 C-D). **(H-I)** WNT3A does not rescue regeneration of WNT-blocked organoids. **(H)** Representative brightfield images of hFT organoids after 8-9 days of the treatments shown. CM is conditioned medium. Scale bar, 500 μm. **(I)** Quantification of OFE for the samples shown in **(H)** and another patient replicate. Red arrows point to WNT3A’s failure to rescue endogenous WNT-blocked organoids. Activity of WNT3A CM was confirmed by the Wnt reporter (TOPFlash) assay (Figure S3A). **(J-K)** Regeneration of mOV organoids is WβS-independent. **(J)** Representative brightfield images of mOV organoids after 11 days of the treatments shown (in the absence of RSPO1). 2,000 cells were plated in 50 μl Matrigel drops on 24-well plates. Scale bar, 500 μm. **(K)** Quantification of OFE (left panel) and organoid size (right panel) of samples shown in **(J)**. OFE quantification (left panel) is representative of n = 5 biological replicates (2 replicates shown in Figure S2 F-G, 2 replicates shown later in a different context, Figure 4G). Right panel shows a box and whisker plot for the sizes of the 10 largest organoids per sample (centre line, median; box limits, upper and lower quartiles; whisker ends, minimum and maximum values). *P-value < 0.0001 (t-test, two-tailed).

In addition, we found that WNT3A conditioned medium (CM; Figure S3A) which is a basic component of the culture conditions of organoids from various tissues (see below), had no activity in hFT organoids and, indeed, reduced organoid regeneration in our optimized conditions (Figure S3B), most likely due to the presence of undefined serum factors which confound analysis of stem cell self-renewal requirements *in vitro*. Using our optimized conditions containing A83.01 / Forsklin and lacking WNT3A, we verified that organoids emerge only from epithelial cells (Figure S3C), display invaginations characteristic of hFT tissue (Figure S3D), possess a spherical tube-like structure with a hollow interior (Supplementary Movie 1) and contain a rare population of KI-67+ proliferative cells (Figure S3E). We also confirmed that single cells dissociated into individual wells formed organoids containing both PAX8+ and TUBB4+ differentiated progeny (Figure 1D), indicating organoids regeneration is driven by multipotent stem cells. Overall, the data above indicate that the serum-free culture conditions that we have optimized robustly support the regeneration of hFT organoids from single stem cells. We set out to characterize the FT organoid-forming stem cells using several different approaches (Figure 1E).

### Wnt/β-catenin signaling is essential for renewal of hFT stem cells

WNT/β-catenin signaling (herein referred to as WβS) was previously associated with hFT/mOV regeneration (Ghosh *et al*., 2017; Kessler *et al*., 2015; Xie *et al*., 2018). To investigate this further, we examined the effects of depletion or inhibition of multiple WβS pathway components on organoid regeneration. Withdrawal of RSPO1, which attenuates Wnt receptor turnover (Hao *et al*., 2012; Koo *et al*., 2012) reduced OFE by more than 90% (shown earlier, Figure 1 B-C), indicating that (WβS) is critical for stem cell renewal in organoids. To confirm this, we treated organoids with the tankyrase 1/2 inhibitor XAV-939 (Huang *et al*., 2009) and the β-catenin / CBP inhibitor PRI-724 (Okazaki *et al*., 2019) which block WβS at midstream (cytoplasmic) and downstream (nuclear) nodes, respectively (Figure S3F). Both treatments completely abolished organoid regeneration (Figure S3G). Furthermore, in our optimized serum-free setting, RSPO1 is the only component that augments WβS (Figure S3H). Since our culture medium does not contain a WNT source, we reasoned that WβS is activated via an endogenously secreted WNT in organoids. Indeed, blocking endogenous WNT secretion using Porcupine inhibitors (Figure S3I) reduced OFE by over 90% (Figure 1 F-G) phenocopying the effect of RSPO1 withdrawal (shown earlier, Figure 1 B-C) and suggesting that RSPO1 cooperates with an endogenous WNT to drive hFT organoid regeneration.

WNT3A is a widely used WNT source for WβS activation and expansion of organoids from various tissues, including hFT/mOV organoids (Hoffmann *et al*., 2020; Kessler *et al*., 2015, 2019; Kopper *et al*., 2019; Lõhmussaar *et al*., 2020). To address whether exogenous WNT3A could rescue organoid growth in cultures in which endogenous WNT secretion was blocked, we cultured ‘WNT-blocked’ (porcupine inhibitor-treated) organoids in the presence of WNT3A. The activity of WNT3A was validated using the Wnt reporter (TOPFlash) assay (shown earlier, Figure S3A). Contrary to the rescue of organoid regeneration seen in WNT-blocked intestinal organoids (Sato *et al*., 2011), WNT3A failed to rescue the regeneration of WNT-blocked hFT organoids (Figure 1 H-I) suggesting it cannot substitute for the endogenously secreted WNT ligand. In contrast to hFT organoids, the regeneration of mOV organoids was unaffected by blocking endogenous WNTs (Figures 1 J-K), suggesting mouse organoids renew using WβS-independent mechanisms.

### WβS-active cells drive organoid regeneration

The results above indicate that Wnt/β-catenin active (WβA) cells drive FT organoid regeneration. To isolate WβA cells, we transduced organoids from benign non-HGSOC patients with the WβS-reporter 7TGC lentiviral vector (Figure 2A), in which mCherry expression is driven by a constitutive (SV40) promoter and EGFP expression is driven by TCF/LEF promoter elements, also called WβS-Reporter Elements (WβS-RE). We expanded FACS-selected transduced cells using our optimized culture conditions, to generate stable WβS-reporter organoids (Figures 2B & S4A) from multiple patients. Confocal imaging of fixed (Figure 2C) and live (Figure S4B) organoids confirmed localized activation of WβS. In addition, WβA cells were PAX8+ / secretory in lineage (Figure 2D) and constituted 1.5-5% of all cells (Figure 2E and Figure S4A for mOV and hFT organoids, respectively), mirroring the proportion of organoid-forming units in hFT organoids. Blocking endogenous WNT secretion by LGK-974 treatment abolished EGFP+ cells (Figure 2E) while WβS activation using the GSK3 inhibitor CHIR99021 increased the proportion of EGFP+ cells over 10-fold (Figures 2E and S4C), suggesting EGFP faithfully marked WβA cells. AXIN2, a reliable marker of WβS activation in several organoid systems (Boonekamp *et al*., 2021), including hFT organoids (Figure S4D), was elevated in EGFP+ cells (Figure 2F), further validating EGFP+ cells as WβA cells in this setting.

**Figure 2:**
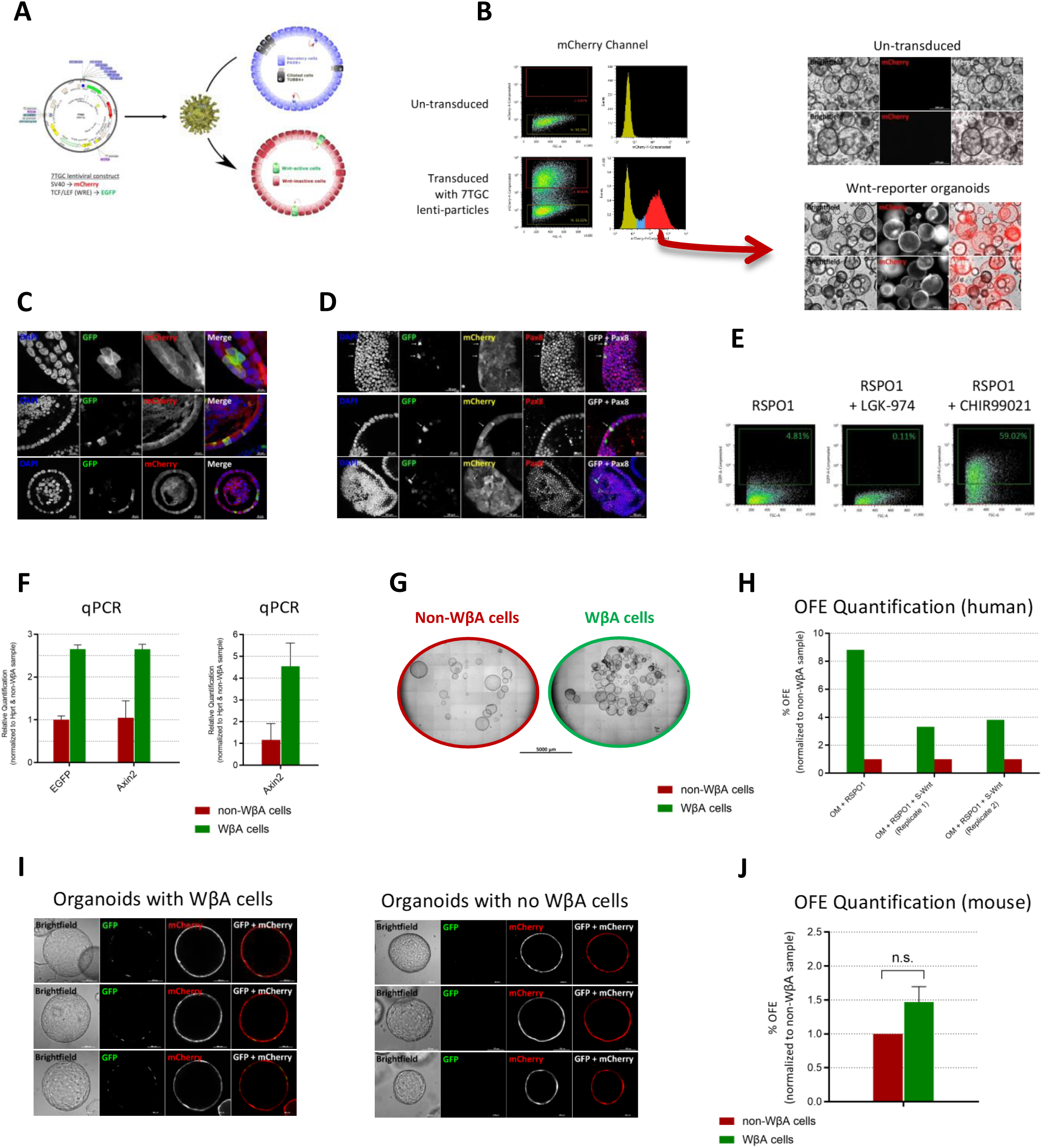
WNT/β-catenin Active (WβA) cells drive organoid regeneration. **(A)** 7TGC lenti-construct used to generate WβS-reporter organoids (see Methods for details). **(B)** Left panel: representative FACS profile of mCherry expression upon viral transduction with 7TGC lenti-particles. Right panel: live cell fluorescence microscopy images of 1-week old organoids established from FACS-selected transduced cells (red arrow). Representative of 4 biological replicates (n=4 patients). Scale bars, 500 μm. **(C-D)** Confocal imaging of fixed, whole-mounted hFT WβS-reporter organoids showing WβA (EGFP+) cells **(C)** and their secretory (PAX8+) lineage shown by white arrows **(D)**. Representative of 4 biological replicates (n = 4 patients). Scale bars as indicated. **(E)** FACS analysis of EGFP expression in mOV WβS-reporter organoids treated for 13 days with RSPO1, GSK-3β inhibitor CHIR99021 or/and Porcupine inhibitor LGK-974, as shown. Representative of 5 biological replicates (two replicates shown later in other contexts: Figures 4B and S4C). **(F)** Representative RT-qPCR analysis of the relative mRNA expression of EGFP and WβS activation marker Axin2 in WβA vs non-WβA cells that were FACS-isolated from WβS-reporter organoids. Error bars represent mean ± 95% confident interval for three technical replicates. Right panel shows a second biological replicate. **(G-H)** hFT WβA cells are enriched in organoid formation ability. **(G)** Representative brightfield whole-well images of organoids formed by WβA or non-WβA cells after FACS isolation and organoid culture for 22 days. 25,000 cells were plated in 50 μl Matrigel drops on 24-well plates. Scale bar, 5000 μm. **(H)** Quantification of OFE enrichment in WβA cells. **(I)** Live cell imaging of mOV WβS-reporter organoids after 1-2 weeks in culture. Images representative of > 100 organoids. Scale bars, 500 μm. **(J)** Quantification of OFE in mOV WβA vs non-WβA cells. Error bars represent mean ± SEM for 5 biological replicates. n.s.; not significant.

FACS-purified WβA cells displayed enhanced OFE relative to non-WβA cells (Figure 2 G-H). Whilst WβA cells were detected in all organoids within hFT organoid cultures, mOV organoids were either positive or negative for WβA cells (Figure 2I & S4F) and mOV WβA cells do not display enhanced OFE (Figure 2J) under a range of experimental conditions (Figure S4 G-I), consistent with WβS-independence seen in mOV organoids (shown earlier, Figure 1 J-K).

### WNT7A is the driver of WβS and FT stem cell renewal

To identify the WNT ligand driving stem cell-mediated expansion of hFT organoids, we isolated WβA (EGFP+) and non-WβA (EGFP-) cells from WβS-reporter organoids and profiled their transcriptomes at the single cell level. We applied the SMART-seq2 protocol (Picelli *et al*., 2014) on a total of 1,021 cells (442 WβA cells; 579 non-WβA cells) from WβS-reporter organoids established from 3 patients. This identified *WNT7A* as the only robustly expressed WNT ligand in hFT organoids (Figure 3A). *WNT7A/Wnt7a* expression was further validated using RNAScope FISH staining in hFT organoids (Figure 3B; controls, Figure S5A), hFT tissue (Figure S5 B-C), mOV organoids (Figure 3C; controls, Figure S5D) and mOV tissue (Figure S5 E-F). We also found that *WNT7A+/Wnt7a*+ cells were not exclusively positive or negative for WβS activation markers *AXIN2* and *LGR5* (Figure 3 B-C), so it remains unclear whether WNT7A signals in an autocrine or paracrine manner. However, our scRNA-seq data implicates WNT7A as the target of Porcupine inhibitors IWP-2 and LGK-974 which abolish organoid regeneration (shown earlier, Figure 1 F-I). In contrast, *WNT3A* expression was not detected in organoids (shown earlier, Figure 3A), suggesting hFT cells are not naturally primed to respond to WNT3A or activate WβS through it. This is consistent with WNT3A’s failure to rescue growth of WNT-blocked organoids (shown earlier, Figure 1 H-I).

**Figure 3:**
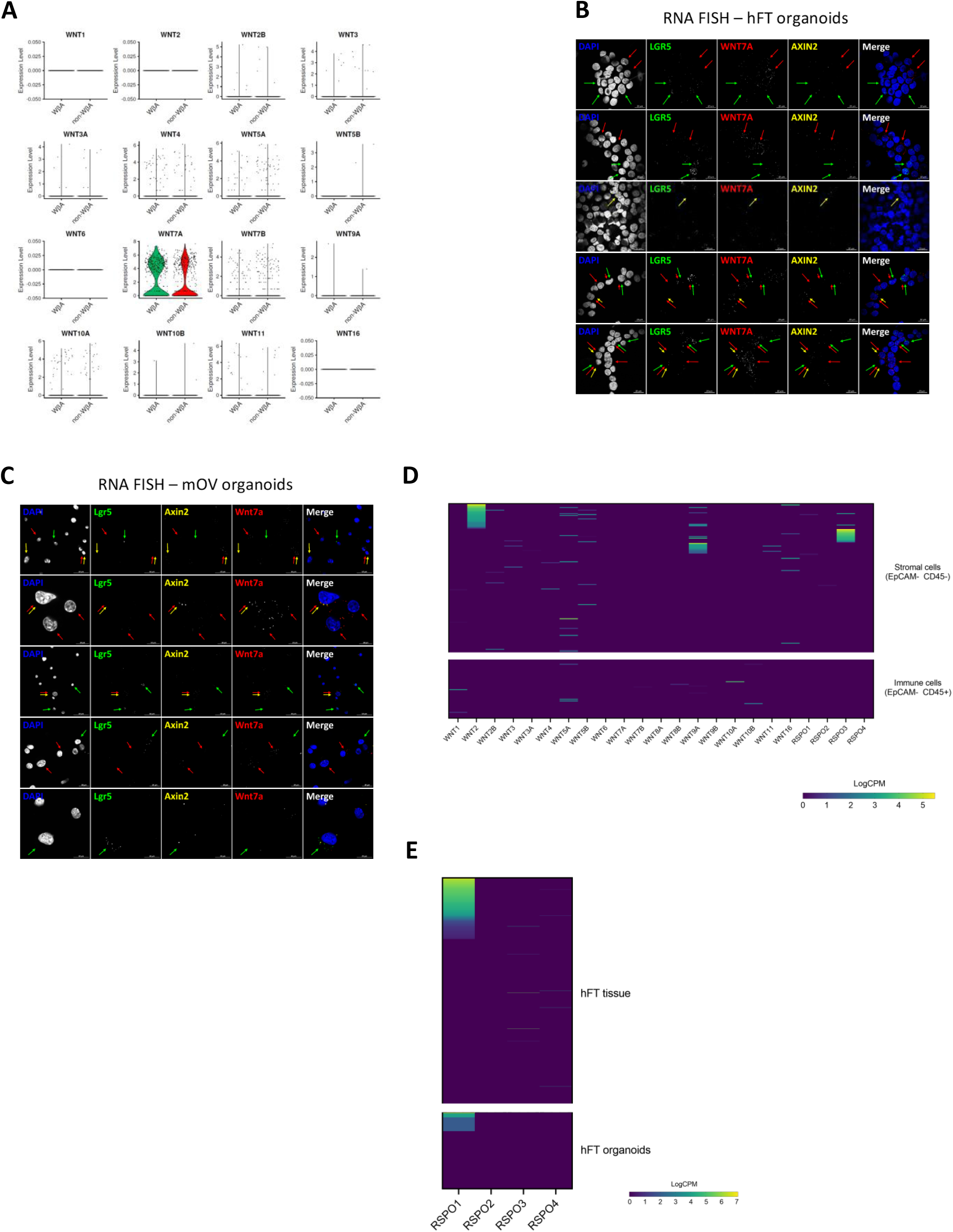
WNT7A is the WNT ligand that cooperates with RSPO1 to drive WβS and organoid regeneration. **(A)** Violin plots showing the single cell expression profile of the WNT family of ligands in WβA (GFP+) and non-WβA (GFP-) cells isolated by FACS from WβS-reporter organoids (n=3 patients). Each dot represents one cell. Based on SCT data, the frequency of WNT7A+ cells is ∼ 5% in hFT tissue and ∼20% in hFT organoids. **(B-C)** Confocal images of RNAScope FISH staining for WNT7A/Wnt7a and WβS activation markers AXIN2/Axin2 and LGR5/Lgr5 in hFT **(B)** and mOV **(C)** organoids. Arrows point to cells expressing colour-matched genes. Arrow clusters indicate cells co-expressing colour-matched genes. LGR5+/Lgr5+ cells and AXIN2+/Axin2+ cells are largely distinct. Organoids were dissociated, cytospinned and fixed prior to RNA FISH staining. Scale bars as indicated. **(D)** Heatmap showing the single cell expression profile of the WNT & R-Spondin family of ligands in the non-epithelial compartments of hFT tissue. Stromal and immune cells were FACS isolated using the antibodies indicated. Each row represents one cell. **(E)** Heatmap showing the single cell expression profile of the R-Spondin family of proteins in hFT tissue (upper panel) and hFT organoids (lower panel). Each row represents one cell.

In addition to epithelial WNT7A, WNT sources from non-epithelial compartments could contribute to WβS activation in epithelial FT cells *in vivo*, as shown for other tissues (Shoshkes-Carmel *et al*., 2018; Zhao *et al*., 2017; Degirmenci *et al*., 2018). To further explore this, we surveyed the expression of the WNT family of ligands in our SCT dataset from stromal and immune cells isolated from patient tissue. This analysis showed that 25% of stromal cells express *WNT2* and 7.5% express *WNT9A* (Figure 3D), suggesting epithelial WNT7A may be redundant for FT renewal *in vivo*. Unlike endogenous WNT7A which is sufficient for organoid regeneration, we found that endogenous RSPO1, the predominant RSPO family ligand expressed in hFT tissue and organoids (Figure 3E), is not sufficient to promote organoid regeneration which relies on exogenously supplied RSPO1 protein (shown earlier, Figures 1 B-C & F-I). We reasoned that another source of RSPOs could be involved *in vivo*, and our non-epithelial SCT dataset showed that 10% of stromal cells robustly express *RSPO3* (Figure 3E) which biochemical studies indicate is over 20-fold more potent in augmenting WβS compared to RSPO1 (Park *et al*., 2018). Notably, the *WNT2+*, *WNT9A+* and *RSPO3+* stromal cells are largely distinct, and we do not see robust WNT or R-Spondin contribution from the limited number of immune cells profiled (Figure 3D).

Next, we sought to functionally confirm whether WNT7A is essential for renewal of FT stem cells in organoids. Direct genetic knockdown of *WNT7A* mRNA was unsuccessful using 10 lentiviral shRNA vectors from two commercial sources (data not shown). We also aimed to test whether WNT7A protein rescues OFE of WNT-blocked organoids. In line with the reported difficulties in generating functional WNTs for *in vitro* assays (Tüysüz *et al*., 2017), all generated WNT7A protein reagents were not functional, including WNT7A CM derived from *WNT7A*-cDNA plasmid-transfected HEK293 cells (Figure S6 A-B), WNT7A CM from primary 2D-cultured FT cells (Figure S6C) which we find endogenously overexpress WNT7A (Figure S6 D-E), native WNT7A protein (Figure S6F) and recombinant WNT7A protein from two commercial sources (one shown in Figure S6G).

Interestingly, the *WNT7A* cDNA expression plasmid robustly activated WβS in the TOPFlash assay, but the CM derived from the same cDNA plasmid-transfected cells failed to activate WβS (Figure S6G) despite containing abundant WNT7A protein (shown earlier, Figure S6 A-B). We reasoned this could be due to a short signaling range, which was shown by biochemical and *in vivo* approaches for certain WNT ligands (Farin *et al*., 2016; Goldstein *et al*., 2006; Alexandre *et al*., 2013). We confirmed this for WNT7A using a simple co-culture assay (Figure S6 H-I) which explained why our protein-based WNT7A reagents were not functional.

Finally, we confirmed the observation above, namely the ability of WNTs to activate WβS as transfected cDNA plasmids but not as secreted proteins derived from the same plasmid, for 4 other canonical WNTs (data not shown). In contrast, we found WNT3A deviates from this pattern, in that it robustly activates WβS as a transfected cDNA plasmid and as a secreted protein derived from the same plasmid as the above WNTs (Figure S6J), implying a unique functional or signaling biology that merits further investigation.

### FZD5 mediates WNT7A-driven maintenance of FT stem cells

To overcome the above limitations and test the functional contribution of WNT7A to organoid renewal, we attempted to identify and biochemically perturb the WNT7A receptor in organoids. SCT profiling identified *FZD3*, *FZD5*, *FZD6* and *FZD10* as the major FZDs expressed in hFT organoids (Figure 4A). Excluding *FZD10*, these were also the major FZDs expressed in hFT tissue (Figure S7A). The FZD family of receptors are categorized into 4 subfamilies based on sequence and structural similarity: FZD5/8 subfamily, FZD4/9/10 subfamily, FZD1/2/7 subfamily and FZD3/6 subfamily. The FZD3/6 subfamily is the most divergent from other FZD family members (Figure S7B) and participates in non-canonical Wnt signaling (Dong *et al*., 2018). FZD5, and not FZDs 3/6/10, was reported to bind WNT7A and activate WβS as shown in *in vitro* WNT-FZD pair screens (Yu *et al*., 2012; Voloshanenko *et al*., 2017) and *in vitro* models (Caricasole *et al*., 2003; Carmon & Loose, 2008). Based on these reports, we ruled out *FZD10* as a receptor that transduces WNT7A-induced WβS in hFT organoids.

**Figure 4:**
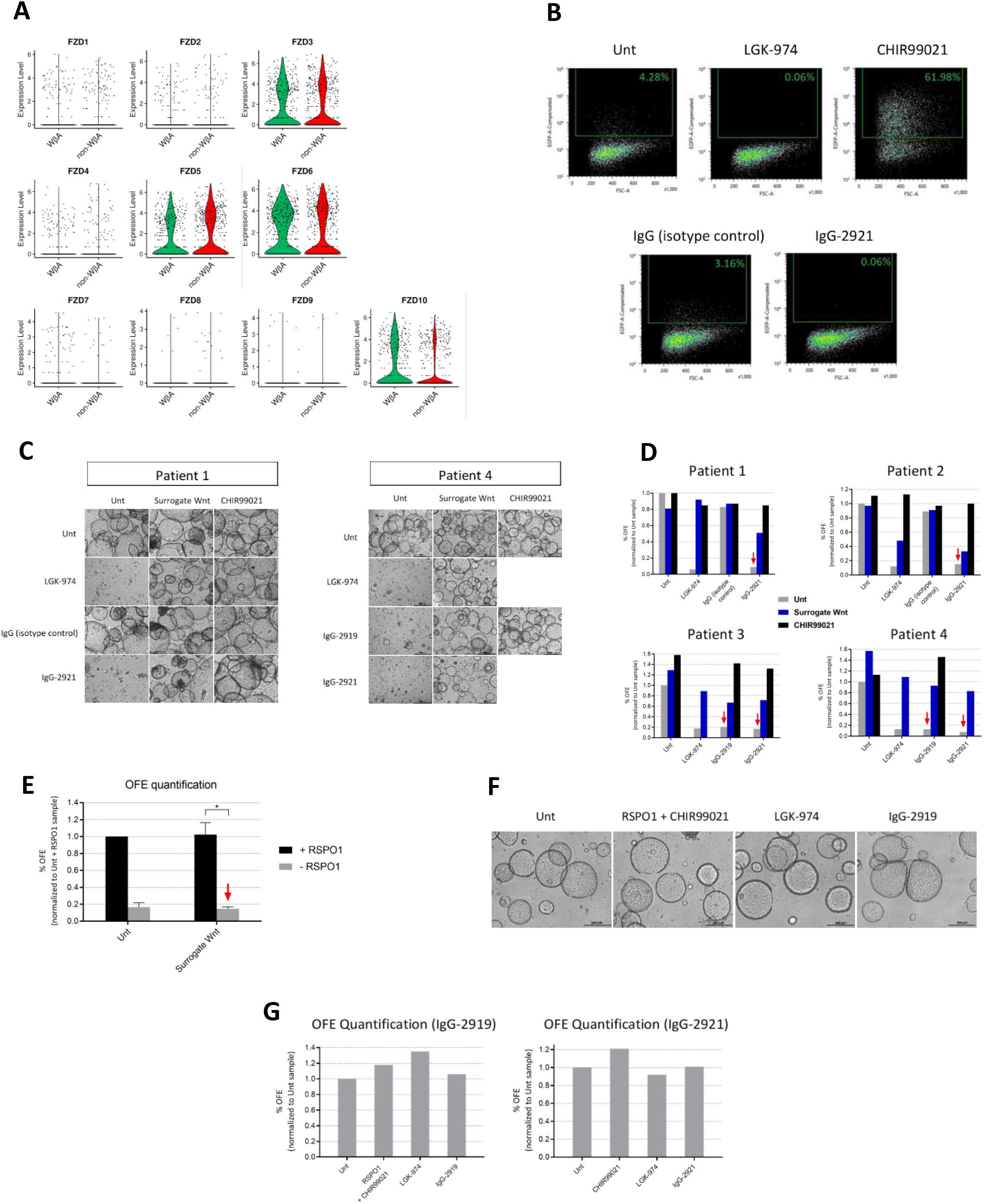
FZD5 is the WNT7A receptor. **(A)** Violin plots showing the single cell expression profile of the Frizzled family of receptors in WβA (GFP+) and non-WβA (GFP-) cells isolated by FACS from WβS-reporter organoids (n=3 patients). Each dot represents one cell. **(B)** FACS analysis of EGFP expression in hFT WβS-reporter organoids upon treatment with anti-FZD5 (IgG-2921) antibody for 7 days. Green gates within plots indicate percentage of EGFP+ cell. LGK-974 or CHIR99021 treatments were used as positive control treatments for abolishing or increasing %EGFP+ (WβA) cells, respectively. **(C-D)** FZD5 mediates WβS-dependent regeneration of hFT organoids. **(C)** Representative brightfield images of p.1 hFT organoids treated for 10-13 days under the conditions shown. Images shown for two biological replicates (n=2 patients). 3-4K cells were plated in 7 μl Matrigel drops. All samples contained RSPO1. Scale bars, 500 μm. **(D)** Quantification of OFE for the samples shown in **(C)** plus 2 additional patient replicates. Red arrows point to substantial OFE reduction by anti-FZD5 antibodies. P-values: Unt vs LGK-974 < 0.0001 (n=4 patients), Unt vs IgG-2921 < 0.0001 (n=4 patients). P-values were calculated using Student’s t-test, two-tailed, paired. **(E)** Quantification of reduction in Surrogate Wnt-driven OFE upon RSPO1 withdrawal (red arrow). Error bars represent mean ± SEM for three biological replicates (n=3 patients). *P = 0.0036 (t-test, two-tailed). **(F-G)** mOV OFE is unaffected by blocking FZD5 receptor. **(F)** Representative brightfield images of mOV organoids treated for 11 days under the conditions shown (in the absence of RSPO1, unless otherwise indicated). 2,000 cells were plated in 50 μl Matrigel drops on 24-well plates. Scale bars, 500 μm. **(G)** Quantification of OFE for the samples shown in (**F**, left panel) as well as for another biological replicate using a different anti-FZD5 IgG (IgG-2921, right panel).

Furthermore, to validate the findings of previous reports, we utilized HEK293 cells which endogenously express FZDs 3/5/6 (Figure S7C). siRNA knockdown (Figure S7D) of *FZD5*, but not *FZD3* or *FZD6*, inhibited *WNT7A*-induced TOPFlash (Figure S7E) while WNT7A induced the highest WβS/TOPFlash signal in a background of FZD5 overexpression compared to FZD3 or FZD6 overexpression (Figure S7F).

Although this data confirms WNT7A can activate WβS through FZD5, direct functional evidence from hFT organoids is lacking. To address this, we utilized IgG-2919 and IgG-2921; two novel, selective, antibody-based inhibitors of FZD5 generated using an antibody phage-display system (Steinhart *et al*., 2017). Anti-FZD5 IgGs reduce TOPFlash signal in WNT7A and FZD5 overexpressing cells by more than 60% (Figure S7G). Consistent with the dual effect of WNT inhibition on abolishing WβA cells (shown earlier, Figure 2E) and OFE (shown earlier, Figure 1 F-I), FZD5 inhibition abolished WβA cells (Figure 4B) and OFE (Figure 4C) by over 90% (Figure 4D). This phenocopies the effect of RSPO1 withdrawal (shown earlier, Figure 1 B-C & 1 F-I). The only other FZD that shows cross-reactivity with anti-FZD5 antibodies is FZD8 (Figures 3C & S5 in Steinhart *et al*., 2017) which is not expressed in hFT organoids (shown earlier, Figure 4A). Furthermore, FZD5-inhibited or WNT-blocked organoids were rescued by WβS activation downstream of ligand-receptor interactions using the selective

GSK-3β inhibitor CHIR99021 (Figure 4 C-D). Partial rescue is also observed when WβS is activated at the ligand-receptor level using Surrogate Wnt (Figure 4 C-D), which competes with anti-FZD5 IgG’s for binding to FZD’s cysteine-rich domain (Janda *et al*., 2017; Steinhart *et al*., 2017). Surrogate Wnt has broad-spectrum activity against FZDs 1/2/5/7/8 but not FZDs 3/6/10 (Janda *et al*., 2017). Since only FZDs 3/5/6/10 are expressed in hFT tissue and organoids, this provides further evidence that FZD5 is the FZD receptor associated with WβS and organoid regeneration. FZD receptors are subject to constant turnover and proteasomal degradation by the action of the RNF43/ZNRF3 ubiquitin ligases (Hao *et al*., 2012), which are inhibited by R-Spondins (Carmon *et al*., 2011; Glinka *et al*., 2011; de Lau *et al*., 2011). As such, withdrawal of RSPO1 from Surrogate Wnt-treated organoids reduces OFE by 75-90% (Figure 4E). Since FZD5 is the only FZD targetable by Surrogate Wnt, this indicates that the reduced OFE seen upon RSPO1 withdrawal from Surrogate Wnt treated (Figure 4E) and untreated (shown earlier; Figures 1 B-C & 1 F-I) organoids is due to FZD5 turnover. Altogether, these data strongly suggest that FZD5 is the cognate receptor for endogenous WNT7A in the FT, and that a WNT7A-FZD5 signaling axis drives WβS activation and renewal of hFT organoids.

While organoids from tested patients in this study showed sensitivity to WβS inhibition, organoids from one patient were resistant to WNT and FZD5 inhibition (Figure S8 A-B) as well as to RSPO1 withdrawal (Figure S8C). Patient 5 organoids were, however, sensitive to downstream WβS inhibition using XAV-939 and PRI-724 (Figure S8 A-B) as seen in other patients (shown earlier, Figure S3G). Tested under selective conditions of WNT-blocking for four passages, Patient 5 organoids show ectopic and robust growth that led to large organoid sizes not typical of normal FT organoids (Figure S8D). Although Patient 5 was diagnosed with serous ovarian cancer, which is thought to derive from the FT (Kuhn *et al*., 2012; Kim *et al*., 2018), Patient 5 organoids do not carry TP53 mutations, which are known to be early events in high grade serous ovarian cancer (Labidi-Galy *et al*., 2017). While further characterization and more patient replicates are required to reproduce this finding, these WNT/FZD5-resistant organoids may represent mutant clones with early genetic changes that confer selective growth advantage and independence from stem cell niche factors.

Finally, FZD5 inhibition has no effect on mOV organoids (Figure 4 F-G) consistent with the lack of effect seen upon blocking WNT secretion (shown earlier, Figure 1 J-K) and lack of OFE enrichment in isolated WβA mOV cells (shown earlier, Figure 2J). This is despite WNT blocking and FZD5 inhibition abolishing WβA cells in mOV WβS-reporter organoids (shown earlier, Figure 4B), suggesting that a WNT-FZD5 signaling axis also regulates WβS in mOV organoids, but unlike in hFT organoids, WβS is not essential for stem cell renewal in the mOV organoids.

### Estrogen downregulates WNT7A and triggers differentiation

Female Reproductive Tract (FRT) organs, including the hFT/mOV, are subject to cyclic hormonal influences. Estrogen acts as a ligand that dimerizes estrogen receptors (ER) expressed in cells within hormone-responsive tissues (Björnström & Sjöberg, 2005). Estrogen’s influences on WβS in the hFT remain to be elucidated, as some studies indicate estrogen activates WβS in the FRT (Hideyuki *et al*., 1999; Kouzmenko *et al*., 2004; Hou *et al*., 2004) while others provide evidence that estrogen exerts an inhibitory influence on WNT7A (McLachlan *et al*., 1980; Couse *et al*., 2001; Wagner & Lehmann, 2006). To address this knowledge gap, we examined the effects of estrogen on WNT7A and WβS in our FT organoid model.

Our SCT data confirms that our optimized organoid culture conditions successfully maintain hormone receptor-expressing cells (Figure 5A) as seen in human tissue (Figure 5B). In both settings, ERα is the predominant hormone receptor expressed, and we harnessed this model to understand estrogen’s influence on the FT. Estrogen treatment of hFT organoids triggers a phenotype of shrivelled, condensed and darker morphology organoids with extensive internal folding and invaginations (Figure 5 C-D), reminiscent of the differentiation morphology that appeared in long-term cultured organoids (shown earlier, Figure 1D) and in differentiated organoids of other tissues (Yin *et al*., 2013). However, estrogen triggered these changes within 72 hrs. On the molecular level, hFT organoids respond robustly to estrogen treatment by upregulating the expression of the canonical estrogen target genes *PGR* and *TFF1* (Figure 5E) and in this setting, we found that estrogen downregulated expression of *WNT7A* and the RSPO1 receptor *LGR6*, as well as the WβS reporter *AXIN2* (Figure 5F). Furthermore, we detected downregulation of the secretory cell marker *PAX8* and upregulation of *FOXJ1* (Figure 5G), an established master regulator and marker of ciliated cells (You *et al*., 2004), as well as upregulation of *CAPS*, *CCDC17* (Figure 5G) and *CCDC78* (Figure 5H); all novel ciliated cell markers we previously identified in a tissue-based SCT study (Hu *et al*., 2020). Progesterone did not antagonize estrogenic molecular changes in this setting (Figure 5 E-G). Therefore, estrogen triggers hFT organoid differentiation towards the ciliated cell lineage.

**Figure 5:**
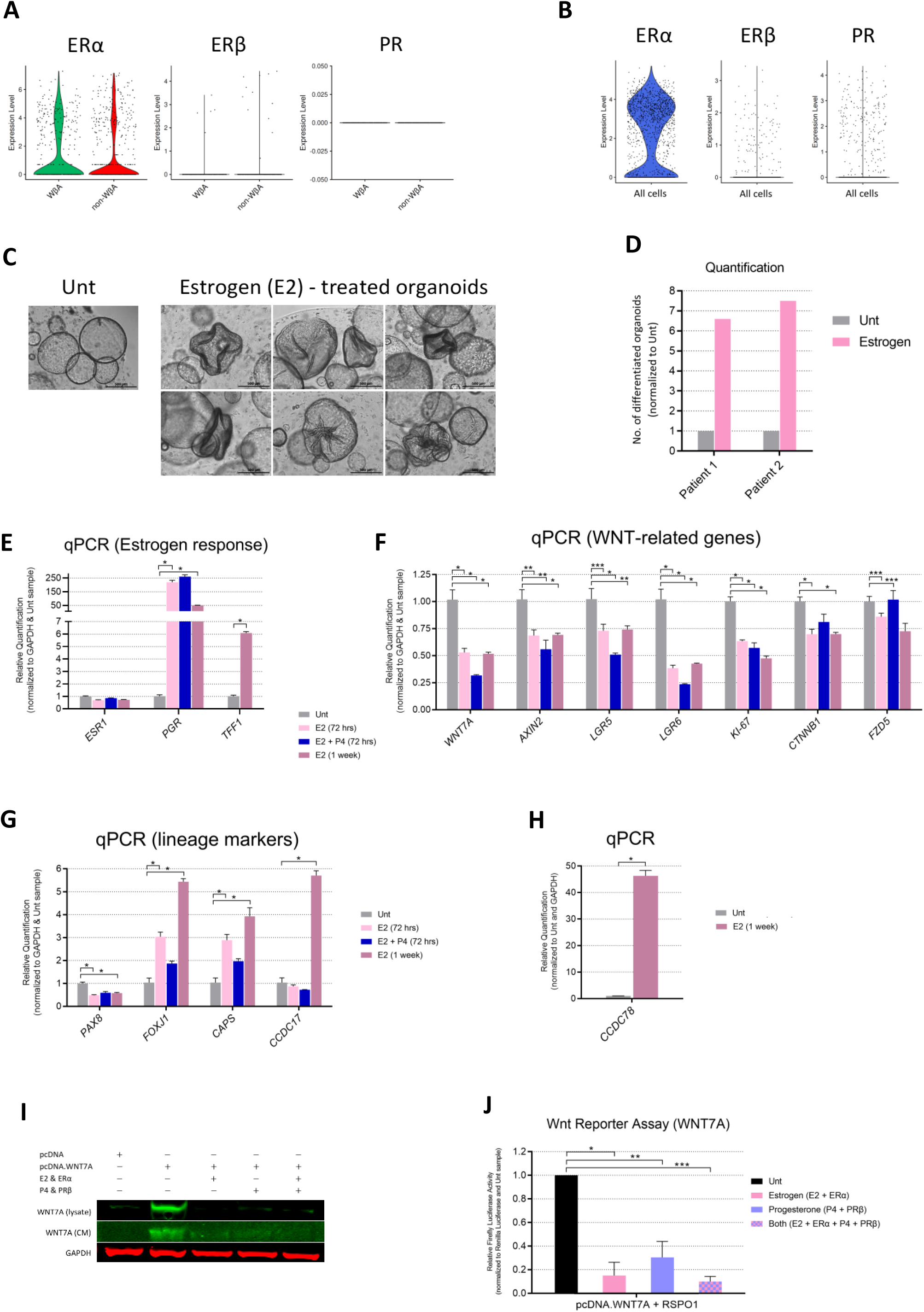
Estrogen suppresses WNT7A and WβS and promotes differentiation of hFT organoids. **(A)** Violin plots showing the single cell expression profile of female hormone receptors in WβA (GFP+) and non-WβA (GFP-) cells isolated by FACS from WβS-reporter organoids (n=3 patients). Each dot represents one cell. **(B)** Violin plots showing the single cell expression profile of female hormone receptors in fresh EpCAM+ CD45-cells isolated by FACS from hFT tissue (Hu et al., 2020) and subjected to SMART-seq2 protocol (Picelli et al., 2014). Each dot represents one cell. **(C-D)** Estrogen trixggers differentiation and morphological changes in hFT organoids. **(C)** Representative brightfield images of differentiated organoids that arise after treatment with estrogen (17β-estradiol, 100 nM) for 72 hrs. Scale bar, 500 μm. **(D)** Quantification of enrichment in differentiated organoids shown in **(C)** plus another replicate. **(E-H)** Estrogen suppresses WNT7A and WβS, and induces a ciliogenesis differentiation programme in hFT organoids. RT-qPCR analysis of the relative mRNA expression of: **(E)** ESR1 and the canonical ER target genes PGR and TFF1; **(F)** WNT7A, AXIN2, LGR6 and other WNT-related genes shown; **(G-H)** secretory and ciliated cell markers. For **(E-H)**, treatments were administered to expanded organoids 7-9 days after organoid passage. Estrogen (17β-estradiol, 100 nM) or progesterone (1 μM) were administered at the indicated concentrations, for 72 hrs (short-term) or 1 week (long-term) as indicated on the panels. Data in **(E-H)** are for the same samples. E2 (1 week) treatment is normalized to Unt (1 week) control sample not shown for simplicity. Error bars represent mean ± SEM 3-6 technical replicates. Asterisks denote statistical significance calculated using unpaired student’s t-test, as follows: **(E)** *P < 0.00008; **(F)** *P < 0.009, ** P < 0.05, *** P = n.s.; **(G)** *P < 0.004; **(H)** *P < 0.0001. Additional replicates (3-12 technical replicates, n = 2 patients) for the effect of estrogen on the indicated genes are shown in Figure S9 A-D in a different context. **(I-J)** Estrogen (& progesterone) downregulate WNT7A protein level **(I)** and activity **(J)**. **(I)** Western blot showing cytoplasmic (lysate) and secreted (conditioned medium, CM) levels of WNT7A protein. Representative of n = 2 biological replicates. **(J)** TOPFlash assay with the indicated treatments. All samples were transfected with the pcDNA.WNT7A construct and treated with RSPO1. Error bars represent mean ± SD for two biological replicates. P-values = *0.009, **0.019, ***0.001 (t-test, two-tailed).

Inhibition of the Notch signaling pathway was reported to promote the ciliated cell lineage in hFT/mOV organoids (Kessler *et al*., 2015; Xie *et al*., 2018). To determine whether estrogen induces ciliogenesis through inhibiting Notch signaling, we treated hFT organoids with the Notch signaling inhibitor DAPT, which reduced the expression of Notch target genes (Figure S9A). In addition to downregulating WNT7A and WβS, estrogen downregulated the expression of Notch target genes to levels comparable with the Notch inhibitor DAPT (Figures S9A). However, unlike estrogen, inhibition of Notch signaling alone did not potently induce ciliogenesis in hFT organoids (Figures S9 C-D) as reported (Kessler *et al*., 2015; Xie *et al*., 2018). We next asked whether estrogen’s influence could be phenocopied by inhibiting both WβS and Notch signaling. To this end, we found that treatment of organoids with small molecule inhibitors of endogenous WβS (LGK-974) and endogenous Notch signaling (DAPT) phenocopied the ciliogenesis-inducing effect triggered by estrogen (Figures S9 C-D), suggesting that estrogen influences cell fate decisions in hFT organoids, at least in part, through downregulating the WNT and Notch pathways.

To further examine the effects of estrogen and progesterone on WβS, we directly examined the influence of these hormones on transcription at the WβS responsive elements (WβS-RE) using the TOPFlash assay in HEK293 cells. Estrogen robustly activated transcription from WβS-RE (Figure S9E), independently of WNT ligands (Figure S9F). Estrogen’s activation of transcription at WβS-RE elements was unaffected by WβS inhibitors at midstream (XAV-939) and downstream (PRI-724) levels (Figure S9F). Estrogen also attenuated ligand-dependent and ligand-independent hyperactivation of WβS (Figure S9G) to levels seen in estrogen-treated samples (shown earlier, Figure S9E), indicating that liganded estrogen-ERα complex may compete with β-catenin for binding and activation at WβS-REs. To test this hypothesis, we investigated the effect of estrogen on TOPFlash signal upon β-catenin knockdown. β-catenin siRNAs were validated to effectively abolish β-catenin protein levels (Figure S9H) and activity (Figure S9 I-J). In this setting, liganded ERα activated transcription at WβS-RE elements independently of β-catenin (Figure S9K) and induced WβS target gene transcription 45-88% higher in the absence of β-catenin, suggesting the pair compete for binding at WβS-RE promoter elements. Next, we sought to delineate the effect of estrogen on WNT7A. Interestingly, estrogen (as well as progesterone) dramatically reduced intracellular and secreted WNT7A protein levels (Figure 5I) and activity (Figure 5J) without independently activating WβS (as shown earlier; Figure S9E). Overall, these data suggest that estrogen can activate WβS target gene expression in ERα-expressing hormone-responsive tissues, but in the presence of WNT7A (exogenously expressed in HEK293 cells and endogenously present in FT organoids), estrogen specifically suppresses WNT7A and WNT7A-induced WβS.

Finally, WβS inhibition was recently shown to downregulate the expression of DNA double-strand break repair genes, including *BRCA1*, *BRCA2*, *RAD51* and the *FANC* gene family, in various tissues via a WβS-MYBL2 signaling axis (Kaur *et al*., 2021; Angers, 2021). These DNA repair genes were shown to be upregulated endogenously in WβA cells of other tissues (Kaur *et al*., 2021). In addition to inhibiting WβS, we noted that estrogen suppresses *MYBL2* as well as *BRCA1* and *BRCA2* expression in hFT organoids (Figure S9L). This is phenocopied by estrogen-independent WNT inhibition (Figure S9L), raising the possibility that estrogen may regulate *MYBL2* and *BRCA1/2* expression, at least in part, through regulating WβS.

Collectively, the data presented above indicate that estrogen suppresses *WNT7A* and WβS, robustly induces a transcriptional ciliogenesis programme and may regulate *BRCA1/2* expression in hFT organoids.

### Transcriptomic characterization of WβA cells

Our data thus far reveal a hormonally-regulated WNT7A-FZD5 signaling axis which activates WβA cells that drive hFT organoid regeneration, identifying WβA cells as candidate FT stem cells. To further characterize these cells, we performed scRNA-seq (SMART-seq2 protocol; Picelli *et al*., 2014) on WβA and non-WβA cells isolated from WβS-reporter organoids established from three patients (Figure S10A). Unsupervised clustering using Uniform Manifold Approximation and Projection (UMAP) showed that WβA cells are enriched in specific clusters of cells that are distinct from non-WβA cells (Figure 6A). To identify the genes responsible for driving this difference, we conducted an intra-patient differential gene expression analysis between WβA and non-WβA cells, identifying expression signatures that are enriched in these cell types (Figure 6B; raw data containing full gene lists is provided in Table S1). Among these we noted a small number of specific or highly differentially expressed genes which could potentially serve as biomarkers for future studies focusing on gene-based identification or isolation of WβA cells (Figure 6C).

**Figure 6:**
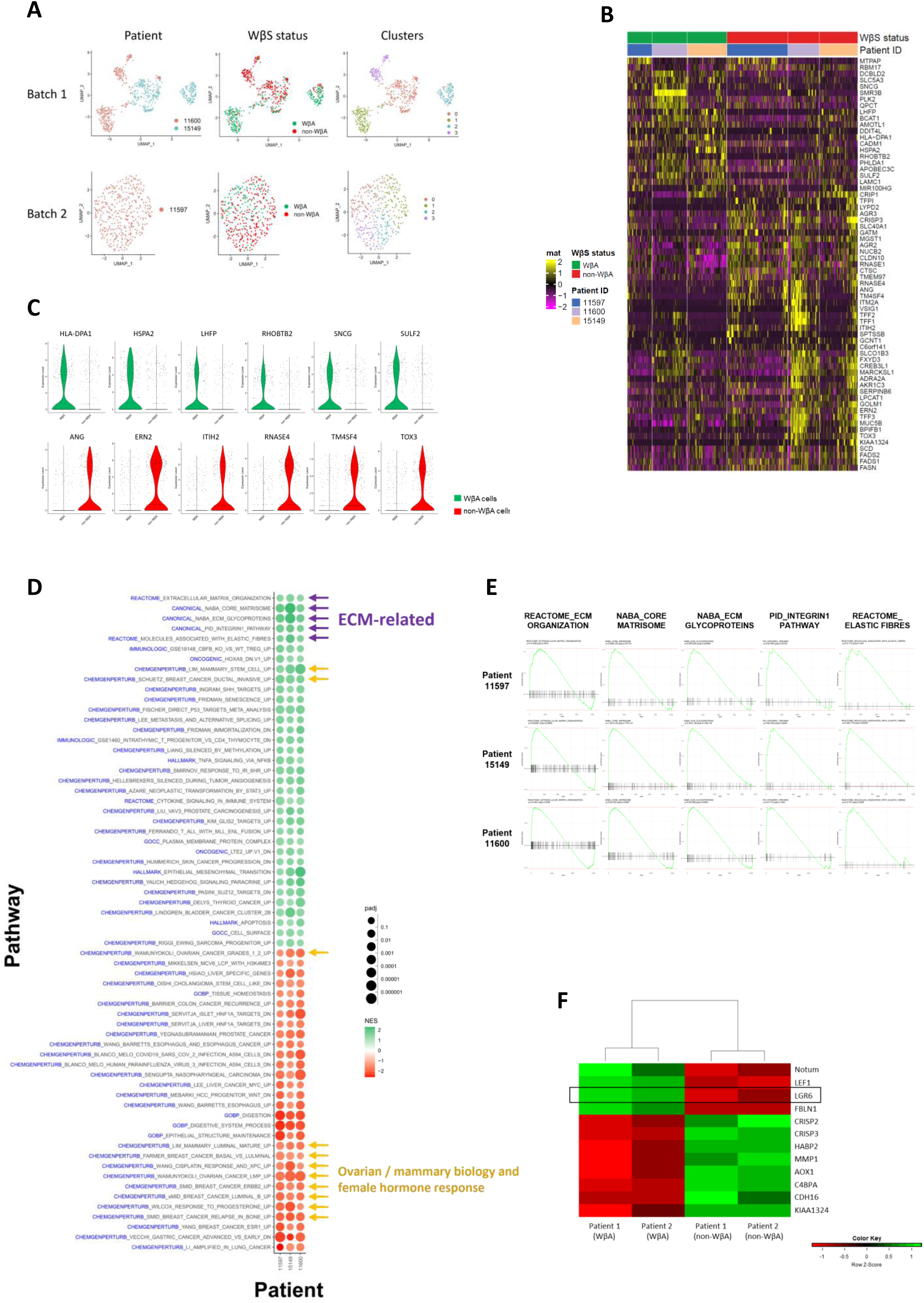
Single cell transcriptomic characterization of WβA cells. **(A)** Uniform Manifold Approximation and Projection (UMAP) showing dimensionality reduction of the single cell transcriptomes of WβA (GFP+) and non-WβA (GFP-) cells. The cells were FACS-isolated from patient-derived hFT WβS-reporter organoids and processed using the SMART-seq2 protocol for scRNA-seq. UMAPs are shown by patient (left panel), WβS status (middle panel) or clusters (right panel). Each batch is shown separately to limit confounding batch effects. **(B)** Heatmap showing patient-shared significant (p-value < 0.05) DEGs derived from intra-patient comparison of the single cell transcriptomes of WβA vs non-WβA cells. Each column represents a single cell. Each row represents a shared DEG. Heatmap colours indicate expression level as shown in the scale bar. **(C)** Violin plots showing the single cell expression profile of specific DEGs that mark WβA cells (green; upper panel) or non-WβA cells (red; lower panel). Each dot represents a cell (n=3 patients). **(D)** Bubble plot showing the results of intra-patient GSEA analysis of pathways that are consistently upregulated (green, +ve NES score) or downregulated (red, -ve NES score) in WβA cells relative to non-WβA cells across the three patients. Each row represents a pathway (blue text indicates gene set collection from which the pathway is derived). Purple arrows point to ECM-related pathways. Orange arrows point to gene sets associated with ovarian & mammary biology and cancer (including female hormone response). GSEA, Gene Set Enrichment Analysis; NES, Normalized Enrichment Score; padj, adjusted p-value; CHEMGENPERTURB, Chemical and Genetic Perturbations; GOBP, Gene Ontology: Biological Processes; GOCC, Gene Ontology: Cellular Component. **(E)** Enrichment plots for the patient-shared, ECM-related pathways shown in **(D)**. GSEA enrichment plots for patient-unique ECM-related processes can be found in Figure S10 E-G. **(F)** Heatmap showing hierarchical clustering of the shared DEGs from the bulk RNA-seq of WβA and non-WβA cells derived from WβS-reporter organoids established from 2 patients. The top DEG’s are shown. Colours indicate the expression levels shown in the scale bar.

To further explore the nature of pathways and processes that characterize WβA cells, we conducted gene set enrichment analyses (GSEA) based on our scRNA-seq data (raw data of the full gene set lists is provided in Table S2). Several WβS-related pathways and processes were enriched in WβA cells (Figure S10 B-D), validating our strategy for capturing WβA cells using WβS-reporter organoids. WβA cells were also enriched in pathways associated with female hormone responsiveness as well as ovarian and mammary tissue biology and cancer (Figure 6D), consistent with the tissue-of-origin of our reporter organoid lines. The most common WβA cell-enriched gene sets, however, are associated with extracellular matrix (ECM) remodelling and integrin signaling in inter-patient (Figure 6 D-E) and intra-patient (Figure S10 E-G) analyses, suggesting a dominant role for ECM processes in maintaining WβA cells.

Finally, we probed the scRNA-seq dataset to identify the RSPO1 receptor responsible for the obligate RSPO1 requirement in hFT organoid regeneration (shown earlier, Figure 1 B-C & F-I). We found that our organoid SCT data does not capture the expression of R-Spondin receptors (LGRs 4-6) possibly due to dropout of low abundance mRNAs. To address this, we performed bulk RNA-seq of WβS-reporter organoids. Similar to the SCT organoid data, the bulk RNA-seq data confirmed enrichment of WβS-related processes and pathways in WβA cells (Table S3) and identified *LGR6* as the only R-Spondin receptor enriched in WβA cells (Figure 6F). This is consistent with our previous functional data, in which organoid differentiation-inducing conditions significantly reduce the expression levels of the WβS reporter gene *AXIN2* concomitantly with reducing *LGR6* expression, with the 6-fold increase in *LGR5* expression unable to rescue this (shown earlier, Figure S9B). Furthermore, we utilized this bulk RNA-seq dataset to probe for the ECM signatures we noted above, and TopFunn pathway analyses identified a strong signature for ECM-related processes in the two patients analyzed, validating our observations in the SCT-derived GSEA analyses that ECM-related pathways are enriched in WβA cells. This suggests that localized ECM remodelling within the physical niche of FT stem/WβA cells in organoids (and potentially *in vivo*) may play a crucial role in maintaining stem cell self-renewal and multipotency as reported for other tissues (Guilak *et al*., 2009; Gattazzo *et al*., 2014; Watt & Huck, 2013) and as elaborated in the discussion below.

## DISCUSSION

The first organotypic culture techniques for human fallopian tube (hFT) and mouse oviduct (mOV) tissues were described relatively recently (Xie *et al*., 2018; Kessler *et al*., 2015), and a number of studies have utilized these culture methods to make important progress in understanding FT and HGSOC biology (Zhang *et al*., 2019; Lõhmussaar *et al*., 2020; Kessler *et al*., 2019; Kopper *et al*., 2019; Hoffmann *et al*., 2020; Hill *et al*., 2018; de Witte *et al*., 2020). While organoids containing distinct FT cell lineages emerge from single multipotent cells in bulk and single cell culture, pointing to stem cell activity, no studies have thus far harnessed hFT organoids for expanding and characterizing stem cells. This is due to limitations in published protocols in which culture conditions are not amenable to single cell isolation and culture. In this report, we optimize serum-free culture conditions that enable stable stem cell expansion from single cells to yield self-renewing, multi-lineage, hormonally-responsive organoids that recapitulate structural and molecular features of hFT tissue. We harness this tool to conduct functional analyses on self-renewal requirements and devise a strategy for biomarking, isolation and transcriptomic characterization of putative stem cells, generating the first single cell transcriptomic dataset of hFT organoids.

Our work establishes the essentiality of WβS for FT epithelial renewal and delineates the molecular factors that regulate this renewal programme. We find that WβS plays a critical role in FT renewal, as demonstrated by our data showing that multi-lineage organoid regeneration is strongly impaired by either: 1) withdrawal of R-Spondin; 2) blocking of endogenous WNT ligand secretion using porcupine inhibitors; 3) stabilization of AXIN by a tankyrase inhibitor; or 4) blocking nuclear β-catenin activity. Furthermore, FT cells with active WβS (WβA cells) that were isolated from our WβS-reporter organoids are enriched in organoid forming capacity, and SCT analysis shows that these cells are transcriptionally distinct and enriched in defined clusters. In addition, our work implicates WNT7A as the endogenous ligand driving WβS activation and organoid generation based on the following evidence: 1) our SCT analysis identified *WNT7A* as the only expressed canonical WNT ligand in hFT organoids; 2) the analysis identified *FZD5* as the mediator of WNT7A-induced WβS signaling; 3) functional perturbation of the WNT7A receptor using anti-FZD5 antibodies strongly impaired organoid regeneration.

Furthermore, our bulk RNA-seq analysis identified *LGR6* (and not *FZD5* or *LGR5*) as enriched in WβA cells, suggesting that LGR6 is one of the limiting factors dictating multipotency of *FZD5*+ *LGR6*+ WβA cells. Finally, our data from hormonally-treated organoids and hormone receptor over-expressing HEK293 cells suggest that estrogen signaling suppresses WβS by downregulating *WNT7A* and *LGR6* mRNA levels, as well as by downregulating WNT7A protein levels.

### WNT7A and FT/oviduct development

Our work implicates WNT7A in FT homeostasis and maintenance. This is consistent with evidence from *Wnt7a* knockout (KO) mice that contain no oviducts or severely compromised oviducts with diminished invaginations, as well as global abnormalities in the correct patterning of the neonatal female reproductive tract or FRT (Miller & Sassoon, 1998). Although viable, *Wnt7a* KO mice are reproductively sterile and the Mullerian duct fails to regress in male mice (Parr & McMahon, 1998). WNT7A is known to play a critical role in patterning and differentiation of other tissue types, including limb, cardiac and neuronal synapses (Bond *et al*., 2003; Ingaramo *et al*., 2016; Hwang *et al*., 2004; Hall *et al*., 2000).

### Mesenchymal WNTs and RSPOs

Although our data point to epithelial-derived WNT7A as sufficient to drive epithelial regeneration of hFT organoids (and possibly hFT tissue), our tissue SCT data indicates that mesenchymal-derived WNT2 could also contribute to WβS activation in FT cells, offering an explanation as to why conditional *Wnt7a* loss does not disrupt homoeostasis of adult mOV tissue *in vivo* (Dunlap *et al*., 2011). As FZDs 3/6/10 are largely non-canonical Wnt signaling mediators (Dong *et al*., 2018), we propose mesenchymal-derived WNT2 also signals through FZD5, consistent with *in vitro* WNT-FZD pair screens showing that WNT2 can activate WβS through FZD5 but not FZDs 3/6/10 (Yu *et al*., 2012), as well as *in vitro* studies in cell lines (Miller *et al*., 2012) and primary cultures obtained from *Fzd5* ^+/-^ or *Fzd5* ^-/-^ mouse models (Lu *et al*., 2013).

Unlike epithelial-derived WNT7A, endogenous epithelial-derived RSPO1 is not sufficient for FT regeneration *in vitro* (and possibly *in vivo*), and our patient stroma SCT data hints at the possibility that the FT epithelium relies on the mesenchyme as an essential source of RSPO3 necessary for epithelial homeostasis. We propose that future *in vivo* gene perturbation studies should account for WNT2 and RSPO3 as mesenchymal-derived WβS factors that may confound epithelial-specific gene knockdown or KO approaches.

### FT stem cells and their niche

This report provides a description of the molecular factors associated with the FT stem cell niche (Figure 7A). In this working model, the renewal programme of FT stem cells is regulated by WβS that is driven by epithelial-derived WNT7A and mesenchymal-derived WNT2, both signaling through the FZD5 receptor. R-Spondins are essential for sustaining WNT7A/2-induced WβS through epithelial-derived RSPO1 and possibly mesenchymal-derived RSPO3, both signaling through the LGR6 receptor. Inhibition of TGF-β signaling is critical for maintaining this WβS-driven renewal programme, and endogenous or secreted inhibitors of this pathway likely emanate from juxtaposed epithelial or mesenchymal cells within the niche.

**Figure 7:**
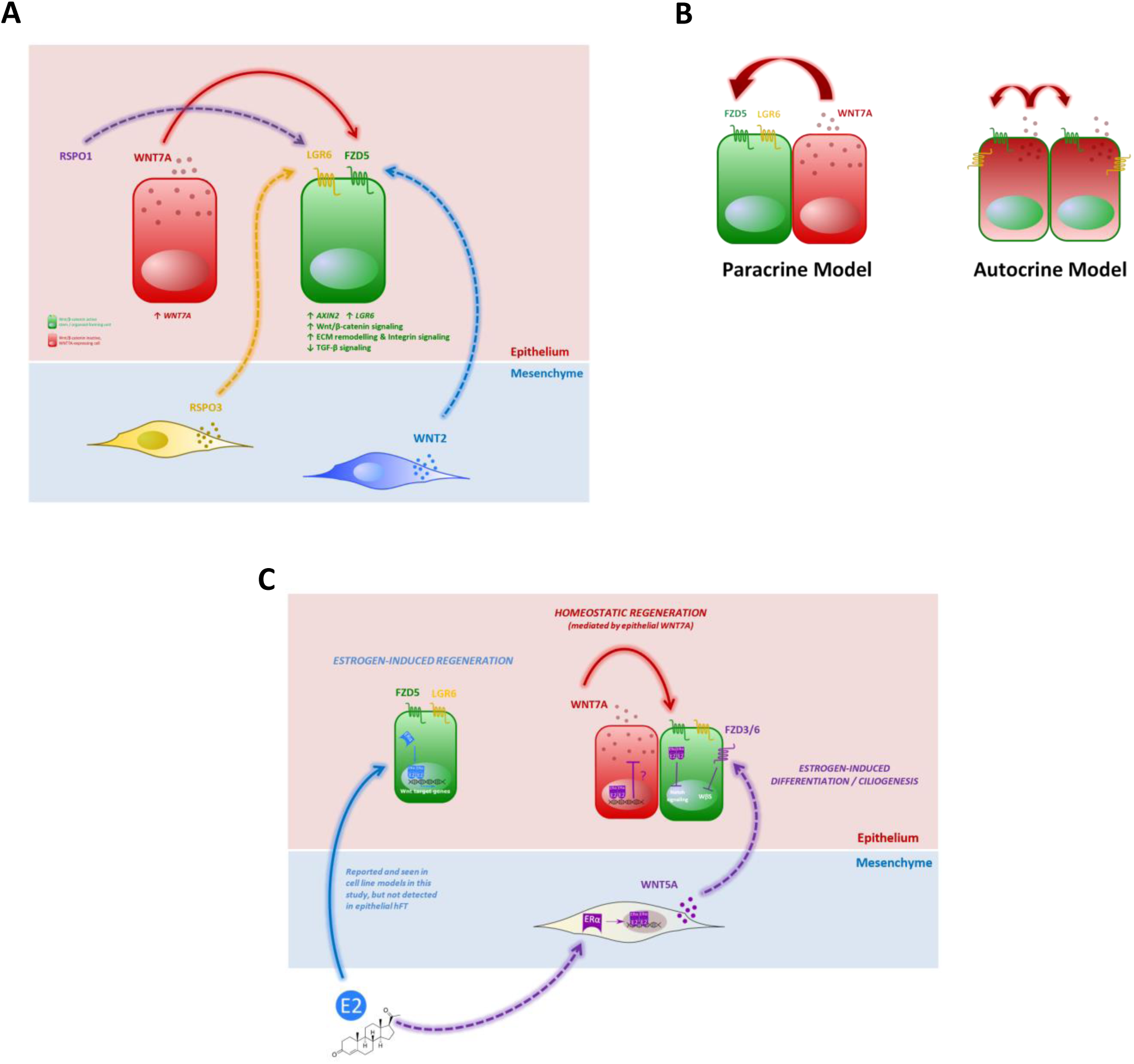
Working model on the molecular maintenance (A), signaling mode (B) and hormonal regulation (C) of FT stem cells. Dashed lines represent hypothesized interactions not shown in this study. WβA cells are depicted in green; non-WβA cells are depicted in red. In (B), the autocrine model proposes that these cell states may co-exist in the same cell.

Our SCT and RNA FISH data suggest WNT7A is also expressed in WβA cells, so it is possible that WNT7A signals in an autocrine manner (Figure 7B). Further biochemical work is required to conclusively establish the signaling mechanism of WNT7A in the FT.

Our work also provides a comprehensive characterization of WβA / FT stem cells. In particular, our gene set enrichment analyses from bulk and single cell transcriptomic assays point to a dominant role played by extracellular matrix (ECM) processes in maintaining WβA / FT stem cells. This is consistent with several *in vitro* and *in vivo* studies which showed that the ECM composition (Battista *et al*., 2005), stiffness (Urciuolo *et al*., 2013), topography (Janson & Putnam, 2015) and the mechanical forces controlled by matrix adhesion (Guilak *et al*., 2009) can regulate stem cell renewal and differentiation across various tissues. ECM-driven influences are thought to be mediated by intracellular mechano-sensing and transduction pathways involving a network of structural support proteins such as integrins, focal adhesions and the actin cytoskeletal network which are hypothesized to transduce changes to the nuclear envelope and influence the activity of transcription factors and chromatin remodelling enzymes (Lim *et al*., 2022; Gattazzo *et al*., 2014; Watt & Huck, 2013). Based on these and other studies, it can be hypothesized that FT stem cell maintenance is driven, at least in part, by cellular programmes that remodel the underlying ECM.

Indeed, WNT7A, which we identify as the molecular signal necessary for maintaining WβA cells that are enriched in ECM processes, synergizes with increased ECM stiffness to promote renewal of muscle satellite stem cells during skeletal muscle regeneration (Moyle *et al*., 2020). In this context, WNT7A synergizes with fibronectin which binds the SDC4 receptor to enable WNT7A-induced self-renewal of satellite stem cells (Bentzinger *et al*., 2013). Another example of the crosstalk between WNT7A and the ECM is seen in the neuroepithelium, where the ECM receptor integrin-β1 was shown to synergize with WNT7A and the ECM protein Decorin in regulating the expansion of neuroepithelial progenitors (Long *et al*., 2016). In a pathological context, tumour-secreted WNT7A was shown to remodel the underlying ECM and enable tumour invasion and metastasis in mouse models of breast cancer (Avgustinova *et al*., 2016). Of note, WNT7A protein expression drives tumour progression and invasion, and clinically correlates with poor prognosis in breast (Avgustinova *et al*., 2016), endometrial (Liu *et al*., 2013), ovarian (Yoshioka *et al*., 2012; King *et al*., 2015; Zhang *et al*., 2010), tongue squamous cell (Jia *et al*., 2019) and bladder (X. Huang *et al*., 2018) cancers. In these settings, WNT7A remodels the ECM by recruiting and activating TGF-β signaling in cancer-associated fibroblasts to induce ECM remodelling (Avgustinova *et al*., 2016) or by inducing the epithelial secretion of ECM remodelling enzymes such as MMP7 (Yoshioka *et al*., 2012) or MMPs 1 and 10 (Huang *et al*., 2018).

Based on these and other studies, it is tempting to speculate that WNT7A’s maintenance of FT stem cells is a cell-extrinsic process, in which WNT7A could maintain stem cell potency not only through cell-intrinsic WβS activation, but also by synergizing with or activating ECM modulating programmes within WβA cells, which lead to ECM remodelling in the extracellular space that, in turn, lead to ECM-to-WβA cell signaling that could contribute to stem cell maintenance. This notion is supported by our observation in patient-derived 2D-cultured FT cells, which lack an exogenously-supplemented ECM and fail to generate multi-lineage FT organoids when transferred onto an ECM/Matrigel scaffold (data not shown) despite endogenously overexpressing *WNT7A* by over 20-fold compared to fresh cells (Figure S6 C-E). However, functional studies are required to elucidate the ECM’s molecular influences on the maintenance of FT stem cells as well as the molecular basis of WNT7A’s remodelling of the ECM. Both are of high clinical relevance: the former could allow engineering of artificial and defined matrices that enable mechanical reprogramming or enriching for patient-derived FT stem cells for cell-based therapies and regenerative medicine. The latter enables the interrogation of the unconventional role of WNT7A in ECM remodelling, which could be the underlying theme in WNT7A-driven tumour invasion and poor prognosis in the above-mentioned cancers.

### WβA cells and mouse models

WβS is recognized as a major driver of adult stem cell (ASC) renewal and tissue regeneration under homeostatic and injury conditions (Clevers, 2006). Amongst various WβS reporters, AXIN2 and LGR5 are the most faithful (Boonekamp *et al*., 2021) and as such, AXIN2 and LGR5-based mouse models have been employed in biomarking and lineage tracing of WβA stem cells in various tissues (Clevers, Loh & Nusse, 2014; Van Camp *et al*., 2014). To this end, *Lgr5*+ cells were shown to contribute to mOV development, but *Lgr5* expression disappears by day 7 after birth (Seishima *et al*., 2019) and lineage tracing studies in adult mice have shown that *Lgr5*+ cells do not contribute to mOV homeostasis (Ng *et al*., 2014). This is consistent with our bulk transcriptomic analysis from WβS-reporter organoids, showing that *LGR5* expression is not enriched in WβA cells, which are instead marked by *AXIN2* and *LGR6*. This is also consistent with our functional work on organoid differentiation, which showed concordance between *LGR6* (not *LGR5*) and *AXIN2* levels, as well as with our RNA FISH data in human and mouse organoids, suggesting that *LGR5*+/*Lgr5*+ cells and *AXIN2*+/*Axin2*+ cells are largely distinct.

Preferential marking of WβA-ASCs by *Axin2* instead of *Lgr5* has been reporter in various tissues. For example, *Lgr5* specifically marks homeostatic WβA-ASCs in various tissues including the intestines (Barker *et al*., 2007; Sato *et al*., 2011) and the pyloric stomach (Barker *et al*., 2010); in the latter *Axin2*+ *Lgr5*-cells are facultative ASCs mobilized upon injury (Sigal *et al*., 2017). However, the opposite trend is seen in other tissue systems such as the liver (Wang *et al*., 2015; Huch *et al*., 2013), stomach (corpus) region (Stange *et al*., 2013; Leushacke *et al*., 2017) and adult vagina (Ali *et al*., 2020; Ng *et al*., 2014), in which *Axin2* is a better marker of homeostatic WβA-ASCs. This was also shown in the endometrium, which shares embryonic origins and a developmental continuum with the oviduct. Here, *Lgr5*+ cells contribute to development but not adult tissue homeostasis (Seishima *et al*., 2019) as seen in the oviduct, and *in vivo* lineage tracing has demonstrated *Axin2*+ cells to be long-lived multipotent stem cells that replenish glandular and luminal epithelia in the adult endometrium (Syed *et al*., 2020). Based on our data, we propose *Axin2* and *Lgr6* as strong candidates for marking putative multipotent FT/oviduct progenitors in future lineage tracing studies. Alternatively, our SCT-derived panel of highly differentially expressed genes (Figure 6C) provide a basis for further validation of these FT WβA-specific markers and their use in gene expression-based lineage tracing approaches.

### Human fallopian tube versus mouse oviduct regeneration

While our work underscores the essentiality of WβS for regeneration of hFT organoids, other reports have shown that regeneration of mOV organoids is WβS-independent (Lõhmussaar *et al*., 2020; Xie *et al*., 2018). We extend this observation to show that: 1) mOV organoid regeneration is unaffected by blocking endogenous WNTs using porcupine inhibitors; 2) mOV cells with active WβS, isolated from our WβS-reporter organoids, do not display enhanced organoid forming capacity under a range of tested conditions; 3) mOV organoid regeneration is unaffected by anti-FZD5 antibodies, which abolish the proportion of cells with active WβS when applied on mOV WβS-reporter organoids. These observations, coupled with our RNA FISH data confirming the presence of *Wnt7a+* cells in mOV tissue and organoids, suggests that a WNT7A-FZD5 signaling axis also regulates WβS in mOV organoids, but unlike in the human setting, this signaling axis is not essential for renewal of mOV stem cells *in vitro*. Similarly, TGF-β signaling inhibition, which is essential for hFT organoid renewal, is dispensable for mOV organoid regeneration. We find these differences to be striking and warrant further investigation, particularly to address whether these deviations represent a biological inter-species difference or are the result of *in vitro* culture conditions.

### Hormonal regulation of the FT stem cell niche

We report novel findings which suggest that estrogen-ERα circumvents the cellular WβS pathway to activate transcription of WβS target genes. However, in models where WNT7A is expressed (hFT organoids and WNT7A-overexpressing HEK293 cells), we find that estrogen suppresses WNT7A protein and WNT7A-induced WβS, and in hFT organoids estrogen downregulates *WNT7A* at the mRNA level, leading to efficient ciliogenesis. Although ciliated cell differentiation was also associated with Notch inhibition, the effect of Notch inhibition alone in triggering ciliogenesis is relatively modest (Xie *et al*., 2018; Kessler *et al*., 2015) and we confirm this in our work. By characterizing estrogen-treated organoids, we report that estrogen potently induces ciliogenesis by coupling WNT inhibition with Notch signaling inhibition, and that inhibition of these pathways independently of estrogen phenocopies this effect. Our work therefore outlines the precise *in vitro* expansion and differentiation conditions to establish long-term patient-derived ciliated cell models that can serve as useful tools for the study of ciliogenesis and epithelial cilia motility (Enuka *et al*., 2012), its hormonal regulation (Li *et al*., 2010; Bylander *et al*., 2010) and the mechanisms of ciliary dysfunction due to pathogenic infections (Lyons *et al*., 2006) or inherited disorders such primary ciliary dyskinesia (Raidt *et al*., 2015).

The estrogenic inhibitory effect we see on WNT7A is consistent with reports showing that W*NT7A/Wnt7a* expression drops in the endometrium during the estrogenic phase of the menstrual cycle in humans (Bui *et al*., 1997) and mice (Miller *et al*., 1998). Our data is also consistent with *in vivo* studies in which treatment of mice with Diethylstilboestrol (DES; an estrogen analogue) phenocopies the above-described *Wnt7a* KO phenotypes (McLachlan *et al*., 1980) and downregulates *Wnt7a* mRNA levels in the neonatal FRT (Miller *et al*., 1998). This downregulation is absent in ERα KO mice (Couse *et al*., 2001) further confirming this is an estrogenic effect. Clinically, human patients exposed *in utero* to DES showed similar phenotypes to *Wnt7a* KO mice, including FT abnormalities, lack of tubal coiling, withered Fimbria and infertility (DeCherney *et al*., 1981).

The molecular mechanism of estrogen’s suppression of WNT7A remains unclear. However, *in vivo* studies in the endometrium suggest mesenchymal WNT5A, activated by ER+ fibroblasts, mediates estrogen’s suppression of *Wnt7a*, as seen by DES treatment of mice or during the estrogen-dominated phase of the estrous cycle of untreated mice (Mericskay, Kitajewski, & Sassoon, 2004). As WNT5A is a documented inhibitor of WβS via non-canonical Wnt signaling (He *et al*., 2008; Olson & Gibo, 1998; Topol *et al*., 2003), we propose it exerts its influence through epithelial FZDs 3/6. Although we do not detect ER+ WNT5A+ cells in our patient stroma SCT dataset, it is plausible that aged patients (which constitute the bulk of our patient cohort) do not activate mesenchymal WNT5A expression due to menopausal absence of estrogen, and further investigation of samples from premenopausal hormone-exposed cohorts is required to ascertain this. Overall, we propose a working model for hormonal regulation of the proposed FT stem cell niche (Figure 7C).

An inherent limitation of this model is the notion of modelling estrogen’s complex interactions *in vitro*. Cells are known to elicit distinct responses to estrogen depending on its local concentration, and the cyclic nature and physiological doses of estrogen *in vivo* are difficult to recapitulate *in vitro*. Therefore, the phenotypes observed in estrogen-supplemented organoids, while providing a preliminary understanding of estrogenic molecular changes, are likely not to capture the full spectrum of molecular and phenotypic changes induced by estrogen *in vivo*. Another limitation of our work is the absence of the stroma in our organoid model, precluding the study of estrogenic stromal changes that regulate mesenchymal-to-epithelial signaling. To address this, we propose the optimization of complex FT organoid co-cultures methods incorporating stromal and other microenvironmental components to dissect these changes as well as study the effect of mesenchymal-derived factors such as WNT2, RSPO3 and TGF-β inhibitors, and their molecular influence on FT stem cells.

### WβA cells & HGSOC

Finally, and as abovementioned, our work defines the *in vitro* expansion and differentiation conditions to establish long-term patient-derived ciliated cell models. These will be valuable tools in interrogating HGSOC, predisposition to which has been associated with ciliated cell genes in genetic linkage studies (Coan *et al*., 2018). It is hypothesized that, although ciliated cells are not the cells-of-origin of HGSOC, disruption of ciliated cell differentiation or ciliary beating function impairs the flow of fluid in the FT lumen. This reduces the clearance of ovulation-derived genotoxic follicular fluid that is rich in inflammatory and oxidative stress factors that can trigger DNA damage and HGSOC initiation (Bahar-Shany *et al*., 2014; King *et al*., 2011; Huang *et al*., 2015). Hence, disruption of ciliated cell differentiation or ciliary beating function can create an environment permissive of HGSOC initiation (Coan *et al*., 2018). In this context, our work suggests that disruption of ciliated cell differentiation can be caused by absence of estrogen in aged postmenopausal patients. This oncogenic disruption in ciliogenesis may also manifest by germline *BRCA* mutation in younger premenopausal patients whose FTs contain a 50% reduction, on average, in the number of ciliated cells (Li *et al*., 2013), or by mutant TP53 which was shown to prevent ciliated cell differentiation in the FT (George *et al*., 2015) as well as in the airway epithelium (McConnell *et al*., 2016). Overall, our differentiation culture conditions are well suited to evaluate these hypotheses.

## Conclusion

In summary, we provide a deep characterization of FT stem cells and the molecular determinants of their renewal and cell fate specification. Our work lays the foundation for subsequent functional and *in vivo* studies on FT homeostasis, disease, hormonal regulation and epithelial-mesenchymal crosstalk. Our findings provide a basis for mechanistic work investigating the role of FT stem cells in ovarian cancer initiation.

## MATERIALS & METHODS

### Human FT clinical samples

Fallopian tube tissue samples were obtained from patients undergoing cancer surgery at the Department of Gynaecological Oncology, Churchill Hospital, Oxford University Hospitals, United Kingdom. Patients were appropriately informed and consented, and cases were recruited as part of the Gynaecological Oncology Targeted Therapy Study 01 (GO-Target-01, Research Ethics Approval #11-SC-0014, Berkshire NRES Committee), as well as under the Oxford Centre for Histopathology Research (OCHRe) / Oxford Radcliffe Biobank (ORB) research tissue bank ethics reference 19/SC/0173.

### Mouse Tissue samples

All mouse tissue samples were harvested and provided by the Biomedical Sciences Facility, University of Oxford. Mouse colonies were maintained in certified and licensed animal facilities and in accordance with the United Kingdom’s Home Office Animals (Scientific Procedures) Act 1986. All personnel handling animals hold Home Office-issued Personal Licenses. Tissues were obtained from female mice, strain C57BL/6 and aged 7-12 weeks.

### Tissue Dissociation & Primary Culture

Primary tissue was washed, cut longitudinally to expose the epithelium and dissociated in pre-warmed Digestion medium containing 2 mg/ml Trypsin (Sigma), 0.5 mg/ml DNase I (Sigma) and 100 U/ml Collagenase Type I (Invitrogen) for 45 min to 1 hour at 37 ^0^C in constant rotation. Cells were passed through a 70, 100 or 250 μm cell strainer and pelleted by centrifugation at 300 g / 5 min / 4 ^0^C, washed with cold DPBS and used for downstream analysis. For primary 2D culture, isolated cells were resuspended in BM2 culture medium containing Advanced DMEM/F12 (ThermoFisher), 12 mM HEPES (ThermoFisher), 5% FBS (GIBCO), 1% Penicillin/Streptomycin (GIBCO), 100 ng/ml EGF (ThermoFisher) and 10 μM Y-27632 (ROCK inhibitor, Sigma), as described (Kessler *et al*., 2015). For Western blot, primary FT 2D culture was expanded in 6-well plates and treated as described. For siRNA knockdown, 40 nM of siRNA was transfected for 72-96 hrs using Lipofectamine 3000 (ThermoFisher Scientific) according to the manufacturer’s protocol.

### Human FT Organoid Culture

To establish human FT organoids, processed cell pellets from dissociated FT tissue were resuspended in Extracellular Matrix (ECM, Matrigel, Corning), plated as 50 μl drops on pre-warmed culture plates and incubated at 37 ^0^C for 20-30 min to allow Matrigel polymerization. Cells were then overlaid with pre-warmed Organoid Medium containing BM2 medium above (without FBS), supplemented with Noggin (100 ng/ml), Fibroblast Growth Factor-10 (100 ng/ml), N2 supplement (1%), B27 supplement (2%), Nicotinamide (1 mM), N-acetyl L-cysteine (1 mM), A83-01 (5 μM), Forskolin (10 μM) and R-Spondin 1 (500 ng/ml) at the indicated concentrations (refers to key resources table for material sources). Surrogate Wnt (0.5 nM, ImmunoPrecise) was used wherever indicated. Y-27632 (10 μM) was added for the first 2-3 days of organoid culture and removed in the first medium change, until the next passage. WNT3A conditioned medium was generated using L-WNT3A cells (ATCC) or in-house by transient transfecting of pcDNA.WNT3A into HEK293 cells (see below).

For maintenance of hFT organoids, organoids were passaged at a ratio of 1:3 to 1:5 every 10-14 days. Briefly, organoids were released from Matrigel by incubation with Organoid Harvesting Solution (Culturex) at 4 ^0^C in rotation, for 45-90 min. Organoids were then collected into a 15 mL or 50 mL falcon tube, pelleted by centrifugation at 300 g / 10 min / 4 ^0^C, washed with cold PBS once and used for passage or flow/FACS-related downstream analysis. For passage, dissociation was performed by mechanically shearing organoids using a p200 pipette. For flow analysis or FACS sorting, single cell dissociation was performed by resuspending organoids in pre-warmed 7.5X TrypLE Express (ThermoFisher) diluted in Organoid Wash Buffer (OWB), for 5-10 min. OWB buffer is composed of complete organoid medium + Y26632 but lacking growth factors (EGF, FGF-10, Noggin and RSPO1). Organoids (now single cells) were then washed twice with cold PBS and utilized for FACS or single cell culture. For cryopreservation, pelleted organoids were mechanically fragmented, embedded in Recovery Cell Culture Freezing Medium (ThermoFisher), transferred to -80 °C freezer overnight and finally transferred to Liquid Nitrogen for long-term storage. For culture re-establishment after cryopreservation, thawed organoids were resuspended in 9 ml OWB buffer, pelleted, washed and cultured in a well of a 24-well plate, as described.

### Mouse Oviduct Organoid Culture and passage

Mouse oviduct organoids were established from primary tissue as described above for hFT organoids. Culture medium for mouse organoids was the same as hFT organoids, excluding A83.01 and Forskolin. Mouse organoids were passaged as reported (Xie *et al*., 2018).

### Organoid Formation Efficiency assay

OFE analysis was performed on single cell dissociated tissue or organoids, derived as described above. After quantifying cell number, cells were resuspended in Matrigel and plated as Matrigel 50 μl drops in 24-well plates or 7 μl drops in 48 or 96-well plates. Unless otherwise stated, treatments were started on plating day for 7-12 days. OFE was estimated as number of organoids emerging from the total number of cultured cells per well. For experiments requiring FACS isolation of FT cells, cells were prepared for FACS by cold PBS washing followed by blocking non-specific antibody staining using FcR Blocking Reagent (Miltenyi) for 10 min in the dark in a fridge. Further antibody incubation was performed using CD45-FTIC and EpCAM-APC (Biolegend), in a total volume of 100 μl volume. WβS-reporter organoids (see below) were FACS isolated using mCherry gating, and non-transduced parental organoid lines were used as negative controls. FACS-isolated cells were sorted directly into Matrigel, cultured for 10-14 days and OFE quantified as described.

### Generation of Wnt/β-catenin signaling reporter organoids

To generate WβS-reporter organoids, viral lenti-particles were generated by transfecting HEK293 cells in a T25 flask with 15 μg of the lentiviral WβS-reporter vector (7TGC; Addgene #24304; gift from Roel Nusse; see Figure 2A for map) and 15 μg of each of the viral envelope (pMD2.G; Addgene #12259) and viral packaging plasmids (psPAX2, Addgene #12260); both gifts from Didier Trono. Transfections were performed using the Lipofectamine 3000 protocol (ThermoFisher). Viral supernatant was concentrated using the Lenti-X Concentrator (Takara). Primary 2D-cultured FT cells were transduced at p.0 in BM2 medium (see above) containing 8 ug/ml polybrene, for 72 hrs. Transduced cells were selected by FACS and sorted directly into Matrigel. Organoid culture was established and expanded from transduced cells and passaged / expanded as described above. WβS-reporter organoids are available upon request.

### Single cell RNA-sequencing of Wnt/β-catenin reporter organoids

Single cell RNA sequencing (scRNA-seq) was performed on low passage hFT WβS-reporter organoid lines. Organoids were harvested on days 8-11 as described above. RNasin Plus RNase inhibitor (Promega) was included at the harvesting and dissociation steps, to protect RNA integrity while organoids are recovered. Organoids (now single cells) were resuspended in OWB buffer containing RNase inhibitor, 2 mM EDTA and 1% RNase-free BSA (Sigma). Cells were passed through a 30 μm cell strainer and single cell FACS sorting performed using the MA900 Sony Sorter. Isolated WβA or non-WβA cells were sorted into 96-well plates containing 4 μl lysis buffer supplemented with 0.1 μl RNase inhibitor (Clonetech), 1.9 μl 0.4% Triton X-100, 1 μl 10 μM 5‘-biotinylated oligo-dT30VN (IDT) and 1 μl 10 mM dNTP (Thermo Scientific). Cells were sorted at one cell per well, with bulk controls (10 cells) and empty well controls (0 cells) included for each plate. Plates were snap frozen on dry ice and stored at - 80 ^0^C for less than 4 weeks.

Single cell cDNA synthesis and library generation were performed according to the SMART-seq2 protocol (Picelli *et al*., 2014), as previously described (Hu *et al*., 2020). Briefly, cells were lysed by removing plates from -80 ^0^C and heating at 72 ^0^C for 3 min. Plates were then placed at 4 ^0^C before adding the reverse transcription mix containing 5’-biotinylated TSO (Qiagen). PCR products were cleaned up using 0.8:1 Ampure XP beads (Beckman Coulter) with Biomek FxP Laboratory Automation Workstation (Biomek). Quality of single-cell cDNA was tested using TapeStation, as well as by single cell qPCR for GAPDH or ACTB using the QuantiNova SYBR Green PCR Kit (Qiagen). cDNA concentration was measured using Quant-iT^TM^ PicoGreen^TM^ dsDNA Assay Kit (Invitrogen) on the CLARIOstar Plate Reader (BMG Labtech). Wells with C_T_ values of GAPDH or ACTB below 20 were selected as wells with good quality cDNA. Libraries from single cell cDNA were generated using miniaturized Nextera XT (Illumina) protocol (Mora-Castilla *et al*., 2016) with Mosquito HTS (TTP LabTech), in 384-well Endure plate (Life Technology). Library sequencing was performed by Novogene.

### Bulk RNA-seq of Wnt/β-catenin signaling reporter organoids

hFT WβS-reporter organoids were generated and dissociated as described above. RNA extraction and DNase digestion were performed using the RNAqueous-Micro Total RNA Isolation Kit (Thermo Fisher Scientific) according to the manufacturer’s protocol. RNA integrity was evaluated using the 2200 TapeStation system (Agilent). The SMARTer Stranded Total RNA-seq kit v2 - Pico Input (Takara) was used to prepare sequencing libraries, which were then assessed with TapeStation (Agilent) and quantified by Qubit (Thermo Fisher Scientific). Library sequencing was performed by Novogene.

### Computational analysis and statistics

scRNA-seq analysis was performed using R (v4.1.1) and Seurat (v4.3.0) (Hao *et al*., 2021). Genes detected in < 3 cells, cells with UMI counts < 10,000 and gene counts < 200 were removed, resulting in the detection of 16,969 genes in 1,021 cells across a total of 3 patient-derived WβS-reporter organoids, with a median of ∼340 cells from each sample. Raw counts were normalized using the LogNormalize method and ScaleData function with multiple regression variables, including nCount_RNA, S.Score, and G2M.Score. Cells were then clustered using K-nearest neighbor (KNN) graphs and the Louvain algorithm using the first 30 dimensions from principal component analysis. Clustered cells were visualized by UMAP embedding using the default settings. For each sample, differential expression analysis was performed comparing WβA and non-WβA cells using edgeR (v3.36.0) (Robinson *et al*., 2010). Only features detected in >10% of either cell type using FindMarkers were included. Gene expression signatures were derived by identifying genes that exhibited consistent expression level differences, with pvalue < 0.05, across the 3 samples. Gene set enrichment analysis (GSEA; Liberzon *et al*., 2015; Subramanian *et al*., 2005) was performed using the fgsea package (v1.20.0) and MSigDB collections (v2022.1.Hs). Pathways that were upregulated or downregulated in WβA cells relative to non-WβA cells (fdr-adjusted p<0.25) in all samples were selected.

For bulk RNA-seq, sequencing reads from FASTQ files were trimmed for adapter sequences and quality with Trim Galore! and mapped to the UCSC hg19 human genome assembly using STAR (v2.7.3a). Read counts were obtained using subread FeatureCounts (v2.0.0). For differential expression analysis, this was performed using edgeR (v3.36.0) with cut-offs of P < 0.05 and FDR < 0.05. When the analysis was repeated by relaxing the FDR to 0.1, the same list of DEGs was obtained.

### Organoid RNA extraction & RT-qPCR

For RNA extraction, organoids were harvested as described above. Matrigel was solubilized and washed away using the Organoid Harvesting Solution (AmsBio) and cold DPBS. Pelleted organoids were mechanically fragmented in cold DPBS, pelleted, resuspended in 350 μl RLT buffer (Qiagen) and transferred into a 1.5 ml Eppendorf tube. The tube was incubated for 15 min at room temperature in rotation and then vortexed for 1 min. RNA extraction was performed according to the Qiagen RNeasy Plus Micro Kit. Extracted RNA was tested for concentration (using NanoDrop) and, if required, for quality (using TapeStation). Up to 2 μg of extracted

RNA was used to generate cDNA using the High Capacity cDNA Reverse Transcription Kit (Applied Biosystems). RT-qPCR was set up using the SYBR Green PCR Master Mix (ThermoFisher) and conducted using the StepOnePlus RT-PCR machine (ThermoFisher). All data was normalized to endogenous controls *GAPDH* (human) or *Hprt* (mouse), and fold change was quantified by normalization to untreated samples. All qPCR primers used in this study are listed in Table S4.

### RNA In Situ Hybridization (RNAScope)

A small piece of primary human or mouse tissue was resected and embedded in Fisher Healthcare Tissue-Plus Optimum Cutting Temperature (OCT, ThermoFisher). This was frozen at -80 ^0^C, sectioned into 10 μm sections using the CryoStar NX50 (Thermo Scientific) cryostat, mounted on regular glass slides (SuperFrost Plus, VWR International) and immediately stored at - 80 ^0^C. RNA *In situ* Hybridization was performed using the RNAScope® Multiplex Fluorescent v2 kit (ACD) as described (Wang *et al*., 2012) for fresh frozen human or mouse tissue. Organoid sections were established by dissociating organoids, washing and attaching dissociated cells to a slide by Cytospining (FisherScientific) according to the manufacturer’s protocol. Slides were immediately fixed in 4% PFA. RNA FISH staining was performed as per the RNAScope protocol (ACD).

### Organoid Immunofluorescence Staining

Organoids were prepared for antibody staining by culture for 7-12 days on an 8-well microscopy chamber slide (Thistle Scientific). Once ready, whole-mount staining was performed on organoids within Matrigel. Briefly, organoids were washed with PBS and fixed for 15-20 min using 2% methanol-free Paraformaldehyde diluted in DPBS (ThermoFisher). To reduce background staining, samples were washed three times (10 min each) with PBS containing 0.4M Glycine (Sigma). Permeabilization was performed for 10 min using 0.5% Triton X-100 in PBS. All washing, blocking and antibody staining steps were performed in wash buffer containing 0.2% Triton X-100 and 0.05% Tween-20 in PBS. Blocking was done in 5-10% Normal Donkey Serum (Sigma) for 2-3 hrs. Primary antibody incubation was performed overnight at 4 ^0^C in motion. Secondary antibody incubation was done for 2-3 hrs at room temperature. Samples were mounted in Vectashield Mounting Medium (Vector Laboratories). Images were obtained using the Zeiss LSM 780 Inverted Confocal Microscope.

### Wnt-reporter / TOPFlash Assay

All plasmids, small molecule inhibitors or expressed proteins that modulate the WβS pathway were functionally validated using the TOPFlash assay. In brief, 100K-200K HEK 293 cells were reverse transfected with the M50 Super 8x TOPFlash plasmid (100 ng; Addgene #12456; gift from Randall Moon) and a construct constitutively expressing Renilla luciferase (5 ng) in 24-well plates for 48-72 hrs. After that, WβS pathway modulators were introduced at the desired concentrations in fresh medium and incubated overnight. Next day, lysates and luciferase reaction substrates were prepared using the Dual-Luciferase Reporter Assay System (Promega) according to the manufacturer’s protocol, and luciferase readings were acquired using the automated dual injector GloMax Luminometer (Promega). Relative WβS levels were estimated by normalizing

Firefly luciferase to Renilla luciferase readings and normalizing this quotient to the Unt sample. Depending on the experimental set up, one or more plasmids (see key resources / materials table) or siRNA’s (see Table S5) were co-transfected with the above as indicated in the text. For all TOPFlash assays, plasmids were used at 200 ng per well and siRNAs at 40nM, unless otherwise stated. For hormone stimulation experiments, estrogen (17β-estradiol, also called E2, 100 nM) or/and progesterone (P4, 1 μM) were used at the indicated concentrations for 24 hrs.

### Cell Culture & Protein Expression

RSPO1 protein expression and isolation was performed using the HEK293T HA-R-Spondin1-Fc cell line, according to the manufacturer’s protocol (Culturex). WNT3A protein expression was performed using the L-WNT3A cell line according to the manufacturer’s protocol (ATCC). WNT7A protein was expressed by transfecting the pcDNA.WNT7A or empty control plasmids (see key resources / materials table) into HEK293 cells and harvesting the conditioned medium after 96 hrs. For investigating the effect of estrogen and progesterone on WNT7A protein, 800K HEK293 cells were transfected in 6-well plates for 96 hrs with 1 μg of the pcDNA.WNT7A, ERα or PRβ constructs in the combinations indicated in the text. Estrogen (17β-estradiol, also called E2, 100 nM) or/and progesterone (P4, 1 μM) were introduced on transfection day for 96 hrs at the indicated concentrations. Western blot was performed on cell lysates and harvest conditioned medium.

### Key Resources / Materials table

| REAGENT | SOURCE and IDENTIFIER |
| --- | --- |
| **CULTURE MEDIUMS AND REAGENTS** |  |
| Fetal Bovine Serum, qualified, heat inactivated, E | ThermoFisher (#10500064) |
| Penicillin Streptomycin (Pen Strep) | ThermoFisher (#15070063) |
| Lipofectamine 3000 Transfection Reagent | ThermoFisher (#L3000008) |
| Lipofectamine RNAiMAX Transfection Reagent | ThermoFisher (#13778030) |
| Microscope slide, Superfrost plus | VWR (#631-0108) |
| VECTASHIELD anti-fade mounting medium | Vector Laboratories (#H-1000) |
| Immedge Hydrophobic Pen | Vector Laboratories (#H-4000) |
| Dual-Luciferase Reporter Assay System | Promega (#E1910) |
| Brefeldin A (BFA) | ThermoFisher (#00-4506-51) |
| Monensin | ThermoFisher (#00-4505-51) |
| Lenti-X Concentrator | Takari Bio (#631231) |
| **RECOMBINANT PROTEINS & SMALL MOLECULE INHIBITORS** |  |
| CHIR99021 (3 μM) | Bio-Techne (#4423/10) |
| IWP-2 (2 μM) | Bio-Techne (#3533/10) |
| LGK-974 (2 μM) | Strattech Scientific (#S7143-SEL) |
| XAV-939 (5 μM) | Strattech Scientific<br>(#S1180-SEL-10 mg) |
| PRI-724 (10 μM) | Abcam (#ab229168-5mg) |
| Surrogate Wnt (0.5 nM) | UProtein Express (#N001 - 0.025 mg) |
| IgG 2919 (FZD5 inhibitor) | Gift from The Sidhu Lab,<br>University of Toronto |
| IgG 2921 (FZD5 inhibitor) | Gift from The Sidhu Lab,<br>University of Toronto |
| Valproic Acid (1 mM) | Bio-Techne (#2815/100) |
| DAPT (10 μM) | Bio-Techne (#2634/10) |
| Estrogen (100 nM) | Bio-Techne (#2824/100) |
| Progesterone (1 μM) | Bio-Techne (#2835/100) |
| WNT7A Protein, human, recombinant | Abcam (#ab129138) |
| **REAGENTS FOR DISSOCIATION OF PRIMARY TISSUE** |  |
| Trypsin from Bovine pancreas | Sigma-Aldrich (#T9935) |
| Collagenase, Type I | ThermoFisher (#17018029) |
| Dnase I grade II | Sigma-Aldrich (#000000010104159001) |
| Red blood Cell Lysis Solution | Miltenyi Biotech (#130-094-183) |
| **ORGANOID CULTURE** |  |
| Advanced DMEM/F-12, no phenol red | ThermoFisher (#21041033) |
| HEPES | ThermoFisher (#15630056) |
| Penicillin Streptomycin (Pen Strep) | ThermoFisher (#15070063) |
| Epidermal Growth Factor (EGF), human, recombinant | ThermoFisher (#PHG0311) |
| Noggin, human, recombinant | Peprotech (#120-10C) |
| Fibroblast Growth Factor-10 (FGF-10), human, recombinant | Peprotech (#100-26) |
| Human RSPO1 | Peprotech (#120-38-100) |
| Purified mouse Rspo1 | In-house generated |
| N-2 Supplement | ThermoFisher (#17502048) |
| B-27 Supplement | ThermoFisher (#12587010) |
| Nicotinamide | Sigma-Aldrich (#N0636-100G) |
| N-Acetyl L-Cysteine | Sigma-Aldrich (#A9165-5G) |
| Forskolin (FSK) | Bio-Techne (#1099/10) |
| A83.01 | Stemcell Technologies (#72022) |
| Y-27632 (ROCK inhibitor) | Bio-Techne (#1254/10) |
| Cultrex Organoid Harvesting Solution | Bio-Techne (#3700-100-01) |
| Matrigel (phenol red-free) | Corning, supplier SLS (#356237) |
| **PROTEIN EXPRESSION & PLASMIDS** |  |
| L-WNT3A cell line | ATCC (#CRL-2647) |
| HA-R-Spondin1-Fc 293T Cells | AMSBio (#3710-001-01) |
| CD293 medium | ThermoFisher (#11913019) |
| GlutaMAX-1 (100X) | ThermoFisher (#35050-061) |
| M51 Super 8x FOPFlash | Addgene (#12457; gift from Randall Moon) |
| pcDNA.Control plasmid | ThermoFisher (#12489019) |
| pcDNA.WNT7A plasmid | Addgene (#35914; gift from Marian Waterman) |
| pcDNA.WNT7A-V5 plasmid | Addgene (#43816; gift from Xi He) |
| pcDNA.WNT3A plasmid | Addgene (#35908; gift from Marian Waterman) |
| pRK5-mFzd3-1D4 plasmid | Addgene (#42265; gift from Chris Garcia & Jeremy Nathans) |
| pRK5-mFzd5-1D4 plasmid | Addgene (#42267; gift from Chris Garcia & Jeremy Nathans) |
| pRK5-mFzd6-1D4 plasmid | Addgene (#42268; gift from Chris Garcia & Jeremy Nathans) |
| pCMV-hERα plasmid | Addgene (#101141; gift from Elizabeth Wilson) |
| pcDNA3-PRβ plasmid | Addgene (#89130; gift from Elizabeth Wilson) |
| **ANTIBODIES** |  |
| E-cadherin | BD Biosciences (#610181; used 1:200 for IF) |
| Active β-catenin | Merck Millipore (#05-665; used 1:1000 for WB) |
| Ki-67 | Cell Signaling (#9027; used 1:100 for IF) |
| PAX8 | Proteintech (#10336-1-AP; used 1:50 for IF) |
| TUBB4 | Sigma (#T7941; used 1:100 for IF) |
| V5 tag | Abcam (#ab27671; used 1:500 for WB) |
| GAPDH | Proteintech (#60004-1-Ig; used 1:1000 for WB) |
| FcR blocking Reagent, human | Miltenyi Biotec (#130-059-901; used 1:10 for FACS) |
| FcR blocking Reagent, mouse | Miltenyi Biotec (#130-092-575; used 1:10 for FACS) |
| CD45-FITC, human | Biolegend (#304006; used 1:50 for FACS) |
| CD45-FITC, mouse | Miltenyi Biotec (#130-116-500; used 1:50 for FACS) |
| EpCAM-APC, human | Miltenyi Biotec (#130-091-254; used 1:10 for FACS) |
| EpCAM-APC, mouse | Miltenyi Biotec (#130-102-234; used 1:10 for FACS) |
| Phalloidin-Rhodamine |  |
| SINGLE CELL RNA-SEQUENCING |  |
| AMPure XP | Beckman Coulter (#A63881) |
| KAPA HIFI HotStart ReadyMix PCR kit | Roche Diagnostics (#07958935001) |
| SuperScript II Reverse Transcriptase | ThermoFisher (#18064071) |
| Bovine Serum Albumin (BSA, nuclease and protease-free) | Merck Chemicals Ltd (#126609-5GM) |
| Sequencing generated libraries | Novogene |
| RNA EXTRATION AND QPCR |  |
| RNeasy Plus Micro Kit | Qiagen (#74034) |
| High-Capacity cDNA Reverse Transcription Ki | ThermoFisher (#4368814) |
| PowerUp SYBR Green Master Mix | ThermoFisher (#A25780) |
| MicroAmp Fast Optical 96-Well Reaction Plate, 0.1 | ThermoFisher (#4346907) |
| StepOnePlus™ RT PCR System | ThermoFisher (#4376600) |
| RNA IN SITU HYBRIDIZATION (RNAScope) |  |
| Multiplex Fluorescent Detection Kit v2 | Bio-Techne (#323110) |
| SSC solution (20X) | ThermoFisher (#AM9763) |
| ACD HybEZ™ II Hybridization System | Bio-Techne (#321720) |
| Rnase A | Qiagen (#19101) |
| Probe (3-plex human Positive Control) | Bio-Techne (#320861) |
| Probe (3-plex mouse Positive Control) | Bio-Techne (#320881) |
| Probe (3-plex Negative Control) | Bio-Techne (#320871) |
| Probe (human WNT7A) | Bio-Techne (#408231) |
| Probe (human AXIN2) | Bio-Techne (#400241-C2) |
| Probe (human LGR5) | Bio-Techne (#311021-C3) |
| Probe (mouse Wnt7a) | Bio-Techne (#401121) |
| Probe (mouse Axin2) | Bio-Techne (#400331-C2) |
| Probe (mouse Lgr5) | Bio-Techne (#312171-C3) |
| TSA Plus Fluorescein System | PerkinElmer (#NEL741001KT) |
| TSA Plus Cyanine 3 System | PerkinElmer (#NEL744001KT) |
| TSA Plus Cyanine 5 System | PerkinElmer (#NEL745001KT) |

## ABBREVIATIONS

CM: conditioned medium
ER: estrogen receptor
ECM: extracellular matrix
hFT: human fallopian tube
HGSOC: high grade serous ovarian cancer KO knockout
mOV: mouse oviduct
OFE: organoid formation efficiency
RT-qPCR: reverse transcriptase – quantitative polymerase chain reaction
SCT: single cell transcriptomics
WβS: Wnt/β-catenin signaling
WβS-RE: Wnt/β-catenin signaling responsive elements
WβA: Wnt/β-catenin active

## DECLARATIONS

## Acknowledgements

We are grateful to the Weatherall Institute of Molecular Medicine (WIMM) FACS core facility, the WIMM Wolfson Imaging Centre and the WIMM Single Cell core facility for their help in this study. We thank Dr M. Angelica Martinez-Gakidis for valuable editorial feedback.

## Funding

This work was supported by Ovarian Cancer Action, the Oxford Biomedical Research Centre, National Institute of Health Research and the Diane Oxberry Trust.

## Author contributions

Conceptualization, A.A. and A.A.A.; Project Administration, A.A. and A.A.A.; Supervision, A.A.A.; Funding Acquisition, A.A.A. and J.S.B; Investigation, A.A., M.A., Z.H., N.W., M.M., L.S.G., A.A.A.; Data Curation, A.A., M.A., Z.H., N.W., M.M., L.S.G., J.S.B., A.A.A; single cell RNA-seq and analysis, M.A., Z.H., F.S.; bulk RNA-seq and analysis, M.A.; Methodology and Formal Analysis and Visualization, A.A. & A.A.A.; Manuscript Writing (original draft), A.A. and A.A.A.; Manuscript Writing (review & editing), A.A., J.S.B., A.A.A.; Generating anti-FZD5 antibodies, S.S.S., J.A., L.L.B.; Resources (Clinical Samples), N.W., M.M., H.S.M., M.A., J.J., B.A.

## Competing interests

The authors declare no competing interests.

## Notes

### Competing Interest Statement

The authors have declared no competing interest.

### Summary of Updates

- The scRNA-seq data was updated by adding more patient samples of WNT-reporter organoids. - Further methods details were added on the computational analyses, software packages and statistical parameters used in the updated scRNA-seq analysis. A co-author who contributed to this work was added to the list of co-authors. - More functional data were included on estrogen's molecular influences on Fallopian tube renewal and differentiation.

## LIST OF REFERENCES

Alexandre, C., Baena-Lopez, A., & Vincent, J.-P. (2013). Patterning and growth control by membrane-tethered Wingless. Nature 2013 505:7482, 505(7482), 180–185. https://doi.org/10.1038/nature12879

Ali, A., Syed, S. M., Jamaluddin, M. F. B., Colino-Sanguino, Y., Gallego-Ortega, D., & Tanwar, P. S. (2020). Cell Lineage Tracing Identifies Hormone-Regulated and Wnt-Responsive Vaginal Epithelial Stem Cells. Cell Reports, 30(5), 1463–1477.e7. https://doi.org/10.1016/j.celrep.2020.01.003

Angers, S. (2021). Wnt signaling inhibition confers induced synthetic lethality to PARP inhibitors. EMBO Molecular Medicine, 13(4), e14002. https://doi.org/10.15252/EMMM.202114002

Avgustinova, A., Iravani, M., Robertson, D., Fearns, A., Gao, Q., Klingbeil, P., … Isacke, C. M. (2016). Tumour cell-derived Wnt7a recruits and activates fibroblasts to promote tumour aggressiveness. Nature Communications 2016 7:1, 7(1), 1–14. https://doi.org/10.1038/ncomms10305

Bahar-Shany, K., Brand, H., Sapoznik, S., Jacob-Hirsch, J., Yung, Y., Korach, J., … Levanon, K. (2014). Exposure of fallopian tube epithelium to follicular fluid mimics carcinogenic changes in precursor lesions of serous papillary carcinoma. Gynecologic Oncology, 132(2), 322–327. https://doi.org/10.1016/j.ygyno.2013.12.015

Barker, N., Huch, M., Kujala, P., van de Wetering, M., Snippert, H. J., van Es, J. H., … Clevers, H. (2010). Lgr5+ve Stem Cells Drive Self-Renewal in the Stomach and Build Long-Lived Gastric Units In Vitro. Cell Stem Cell, 6(1), 25–36. https://doi.org/10.1016/j.stem.2009.11.013

Barker, N., Van Es, J. H., Kuipers, J., Kujala, P., Van Den Born, M., Cozijnsen, M., … Clevers, H. (2007). Identification of stem cells in small intestine and colon by marker gene Lgr5. Nature, 449(7165), 1003–1007. https://doi.org/10.1038/nature06196

Bartfeld, S., Bayram, T., Van De Wetering, M., Huch, M., Begthel, H., Kujala, P., … Clevers, H. (2015). In vitro expansion of human gastric epithelial stem cells and their responses to bacterial infection. Gastroenterology, 148(1), 126–136.e6. https://doi.org/10.1053/j.gastro.2014.09.042

Battista, S., Guarnieri, D., Borselli, C., Zeppetelli, S., Borzacchiello, A., Mayol, L., … Netti, P. A. (2005). The effect of matrix composition of 3D constructs on embryonic stem cell differentiation. Biomaterials, 26(31), 6194–6207. https://doi.org/10.1016/J.BIOMATERIALS.2005.04.003

Bentzinger, C. F., Wang, Y. X., Von Maltzahn, J., Soleimani, V. D., Yin, H., & Rudnicki, M. A. (2013). Fibronectin Regulates Wnt7a Signaling and Satellite Cell Expansion. Cell Stem Cell, 12(1), 75–87. https://doi.org/10.1016/J.STEM.2012.09.015

Björnström, L., & Sjöberg, M. (2005). Mechanisms of Estrogen Receptor Signaling: Convergence of Genomic and Nongenomic Actions on Target Genes. Molecular Endocrinology, 19(4), 833–842. https://doi.org/10.1210/me.2004-0486

Bond, J., Sedmera, D., Jourdan, J., Zhang, Y., Eisenberg, C. A., Eisenberg, L. M., & Gourdie, R. G. (2003). Wnt11 and Wnt7a are up-regulated in association with differentiation of cardiac conduction cells in vitro and in vivo. Developmental Dynamics, 227(4), 536–543. https://doi.org/10.1002/dvdy.10333

Boonekamp, K. E., Heo, I., Artegiani, B., Asra, P., van Son, G., de Ligt, J., & Clevers, H. (2021). Identification of novel human Wnt target genes using adult endodermal tissue-derived organoids. Developmental Biology. https://doi.org/10.1016/j.ydbio.2021.01.009

Bui, T. D., Zhang, L., Rees, M. C. P., Bicknell, R., & Harris, A. L. (1997). Expression and hormone regulation of Wnt2, 3, 4, 5a, 7a, 7b and 10b in normal human endometrium and endometrial carcinoma. British Journal of Cancer, 75(8), 1131–1136. https://doi.org/10.1038/bjc.1997.195

Bylander, A., Nutu, M., Wellander, R., Goksör, M., Billig, H., & Larsson, D. J. (2010). Rapid effects of progesterone on ciliary beat frequency in the mouse fallopian tube. Reproductive Biology and Endocrinology, 8. https://doi.org/10.1186/1477-7827-8-48

Caricasole, A., Ferraro, T., Iacovelli, L., Barletta, E., Caruso, A., Melchiorri, D., … Nicoletti, F. (2003). Functional Characterization of WNT7A Signaling in PC12 Cells: INTERACTION WITH A FZD5·LRP6 RECEPTOR COMPLEX AND MODULATION BY DICKKOPF PROTEINS. Journal of Biological Chemistry, 278(39), 37024–37031. https://doi.org/10.1074/JBC.M300191200

Carmon, K. S., Gong, X., Lin, Q., Thomas, A., & Liu, Q. (2011). R-spondins function as ligands of the orphan receptors LGR4 and LGR5 to regulate Wnt/β-catenin signaling. Proceedings of the National Academy of Sciences of the United States of America, 108(28), 11452–11457. https://doi.org/10.1073/pnas.1106083108

Carmon, K. S., & Loose, D. S. (2008). Wnt7a interaction with Fzd5 and detection of signaling activation using a split eGFP. Biochemical and Biophysical Research Communications, 368(2), 285–291. https://doi.org/10.1016/J.BBRC.2008.01.088

Clevers, H. (2006). Wnt/β-Catenin Signaling in Development and Disease. Cell, 127(3), 469–480. https://doi.org/10.1016/J.CELL.2006.10.018

Clevers, H., Loh, K. M., & Nusse, R. (2014). An integral program for tissue renewal and regeneration: Wnt signaling and stem cell control. Science, 346(6205). https://doi.org/10.1126/SCIENCE.1248012/ASSET/2B86C789-E77F-4290-AF68-B234AA81AFD5/ASSETS/GRAPHIC/346_1248012_F4.JPEG

Coan, M., Vinciguerra, G. L. R., Cesaratto, L., Gardenal, E., Bianchet, R., Dassi, E., … Nicoloso, M. S. (2018). Exploring the role of fallopian ciliated cells in the pathogenesis of high-grade serous ovarian cancer. International Journal of Molecular Sciences, 19(9). https://doi.org/10.3390/IJMS19092512

Couse, J. F., Dixon, D., Yates, M., Moore, A. B., Ma, L., Maas, R., & Korach, K. S. (2001). Estrogen receptor-α knockout mice exhibit resistance to the developmental effects of neonatal diethylstilbestrol exposure on the female reproductive tract. Developmental Biology, 238(2), 224–238. https://doi.org/10.1006/dbio.2001.0413

de Lau, W., Barker, N., Low, T. Y., Koo, B.-K., Li, V. S. W., Teunissen, H., … Clevers, H. (2011). Lgr5 homologues associate with Wnt receptors and mediate R-spondin signalling. Nature, 476(7360), 293–297. https://doi.org/10.1038/nature10337

de Witte, C. J., Espejo Valle-Inclan, J., Hami, N., Lõhmussaar, K., Kopper, O., Vreuls, C. P. H., … Stelloo, E. (2020). Patient-Derived Ovarian Cancer Organoids Mimic Clinical Response and Exhibit Heterogeneous Inter- and Intrapatient Drug Responses. Cell Reports, 31(11), 107762. https://doi.org/10.1016/j.celrep.2020.107762

DeCherney, A. H., Cholst, I., & Naftolin, F. (1981). Structure and function of the fallopian tubes following exposure to diethylstilbestrol (DES) during gestation. Fertility and Sterility, 36(6), 741–745. https://doi.org/10.1016/S0015-0282(16)45919-4

Degirmenci, B., Valenta, T., Dimitrieva, S., Hausmann, G., & Basler, K. (2018). GLI1-expressing mesenchymal cells form the essential Wnt-secreting niche for colon stem cells. Nature, 558(7710), 449–453. https://doi.org/10.1038/s41586-018-0190-3

Dinh, H. Q., Lin, X., Abbasi, F., Nameki, R., Haro, M., Olingy, C. E., … Lawrenson, K. (2021). Single-cell transcriptomics identifies gene expression networks driving differentiation and tumorigenesis in the human fallopian tube. Cell Reports, 35(2), 108978. https://doi.org/10.1016/j.celrep.2021.108978

Dong, B., Vold, S., Olvera-Jaramillo, C., & Chang, H. (2018). Functional redundancy of frizzled 3 and frizzled 6 in planar cell polarity control of mouse hair follicles. Development (Cambridge), 145(19). https://doi.org/10.1242/dev.168468

Dunlap, K. A., Filant, J., Hayashi, K., Rucker, E. B., Song, G., Deng, J. M., … Spencer, T. E. (2011). Postnatal Deletion of Wnt7a Inhibits Uterine Gland Morphogenesis and Compromises Adult Fertility in Mice1. Biology of Reproduction, 85(2), 386–396. https://doi.org/10.1095/biolreprod.111.091769

Enuka, Y., Hanukoglu, I., Edelheit, O., Vaknine, H., & Hanukoglu, A. (2012). Epithelial sodium channels (ENaC) are uniformly distributed on motile cilia in the oviduct and the respiratory airways. Histochemistry and Cell Biology, 137(3), 339–353. https://doi.org/10.1007/S00418-011-0904-1

Erickson, B. K., Conner, M. G., & Landen, C. N. (2013). The role of the fallopian tube in the origin of ovarian cancer. American Journal of Obstetrics and Gynecology, Vol. 209, pp. 409–414. https://doi.org/10.1016/j.ajog.2013.04.019

Farin, H. F., Jordens, I., Mosa, M. H., Basak, O., Korving, J., Tauriello, D. V. F., … Clevers, H. (2016). Visualization of a short-range Wnt gradient in the intestinal stem-cell niche. Nature, 530(7590), 340–343. https://doi.org/10.1038/nature16937

Gattazzo, F., Urciuolo, A., & Bonaldo, P. (2014). Extracellular matrix: A dynamic microenvironment for stem cell niche. Biochimica et Biophysica Acta (BBA) - General Subjects, 1840(8), 2506–2519. https://doi.org/10.1016/J.BBAGEN.2014.01.010

George, S. H. L., Milea, A., Sowamber, R., Chehade, R., Tone, A., & Shaw, P. A. (2015). Loss of LKB1 and p53 synergizes to alter fallopian tube epithelial phenotype and high-grade serous tumorigenesis. Oncogene 2016 35:1, 35(1), 59–68. https://doi.org/10.1038/onc.2015.62

Ghosh, A., Syed, S. M., & Tanwar, P. S. (2017). In vivo genetic cell lineage tracing reveals that oviductal secretory cells self-renew and give rise to ciliated cells. https://doi.org/10.1242/dev.149989

Glinka, A., Dolde, C., Kirsch, N., Huang, Y. L., Kazanskaya, O., Ingelfinger, D., … Niehrs, C. (2011). LGR4 and LGR5 are R-spondin receptors mediating Wnt/β-catenin and Wnt/PCP signalling. EMBO Reports, 12(10), 1055–1061. https://doi.org/10.1038/embor.2011.175

Goldstein, B., Takeshita, H., Mizumoto, K., & Sawa, H. (2006). Wnt signals can function as positional cues in establishing cell polarity. Developmental Cell, 10(3), 391–396. https://doi.org/10.1016/j.devcel.2005.12.016

Guilak, F., Cohen, D. M., Estes, B. T., Gimble, J. M., Liedtke, W., & Chen, C. S. (2009). Control of Stem Cell Fate by Physical Interactions with the Extracellular Matrix. Cell Stem Cell, 5(1), 17–26. https://doi.org/10.1016/J.STEM.2009.06.016

Hall, A. C., Lucas, F. R., & Salinas, P. C. (2000). Axonal remodeling and synaptic differentiation in the cerebellum is regulated by WNT-7a signaling. Cell, 100(5), 525–535. https://doi.org/10.1016/S0092-8674(00)80689-3

Hao, H. X., Xie, Y., Zhang, Y., Zhang, O., Oster, E., Avello, M., … Cong, F. (2012). ZNRF3 promotes Wnt receptor turnover in an R-spondin-sensitive manner. Nature, 485(7397), 195–202. https://doi.org/10.1038/nature11019

Hao, Y., Hao, S., Andersen-Nissen, E., Mauck, W. M., Zheng, S., Butler, A., … Satija, R. (2021). Integrated analysis of multimodal single-cell data. Cell, 184(13), 3573–3587.e29. https://doi.org/10.1016/J.CELL.2021.04.048/ATTACHMENT/1E5EB5C1-59EE-4B2B-8BFA-14B48A54FF8F/MMC3.XLSX

He, F., Xiong, W., Yu, X., Espinoza-Lewis, R., Liu, C., Gu, S., … Chen, Y. P. (2008). Wnt5a regulates directional cell migration and cell proliferation via Ror2-mediated noncanonical pathway in mammalian palate development. Development, 135(23), 3871–3879. https://doi.org/10.1242/dev.025767

Hideyuki, N., Saito, T., Yamasaki, H., Mizumoto, H., Ito, E., & Kudo, R. (1999). Nuclear localization of β-catenin in normal and carcinogenic endometrium. Molecular Carcinogenesis, 25(3), 207–218. https://doi.org/10.1002/(SICI)1098-2744(199907)25:3<207::AID-MC7>3.0.CO;2-4

Hill, S. J., Decker, B., Roberts, E. A., Horowitz, N. S., Muto, M. G., Worley, M. J., … D’Andrea, A. D. (2018). Prediction of DNA repair inhibitor response in short-term patient-derived ovarian cancer organoids. Cancer Discovery, 8(11), 1404–1421. https://doi.org/10.1158/2159-8290.CD-18-0474

Hoffmann, K., Berger, H., Kulbe, H., Thillainadarasan, S., Mollenkopf, H., Zemojtel, T., … Kessler, M. (2020). Stable expansion of high-grade serous ovarian cancer organoids requires a low-Wnt environment. The EMBO Journal, e104013, 1–23. https://doi.org/10.15252/embj.2019104013

Hou, Y. F., Yuan, S. T., Li, H. C., Wu, J., Lu, J. S., Liu, G., … Shao, Z. M. (2004). ERβ exerts multiple stimulative effects on human breast carcinoma cells. Oncogene, 23(34), 5799–5806. https://doi.org/10.1038/sj.onc.1207765

Hu, Z., Artibani, M., Alsaadi, A., Wietek, N., Morotti, M., Shi, T., … Ahmed, A. A. (2020). The Repertoire of Serous Ovarian Cancer Non-genetic Heterogeneity Revealed by Single-Cell Sequencing of Normal Fallopian Tube Epithelial Cells. Cancer Cell, 37(2), 226–242.e7. https://doi.org/10.1016/j.ccell.2020.01.003

Huang, H. S., Chu, S. C., Hsu, C. F., Chen, P. C., Ding, D. C., Chang, M. Y., & Chu, T. Y. (2015). Mutagenic, surviving and tumorigenic effects of follicular fluid in the context of p53 loss: initiation of fimbria carcinogenesis. Carcinogenesis, 36(11), 1419–1428. https://doi.org/10.1093/carcin/bgv132

Huang, S. M. A., Mishina, Y. M., Liu, S., Cheung, A., Stegmeier, F., Michaud, G. A., … Cong, F. (2009). Tankyrase inhibition stabilizes axin and antagonizes Wnt signalling. Nature, 461(7264), 614–620. https://doi.org/10.1038/nature08356

Huang, X., Zhu, H., Gao, Z., Li, J., Zhuang, J., Dong, Y., … Yan, J. (2018). Wnt7a activates canonical Wnt signaling, promotes bladder cancer cell invasion, and is suppressed by MIR-370-3p. Journal of Biological Chemistry, 293(18), 6693–6706. https://doi.org/10.1074/jbc.RA118.001689

Huch, M., Dorrell, C., Boj, S. F., Van Es, J. H., Li, V. S. W., Van De Wetering, M., … Clevers, H. (2013). In vitro expansion of single Lgr5 + liver stem cells induced by Wnt-driven regeneration. Nature, 494(7436), 247–250. https://doi.org/10.1038/nature11826

Huch, M., Gehart, H., van Boxtel, R., Hamer, K., Blokzijl, F., Verstegen, M. M. A., … Clevers, H. (2015). Long-term culture of genome-stable bipotent stem cells from adult human liver. Cell, 160(1–2), 299–312. https://doi.org/10.1016/j.cell.2014.11.050

Hwang, S. G., Ryu, J. H., Kim, I. C., Jho, E. H., Jung, H. C., Kim, K., … Chun, J. S. (2004). Wnt-7a causes loss of differentiated phenotype and inhibits apoptosis of articular chondrocytes via different mechanisms. Journal of Biological Chemistry, 279(25), 26597–26604. https://doi.org/10.1074/jbc.M401401200

Ingaramo, P. I., Milesi, M. M., Schimpf, M. G., Ramos, J. G., Vigezzi, L., Muñoz-de-Toro, M., … Varayoud, J. (2016). Endosulfan affects uterine development and functional differentiation by disrupting Wnt7a and β-catenin expression in rats. Molecular and Cellular Endocrinology, 425, 37–47. https://doi.org/10.1016/j.mce.2016.02.011

Janda, C. Y., Dang, L. T., You, C., Chang, J., Lau, W. De, Zhong, Z. A., … Garcia, K. C. (2017). Surrogate Wnt agonists that phenocopy canonical Wnt and β-catenin signalling. Nature, 545(7653), 234–237. https://doi.org/10.1038/nature22306

Janson, I. A., & Putnam, A. J. (2015). Extracellular matrix elasticity and topography: Material-based cues that affect cell function via conserved mechanisms. Journal of Biomedical Materials Research Part A, 103(3), 1246–1258. https://doi.org/10.1002/JBM.A.35254

Jia, B., Qiu, X., Chu, H., Sun, X., Xu, S., Zhao, X., & Zhao, J. (2019). Wnt7a predicts poor prognosis, and contributes to growth and metastasis in tongue squamous cell carcinoma. Oncology Reports, 41(3), 1749–1758. https://doi.org/10.3892/OR.2019.6974/HTML

Karthaus, W. R., Iaquinta, P. J., Drost, J., Gracanin, A., Van Boxtel, R., Wongvipat, J., … Clevers, H. C. (2014). Identification of multipotent luminal progenitor cells in human prostate organoid cultures. Cell, 159(1), 163–175. https://doi.org/10.1016/j.cell.2014.08.017

Kaur, A., Lim, J. Y. S., Sepramaniam, S., Patnaik, S., Harmston, N., Lee, M. A., … Madan, B. (2021). WNT inhibition creates a BRCA-like state in Wnt-addicted cancer. EMBO Molecular Medicine, 13(4). https://doi.org/10.15252/EMMM.202013349

Kessler, M., Hoffmann, K., Brinkmann, V., Thieck, O., Jackisch, S., Toelle, B., … Meyer, T. F. (2015). The Notch and Wnt pathways regulate stemness and differentiation in human fallopian tube organoids. Nature Communications, 6(May), 8989. https://doi.org/10.1038/ncomms9989

Kessler, M., Hoffmann, K., Fritsche, K., Brinkmann, V., Mollenkopf, H.-J., Thieck, O., … Meyer, T. F. (2019). Chronic Chlamydia infection in human organoids increases stemness and promotes age-dependent CpG methylation. Nature Communications, 10(1), 1194. https://doi.org/10.1038/s41467-019-09144-7

Kim, J., Park, E. Y., Kim, O., Schilder, J. M., Coffey, D. M., Cho, C. H., & Bast, R. C. (2018, November 12). Cell origins of high-grade serous ovarian cancer. Cancers, Vol. 10. https://doi.org/10.3390/cancers10110433

King, S. M., Hilliard, T. S., Wu, L. Y., Jaffe, R. C., Fazleabas, A. T., & Burdette, J. E. (2011). The impact of ovulation on fallopian tube epithelial cells: Evaluating three hypotheses connecting ovulation and serous ovarian cancer. Endocrine-Related Cancer, 18(5), 627–642. https://doi.org/10.1530/ERC-11-0107

Koo, B. K., Spit, M., Jordens, I., Low, T. Y., Stange, D. E., Van De Wetering, M., … Clevers, H. (2012). Tumour suppressor RNF43 is a stem-cell E3 ligase that induces endocytosis of Wnt receptors. Nature, 488(7413), 665–669. https://doi.org/10.1038/nature11308

Kopper, O., de Witte, C. J., Lõhmussaar, K., Valle-Inclan, J. E., Hami, N., Kester, L., … Clevers, H. (2019). An organoid platform for ovarian cancer captures intra- and interpatient heterogeneity. Nature Medicine, 1. https://doi.org/10.1038/s41591-019-0422-6

Kouzmenko, A. P., Takeyama, K. I., Ito, S., Furutani, T., Sawatsubashi, S., Maki, A., … Kato, S. (2004). Wnt/β-catenin and estrogen signaling converge in vivo. Journal of Biological Chemistry, 279(39), 40255–40258. https://doi.org/10.1074/jbc.C400331200

Kroeger, P. T., & Drapkin, R. (2017). Pathogenesis and heterogeneity of ovarian cancer. Current Opinion in Obstetrics and Gynecology, 29(1), 26–34. https://doi.org/10.1097/GCO.0000000000000340

Kuhn, E., Kurman, R. J., Vang, R., Sehdev, A. S., Han, G., Soslow, R., … Shih, I.-M. (2012). TP53 mutations in serous tubal intraepithelial carcinoma and concurrent pelvic high-grade serous carcinoma-evidence supporting the clonal relationship of the two lesions. The Journal of Pathology, 226(3), 421–426. https://doi.org/10.1002/path.3023

Labidi-Galy, S. I., Papp, E., Hallberg, D., Niknafs, N., Adleff, V., Noe, M., … Velculescu, V. E. (2017). High grade serous ovarian carcinomas originate in the fallopian tube. Nature Communications, 8(1), 1093. https://doi.org/10.1038/s41467-017-00962-1

Leushacke, M., Tan, S. H., Wong, A., Swathi, Y., Hajamohideen, A., Tan, L. T., … Barker, N. (2017). Lgr5-expressing chief cells drive epithelial regeneration and cancer in the oxyntic stomach. Nature Cell Biology, 19(7), 774–786. https://doi.org/10.1038/ncb3541

Li, H. W. R., Liao, S. B., Chiu, P. C. N., Tam, W. W., Ho, J. C., Ng, E. H. Y., … Wai Sum, O. (2010). Expression of adrenomedullin in human oviduct, its regulation by the hormonal cycle and contact with spermatozoa, and its effect on ciliary beat frequency of the oviductal epithelium. Journal of Clinical Endocrinology and Metabolism, 95(9). https://doi.org/10.1210/JC.2010-0273

Li, J., Ning, Y., Abushahin, N., Yuan, Z., Wang, Y., Wang, Y., … Zheng, W. (2013). Secretory cell expansion with aging: Risk for pelvic serous carcinogenesis. Gynecologic Oncology, 131(3), 555–560. https://doi.org/10.1016/J.YGYNO.2013.09.018

Liberzon, A., Birger, C., Thorvaldsdóttir, H., Ghandi, M., Mesirov, J. P., & Tamayo, P. (2015). The Molecular Signatures Database Hallmark Gene Set Collection. Cell Systems, 1(6), 417– 425. https://doi.org/10.1016/j.cels.2015.12.004

Lim, R., Banerjee, A., Biswas, R., Chari, A. N., & Raghavan, S. (2022). Mechanotransduction through adhesion molecules: Emerging roles in regulating the stem cell niche. Frontiers in Cell and Developmental Biology, 10. https://doi.org/10.3389/FCELL.2022.966662

Liu, Y., Meng, F., Xu, Y., Yang, S., Xiao, M., Chen, X., & Lou, G. (2013). Overexpression of wnt7a is associated with tumor progression and unfavorable prognosis in endometrial cancer. International Journal of Gynecological Cancer, 23(2), 304–311. https://doi.org/10.1097/IGC.0b013e31827c7708

Lõhmussaar, K., Kopper, O., Korving, J., Begthel, H., Vreuls, C. P. H., van Es, J. H., & Clevers, H. (2020). Assessing the origin of high-grade serous ovarian cancer using CRISPR-modification of mouse organoids. Nature Communications, 11(1), 2660. https://doi.org/10.1038/s41467-020-16432-0

Long, K., Moss, L., Laursen, L., Boulter, L., & Ffrench-Constant, C. (2016). Integrin signalling regulates the expansion of neuroepithelial progenitors and neurogenesis via Wnt7a and Decorin. Nature Communications 2016 7:1, 7(1), 1–14. https://doi.org/10.1038/ncomms10354

Lu, J., Zhang, S., Nakano, H., Simmons, D. G., Wang, S., Kong, S., … Wang, H. (2013). A positive feedback loop involving Gcm1 and Fzd5 directs chorionic branching morphogenesis in the placenta. PLoS Biology, 11(4). https://doi.org/10.1371/JOURNAL.PBIO.1001536

Lyons, R. A., Saridogan, E., & Djahanbakhch, O. (2006). The reproductive significance of human Fallopian tube cilia. Human Reproduction Update, 12(4), 363–372. https://doi.org/10.1093/HUMUPD/DML012

McConnell, A. M., Yao, C., Yeckes, A. R., Wang, Y., Selvaggio, A. S., Tang, J., … Stripp, B. R. (2016). p53 Regulates Progenitor Cell Quiescence and Differentiation in the Airway. Cell Reports, 17(9), 2173–2182. https://doi.org/10.1016/J.CELREP.2016.11.007

McLachlan, J. A., Newbold, R. R., & Bullock, B. C. (1980). Long-Term Effects on the Female Mouse Genital Tract Associated with Prenatal Exposure to Diethylstilbestrol. Cancer Research, 40(11).

Mericskay, M., Kitajewski, J., & Sassoon, D. (2004). Wnt5a is required for proper epithelial-mesenchymal interactions in the uterus. Development, 131(9), 2061–2072. https://doi.org/10.1242/dev.01090

Miller, C., Degenhardt, K., & Sassoon, D. A. (1998). Fetal exposure to DES results in de-regulation of Wnt7a during uterine morphogenesis [2]. Nature Genetics, Vol. 20, pp. 228–230. https://doi.org/10.1038/3027

Miller, C., & Sassoon, D. A. (1998). Wnt-7a maintains appropriate uterine patterning during the development of the mouse female reproductive tract. Development, 125(16).

Miller, Cary, Pavlova, A., & Sassoon, D. A. (1998). Differential expression patterns of Wnt genes in the murine female reproductive tract during development and the estrous cycle. Mechanisms of Development, 76(1–2), 91–99. https://doi.org/10.1016/S0925-4773(98)00112-9

Miller, M. F., Cohen, E. D., Baggs, J. E., Lu, M. M., Hogenesch, J. B., & Morrisey, E. E. (2012). Wnt ligands signal in a cooperative manner to promote foregut organogenesis. Proceedings of the National Academy of Sciences of the United States of America, 109(38), 15348– 15353. https://doi.org/10.1073/PNAS.1201583109/SUPPL_FILE/PNAS.201201583SI.PDF

Mora-Castilla, S., To, C., Vaezeslami, S., Morey, R., Srinivasan, S., Dumdie, J. N., … Laurent, L. C. (2016). Miniaturization Technologies for Efficient Single-Cell Library Preparation for Next-Generation Sequencing. Journal of Laboratory Automation, 21(4), 557–567. https://doi.org/10.1177/2211068216630741

Moyle, L. A., Cheng, R. Y., Liu, H., Davoudi, S., Ferreira, S. A., Nissar, A. A., … Gilbert, P. M. (2020). Three-dimensional niche stiffness synergizes with Wnt7a to modulate the extent of satellite cell symmetric self-renewal divisions. Molecular Biology of the Cell, 31(16), 1703–1713. https://doi.org/10.1091/MBC.E20-01-0078/ASSET/IMAGES/LARGE/MBC-31-1703-G004.JPEG

Ng, A., Tan, S., Singh, G., Rizk, P., Swathi, Y., Tan, T. Z., … Barker, N. (2014). Lgr5 marks stem/progenitor cells in ovary and tubal epithelia. Nature Cell Biology, 16(8), 745–757. https://doi.org/10.1038/ncb3000

Okazaki, H., Sato, S., Koyama, K., Morizumi, S., Abe, S., Azuma, M., … Nishioka, Y. (2019). The novel inhibitor PRI-724 for Wnt/β-catenin/CBP signaling ameliorates bleomycin-induced pulmonary fibrosis in mice. Experimental Lung Research, 45(7), 188–199. https://doi.org/10.1080/01902148.2019.1638466

Olson, D. J., & Gibo, D. M. (1998). Antisense wnt-5a mimics wnt-1-mediated C57MG mammary epithelial cell transformation. Experimental Cell Research, 241(1), 134–141. https://doi.org/10.1006/excr.1998.4030

Paik, D. Y., Janzen, D. M., Schafenacker, A. M., Velasco, V. S., Shung, M. S., Cheng, D., … Memarzadeh, S. (2012). Stem-Like Epithelial Cells Are Concentrated in the Distal End of the Fallopian Tube: A Site for Injury and Serous Cancer Initiation. STEM CELLS, 30(11), 2487–2497. https://doi.org/10.1002/stem.1207

Park, S., Cui, J., Yu, W., Wu, L., Carmon, K. S., & Liu, Q. J. (2018). Differential activities and mechanisms of the four r-spondins in potentiating wnt/-catenin signaling. Journal of Biological Chemistry, 293(25), 9759–9769. https://doi.org/10.1074/JBC.RA118.002743/ATTACHMENT/7D21F42A-0024-4AE9-A571-DB0E05669870/MMC1.PDF

Parr, B. A., & McMahon, A. P. (1998). Sexually dimorphic development of the mammalian reproductive tract requires Wnt-7a. Nature, 395(6703), 707–710. https://doi.org/10.1038/27221

Picelli, S., Faridani, O. R., Björklund, Å. K., Winberg, G., Sagasser, S., & Sandberg, R. (2014). Full-length RNA-seq from single cells using Smart-seq2. Nature Protocols, 9(1), 171–181. https://doi.org/10.1038/nprot.2014.006

Raidt, J., Werner, C., Menchen, T., Dougherty, G. W., Olbrich, H., Loges, N. T., … Omran, H. (2015). Ciliary function and motor protein composition of human fallopian tubes. Human Reproduction, 30(12), 2871–2880. https://doi.org/10.1093/HUMREP/DEV227

Robinson, M. D., McCarthy, D. J., & Smyth, G. K. (2010). edgeR: a Bioconductor package for differential expression analysis of digital gene expression data. Bioinformatics, 26(1), 139. https://doi.org/10.1093/BIOINFORMATICS/BTP616

Sakaki-Yumoto, M., Katsuno, Y., & Derynck, R. (2013, February). TGF-β family signaling in stem cells. Biochimica et Biophysica Acta - General Subjects, Vol. 1830, pp. 2280–2296. https://doi.org/10.1016/j.bbagen.2012.08.008

Sato, T., van Es, J. H., Snippert, H. J., Stange, D. E., Vries, R. G., van den Born, M., … Clevers, H. (2011). Paneth cells constitute the niche for Lgr5 stem cells in intestinal crypts. Nature, 469(7330), 415–418. https://doi.org/10.1038/nature09637

Sato, T., Vries, R. G., Snippert, H. J., Van De Wetering, M., Barker, N., Stange, D. E., … Clevers, H. (2009). Single Lgr5 stem cells build crypt-villus structures in vitro without a mesenchymal niche. Nature, 459(7244), 262–265. https://doi.org/10.1038/nature07935

Seishima, R., Leung, C., Yada, S., Murad, K. B. A., Tan, L. T., Hajamohideen, A., … Barker, N. (2019). Neonatal Wnt-dependent Lgr5 positive stem cells are essential for uterine gland development. Nature Communications, 10(1). https://doi.org/10.1038/s41467-019-13363-3

Shoshkes-Carmel, M., Wang, Y. J., Wangensteen, K. J., Tóth, B., Kondo, A., Massassa, E. E., … Kaestner, K. H. (2018). Subepithelial telocytes are an important source of Wnts that supports intestinal crypts. Nature, 557(7704), 242–246. https://doi.org/10.1038/s41586-018-0084-4

Sigal, M., Logan, C. Y., Kapalczynska, M., Mollenkopf, H. J., Berger, H., Wiedenmann, B., … Meyer, T. F. (2017). Stromal R-spondin orchestrates gastric epithelial stem cells and gland homeostasis. Nature, 548(7668), 451–455. https://doi.org/10.1038/nature23642

Snegovskikh, V., Mutlu, L., Massasa, E., & Taylor, H. S. (2014). Identification of putative fallopian tube stem cells. Reproductive Sciences, 21(12), 1460–1464. https://doi.org/10.1177/1933719114553448

Stange, D. E., Koo, B. K., Huch, M., Sibbel, G., Basak, O., Lyubimova, A., … Clevers, H. (2013). Differentiated Troy+ chief cells act as reserve stem cells to generate all lineages of the stomach epithelium. Cell, 155(2), 357. https://doi.org/10.1016/j.cell.2013.09.008

Steinhart, Z., Pavlovic, Z., Chandrashekhar, M., Hart, T., Wang, X., Zhang, X., … Angers, S. (2017). Genome-wide CRISPR screens reveal a Wnt–FZD5 signaling circuit as a druggable vulnerability of RNF43-mutant pancreatic tumors. Nature Medicine, 23(1), 60–68. https://doi.org/10.1038/nm.4219

Subramanian, A., Tamayo, P., Mootha, V. K., Mukherjee, S., Ebert, B. L., Gillette, M. A., … Mesirov, J. P. (2005). Gene set enrichment analysis: A knowledge-based approach for interpreting genome-wide expression profiles. Proceedings of the National Academy of Sciences of the United States of America, 102(43), 15545. https://doi.org/10.1073/PNAS.0506580102

Syed, S. M., Kumar, M., Ghosh, A., Tomasetig, F., Ali, A., Whan, R. M., … Tanwar, P. S. (2020). Endometrial Axin2+ Cells Drive Epithelial Homeostasis, Regeneration, and Cancer following Oncogenic Transformation. Cell Stem Cell, 26(1), 64–80.e13. https://doi.org/10.1016/j.stem.2019.11.012

Tojo, M., Hamashima, Y., Hanyu, A., Kajimoto, T., Saitoh, M., Miyazono, K., … Imamura, T. (2005). The ALK-5 inhibitor A-83-01 inhibits Smad signaling and epithelial-to-mesenchymal transition by transforming growth factor-beta. Cancer Science, 96(11), 791–800. https://doi.org/10.1111/j.1349-7006.2005.00103.x

Topol, L., Jiang, X., Choi, H., Garrett-Beal, L., Carolan, P. J., & Yang, Y. (2003). Wnt-5a inhibits the canonical Wnt pathway by promoting GSK-3-independent β-catenin degradation. Journal of Cell Biology, 162(5), 899–908. https://doi.org/10.1083/jcb.200303158

Turco, M. Y., Gardner, L., Hughes, J., Cindrova-Davies, T., Gomez, M. J., Farrell, L., … Burton, G. J. (2017). Long-term, hormone-responsive organoid cultures of human endometrium in a chemically defined medium. Nature Cell Biology, 19(5), 568–577. https://doi.org/10.1038/ncb3516

Tüysüz, N., van Bloois, L., van den Brink, S., Begthel, H., Verstegen, M. M. A., Cruz, L. J., … ten Berge, D. (2017). Lipid-mediated Wnt protein stabilization enables serum-free culture of human organ stem cells. Nature Communications, 8, 14578. https://doi.org/10.1038/ncomms14578

Urciuolo, A., Quarta, M., Morbidoni, V., Gattazzo, F., Molon, S., Grumati, P., … Bonaldo, P. (2013). Collagen VI regulates satellite cell self-renewal and muscle regeneration. Nature Communications 2013 4:1, 4(1), 1–13. https://doi.org/10.1038/ncomms2964

Van Camp, J. K., Beckers, S., Zegers, D., & Van Hul, W. (2014). Wnt Signaling and the Control of Human Stem Cell Fate. Stem Cell Reviews and Reports, 10(2), 207–229. https://doi.org/10.1007/S12015-013-9486-8/FIGURES/3

van den Heuvel, M., Nusse, R., Johnston, P., & Lawrence, P. A. (1989). Distribution of the wingless gene product in drosophila embryos: A protein involved in cell-cell communication. Cell, 59(4), 739–749. https://doi.org/10.1016/0092-8674(89)90020-2

Voloshanenko, O., Gmach, P., Winter, J., Kranz, D., & Boutros, M. (2017). Mapping of Wnt-Frizzled interactions by multiplex CRISPR targeting of receptor gene families. The FASEB Journal, 31(11), 4832–4844. https://doi.org/10.1096/fj.201700144R

Wagner, J., & Lehmann, L. (2006). Estrogens modulate the gene expression of Wnt-7a in cultured endometrial adenocarcinoma cells. Molecular Nutrition and Food Research, 50(4–5), 368–372. https://doi.org/10.1002/mnfr.200500215

Wang, B., Zhao, L., Fish, M., Logan, C. Y., & Nusse, R. (2015). Self-renewing diploid Axin2 + cells fuel homeostatic renewal of the liver. Nature, 524(7564), 180–185. https://doi.org/10.1038/nature14863

Wang, F., Flanagan, J., Su, N., Wang, L.-C., Bui, S., Nielson, A., … Luo, Y. (2012). RNAscope: A Novel in Situ RNA Analysis Platform for Formalin-Fixed, Paraffin-Embedded Tissues. The Journal of Molecular Diagnostics, 14(1), 22–29. https://doi.org/10.1016/j.jmoldx.2011.08.002

Wang, Y., Sacchetti, A., Dijk, M. R. van, Zee, M. van der, Horst, P. H. van der, Joosten, R., … Fodde, R. (2012). Identification of Quiescent, Stem-Like Cells in the Distal Female Reproductive Tract. PLoS ONE, 7(7), e40691. https://doi.org/10.1371/JOURNAL.PONE.0040691

Watt, F. M., & Huck, W. T. S. (2013). Role of the extracellular matrix in regulating stem cell fate. Nature Reviews Molecular Cell Biology 2013 14:8, 14(8), 467–473. https://doi.org/10.1038/nrm3620

Xie, Y., Park, E.-S., Xiang, D., & Li, Z. (2018). Long-term organoid culture reveals enrichment of organoid-forming epithelial cells in the fimbrial portion of mouse fallopian tube. Stem Cell Research, 32, 51–60. https://doi.org/10.1016/J.SCR.2018.08.021

Yamamoto, Y., Ning, G., Howitt, B. E., Mehra, K., Wu, L., Wang, X., … Xian, W. (2016). *In vitro* and *in vivo* correlates of physiological and neoplastic human Fallopian tube stem cells. The Journal of Pathology, 238(4), 519–530. https://doi.org/10.1002/path.4649

Yin, X., Farin, H. F., van Es, J. H., Clevers, H., Langer, R., & Karp, J. M. (2013). Niche-independent high-purity cultures of Lgr5+ intestinal stem cells and their progeny. Nature Methods, 11(1), 106–112. https://doi.org/10.1038/nmeth.2737

Yoshioka, S., King, M. L., Ran, S., Okuda, H., MacLean, J. A., McAsey, M. E., … Hayashi, K. (2012). WNT7A Regulates Tumor Growth and Progression in Ovarian Cancer through the WNT/ - Catenin Pathway. Molecular Cancer Research, 10(3), 469–482. https://doi.org/10.1158/1541-7786.MCR-11-0177

You, Y., Huang, T., Richer, E. J., Schmidt, J. E. H., Zabner, J., Borok, Z., & Brody, S. L. (2004). Role of f-box factor foxj1 in differentiation of ciliated airway epithelial cells. American Journal of Physiology - Lung Cellular and Molecular Physiology, 286(4 30–4). https://doi.org/10.1152/ajplung.00170.2003

Yu, H., Ye, X., Guo, N., & Nathans, J. (2012). Frizzled 2 and frizzled 7 function redundantly in convergent extension and closure of the ventricular septum and palate: evidence for a network of interacting genes. Development (Cambridge, England), 139(23), 4383–4394. https://doi.org/10.1242/dev.083352

Yu, X., Ng, C. P., Habacher, H., & Roy, S. (2008). Foxj1 transcription factors are master regulators of the motile ciliogenic program. Nature Genetics, 40(12), 1445–1453. https://doi.org/10.1038/ng.263

Zhang, S., Dolgalev, I., Zhang, T., Ran, H., Levine, D. A., & Neel, B. G. (2019). Both fallopian tube and ovarian surface epithelium are cells-of-origin for high-grade serous ovarian carcinoma. Nature Communications, 10(1), 5367. https://doi.org/10.1038/s41467-019-13116-2

Zhang, X. L., Peng, C. J., Peng, J., Jiang, L. Y., Ning, X. M., & Zheng, J. H. (2010). Prognostic role of Wnt7a expression in ovarian carcinoma patients. Neoplasma, 57(6), 545–551. Retrieved from http://www.ncbi.nlm.nih.gov/pubmed/20845993

Zhao, C., Cai, S., Shin, K., Lim, A., Kalisky, T., Lu, W. J., … Beachy, P. A. (2017). Stromal Gli2 activity coordinates a niche signaling program for mammary epithelial stem cells. Science, 356(6335). https://doi.org/10.1126/SCIENCE.AAL3485/SUPPL_FILE/ZHAO.SM.PDF

Zhu, M., Iwano, T., & Takeda, S. (2020). Fallopian tube basal stem cells reproducing the epithelial sheets in vitro—stem cell of fallopian epithelium. Biomolecules, 10(9), 1–15. https://doi.org/10.3390/biom10091270

