## Supplementary Figures for "Single Cell Transcriptomics identifies a WNT7A-FZD5 Signaling Axis that maintains Fallopian Tube Stem Cells in Patient-derived Organoids"

**A**

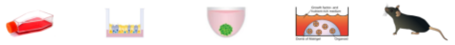

| | 2D culture | Air-Liquid Interface culture | Spheroid culture (suspension) | Organoid culture | *In vivo* models |
|---|---|---|---|---|---|
| **Longevity** | short-term (1-2 months) | short-term | medium | long-term | long-term |
| **Ease of handling** | easy | medium ease | medium ease | medium ease | highly technical |
| **Incubation times** | short | long | medium | medium | very long |
| **Cost** | inexpensive | affordable | inexpensive | affordable | Expensive |
| **Human-derivation** | yes | yes | yes | yes | mouse-specific |
| **Dimensionality** | 2D | 2D | 3D | 3D | 3D |
| **Mature FT features (e.g. invaginations)** | no | no | no | yes | yes |
| **Cell-matrix interactions** | no | limited | no | yes | yes |
| **Epithelial cellular heterogeneity** | no | yes | yes | yes | yes |
| **Non-epithelial microenvironment** | no | may be possible | no | possible | present |
| **Model complexity** | → | | | | |
| **Example studies** | Karst & Drapkin (2012)<br>Shan *et al.* (2012)<br>Jazaeri *et al.*, (2011) | Zhu *et al.* (2020)<br>Levanon *et al.* (2010)<br>Fotheringham *et al.* (2011)<br>(Y. Yamamoto *et al.*, 2016) | Paik *et al.* (2012)<br>Lawrenson *et al.* 2013)<br>Chang *et al.* (2020) | Kessler *et al.* (2015)<br>Xie *et al.* (2018)<br>Hoffmann *et al.*, (2020) | Perets *et al.* (2013)<br>Kim *et al.* (2012)<br>Wang *et al.* (2012)<br>Huang *et al.* (2015) |

**B**

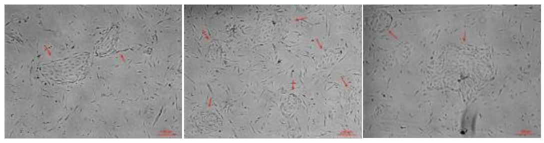

**C**

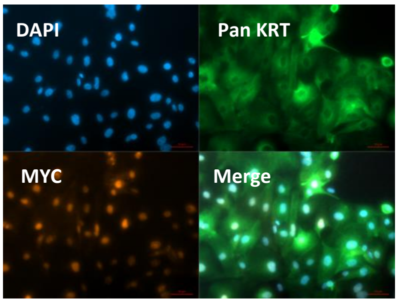


**D**

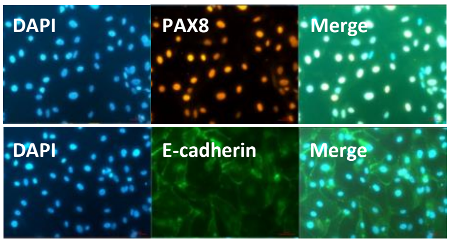


**E**

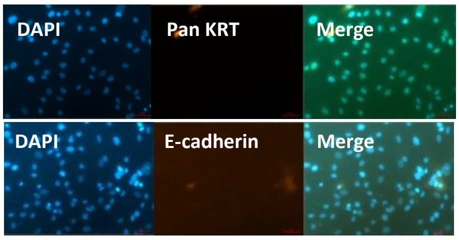


***Figure S1: FT models.*** *(Supplementary to Figure 1A)*

***(A)*** *Table showing the pros (green boxes) and cons of various FT models generated to date. 2D culture of primary FT cells suffers from the drawbacks shown. Spheroid cultures overcome the dimensionality barrier of 2D cultures but generate multi-layered spheroids lacking cell-matrix adhesion, monolayer cell organization and long-term propagation. Air-liquid interface approaches produce well-differentiated cultures that can be effective in modelling stem cell renewal; however they are single-use cultures that cannot be expanded long-term, rely on a continuous supply of patient-derived or murine specimen (Levanon et al., 2010) and expose the epithelium to air; they were originally optimized for airway epithelium (Lin et al., 2007; Pezzulo et al., 2011). In vivo lineage tracing methods are the gold standard for evidencing stem cell activity, but are expensive, highly technical, require protracted incubation times and are not amenable to high throughput approaches. Organoids are a miniaturized form of organs, capturing some of their in vivo complexities while maintaining the convenience of in vitro models.*

***(B)*** *Representative images of primary FT 2D culture on day 3 grown as described in the Methods section. Red arrows point to emerging epithelial 'islands'. Data representative of at least 10 biological replicates (n=10 patients). Scale bar, 200 µm.*

***(C-E)*** *Fluorescence imaging of immunostaining for epithelial cell markers (Pan KRT and E-cadherin) and secretory cell markers (PAX8 and MYC) in primary FT 2D culture < 2 weeks old (C-D) and at ~ day 40* ***(E)****. Ciliated cell markers (TUBB4 and acetylated tubulin) are not expressed in this culture (data not shown). Data representative of 4 biological replicates (n=3 patients). Scar bars, 50 µm.*

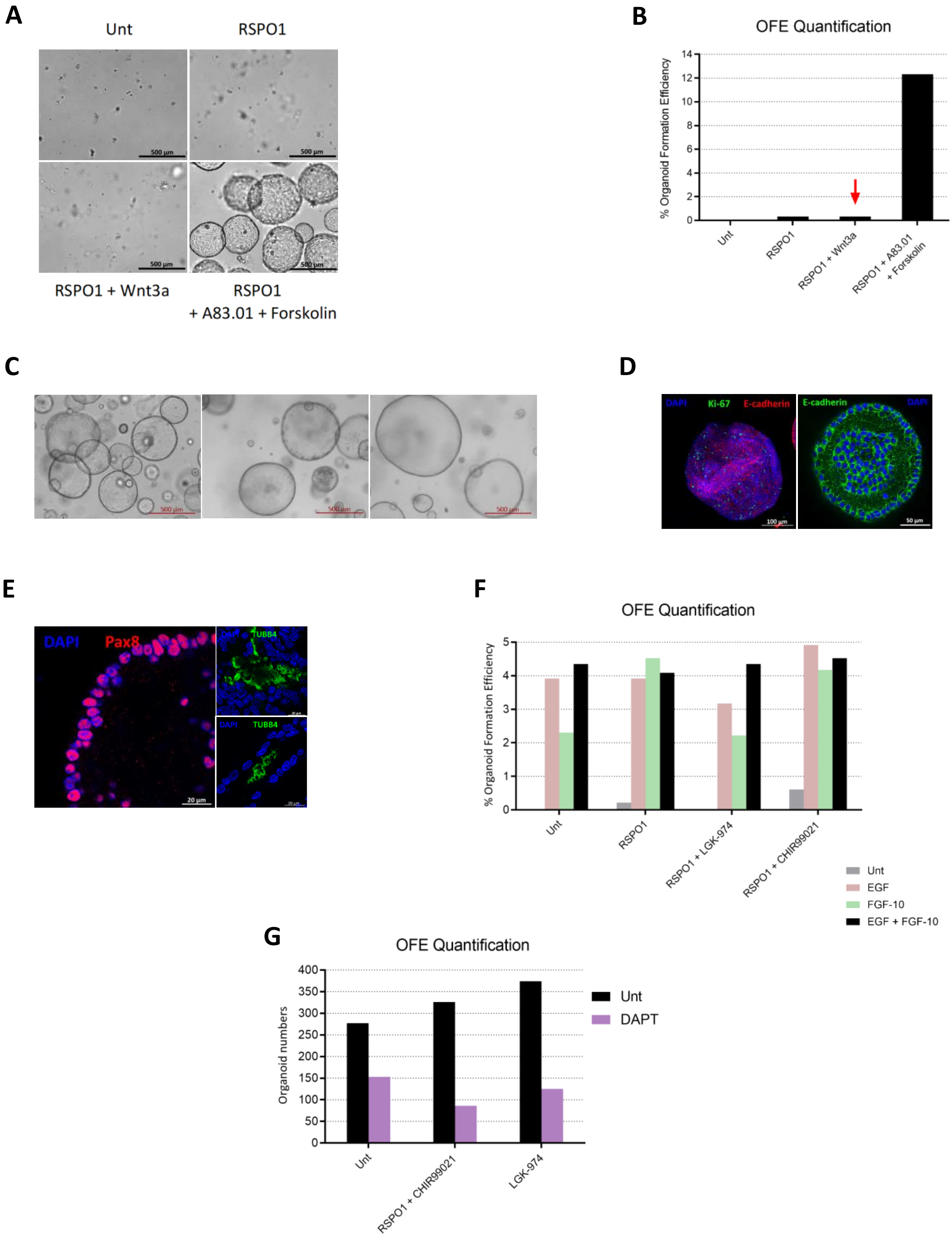
A
Unt
RSPO1
RSPO1 + Wnt3a
RSPO1
+ A83.01 + Forskolin
B
OFE Quantification
% Organoid Formation Efficiency
Unt
RSPO1
RSPO1 + Wnt3a
RSPO1 + A83.01 + Forskolin
C
D
DAPI
Ki-67
E-cadherin
E
Pax8
TUBB4
F
OFE Quantification
% Organoid Formation Efficiency
Unt
RSPO1
RSPO1 + LGK-974
RSPO1 + CHIR99021
Unt
EGF
FGF-10
EGF + FGF-10
G
OFE Quantification
Organoid numbers
Unt
RSPO1 + CHIR99021
LGK-974
Unt
DAPT

***Figure S2: hFT and mouse oviduct (mOV) organoids.***

***(A-B)*** *A83.01 restores organoid formation efficiency (OFE) of single FT cells.* ***(A)*** *Representative brightfield images of EpCAM+ CD45- cells FACS-isolated from p.1 hFT organoid line and cultured 11 days under the treatment conditions shown. 1,200 cells were plated in 50 µl Matrigel drops on 24-well plates. Scale bar, 500 µm.* ***(B)*** *Quantification of OFE for the samples shown in* ***(A)****. Red arrow points to previously reported FT organoid culture methods, which suppress TGF-β signaling using SB431541 at 0.5 µM. This is not sufficient to restore OFE after single cell dissociation or low confluency plating. (Supplementary to Figure 1B-C)*

*Note: A83.01 alone is sufficient to instigate organoid growth after single cell dissociation. Forskolin was shown to significantly improve organoid passage (Huch et al., 2015) and survival of FT-derived cancer (Ince et al., 2015). Effect of Forskolin on organoid passage was not tested in this work.*

***(C-G)*** *mOV organoids regenerate independently of WβS activation and TGF-β suppression. Data shows images* ***(C)****, immunostaining* ***(D-E)*** *and OFE quantification* ***(F-G)*** *of mOV organoids grown in the absence of WβS activators (WNTs, RSPO1, etc; unless indicated) and TGF-β inhibitors (A83.01 and SB431541).*

***(C-E)*** *Similar to hFT organoids, mOV organoids are morphologically spherical* ***(C)*** *and express epithelial* ***(D)*** *and FT-specific markers* ***(E)****.* ***(C)*** *Representative brightfield images of p.1 mOV organoids on day 6. Organoids were established as described in the Methods section, from the oviducts of wild-type female mice, aged 7-12 weeks, C57BL/6 strain. Images are representative of at least 10 biological replicates (n=3 mice). Scale bars as indicated.* ***(D-E)*** *Confocal images of immunostaining for epithelial marker E-cadherin (D) as well as secretory (PAX8) and ciliated (TUBB4) cell markers* ***(E)*** *in whole-mount immunostained mOV organoids. Scale bars as indicated.*

***(F-G)*** *Quantification of mOV OFE after treatment for 11-12 days under the conditions shown.* ***(F)*** *2,300 FACS-sorted cells were plated into 50 µl Matrigel drops on 24-well plates. Colour-coded key shows which growth factor was added in growth factor-free medium.* ***(G)*** *OFE of mOV organoids reduces upon Notch signaling inhibition using the γ-secretase inhibitor DAPT, but not upon WβS activation (RSPO1 + CHIR99021) or inhibition (LGK-974). (C-G supplementary to Figure 1 J-K)*

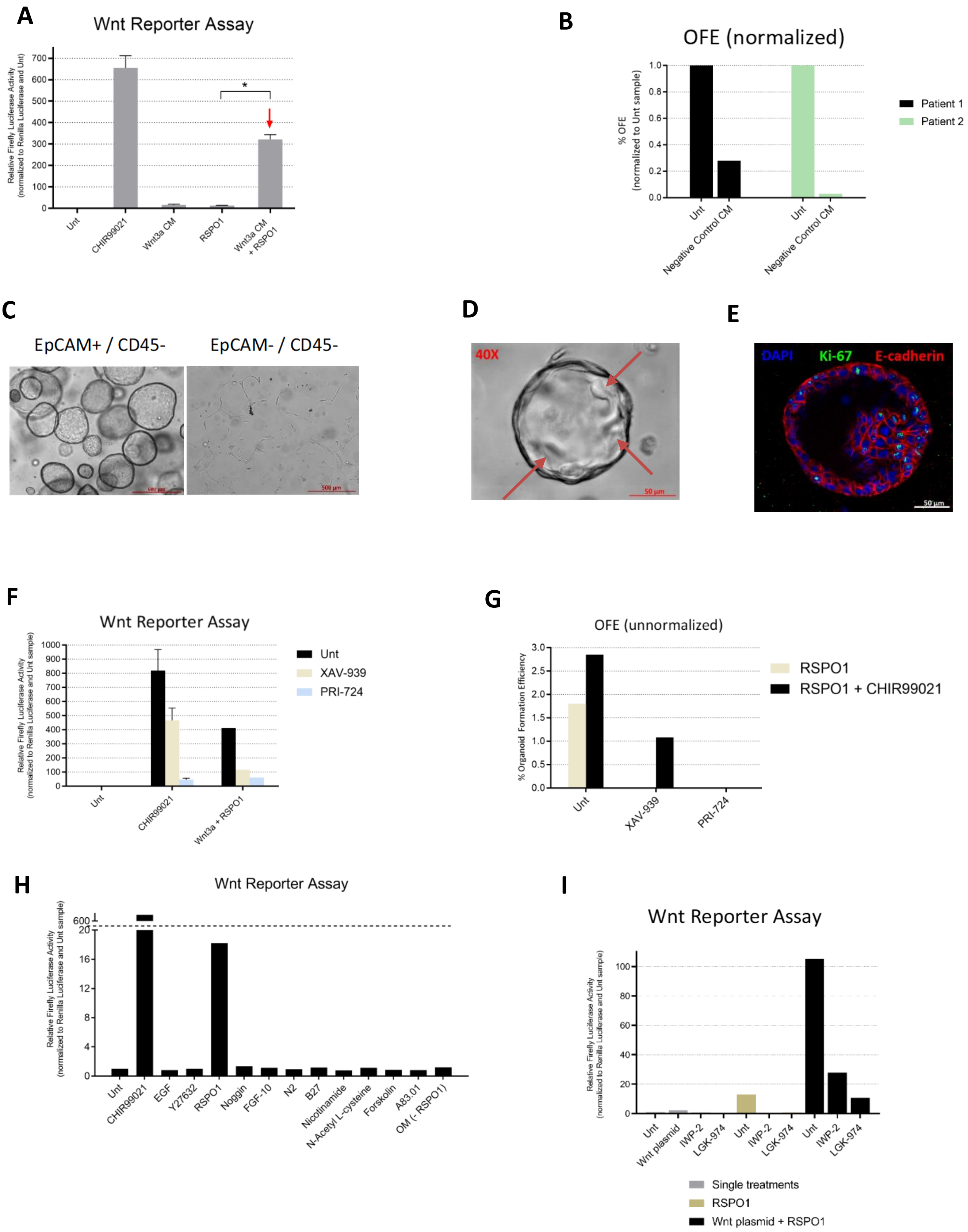

A
Wnt Reporter Assay
Relative Firefly Luciferase Activity
(normalized to Renilla Luciferase and Unt)
700
600
500
400
300
200
100
0
*
Unt
CHIR99021
Wnt3a CM
RSPO1
Wnt3a CM + RSPO1
B
OFE (normalized)
% OFE
(normalized to Unt sample)
1.0
0.8
0.6
0.4
0.2
0.0
Unt
Negative Control CM
Unt
Negative Control CM
Patient 1
Patient 2
C
EpCAM+ / CD45-
EpCAM- / CD45-
500 µm
500 µm
D
40X
50 µm
E
DAPI
Ki-67
E-cadherin
50 µm
F
Wnt Reporter Assay
Relative Firefly Luciferase Activity
(normalized to Renilla Luciferase and Unt sample)
1000
900
800
700
600
500
400
300
200
100
0
Unt
CHIR99021
Wnt3a + RSPO1
Unt
XAV-939
PRI-724
G
OFE (unnormalized)
% Organoid Formation Efficiency
3.0
2.5
2.0
1.5
1.0
0.5
0.0
Unt
XAV-939
PRI-724
RSPO1
RSPO1 + CHIR99021
H
Wnt Reporter Assay
Relative Firefly Luciferase Activity
(normalized to Renilla Luciferase and Unt sample)
600
20
16
12
8
4
0
Unt
CHIR99021
EGF
Y27632
RSPO1
Noggin
FGF-10
N2
B27
Nicotinamide
N-Acetyl L-cysteine
Forskolin
A83.01
OM (- RSPO1)
I
Wnt Reporter Assay
Relative Firefly Luciferase Activity
(normalized to Renilla Luciferase and Unt sample)
100
80
60
40
20
0
Unt
Wnt plasmid
IWP-2
LGK-974
Unt
IWP-2
LGK-974
Unt
IWP-2
LGK-974
Single treatments
RSPO1
Wnt plasmid + RSPO1

***Figure S3: Validating the activity of WβS reagents and their effects on hFT organoids.***
*(Supplementary to Figure 1)*

***(A)*** *Activity of expressed L-WNT3A conditioned medium (CM) was validated using the TOPFlash assay (see Methods for details). WNT3A CM (25%) and RSPO1 (500 ng/mL) were used at the indicated concentrations throughout the study. CHIR99021 was used as positive control for the TOPFlash assay. Error bars represent mean ± SEM for at least 8 biological replicates. *P < 0.0001 (t-test, two-tailed, paired). (Supplementary to Figure 1 H-I)*

***(B)*** *Quantification of OFE of p.1 organoids after treatment with negative control CM for 8-10 days. Data is shown in two biological replicates (n=2 patients). 3,000 cells were plated in 7 µl Matrigel drops on 96-well plates. All samples contained RSPO1.*

***(C)*** *Brightfield images of cells FACS-isolated from patient tissue using the indicated markers, and organoid cultured for 10-14 days. Scale bar, 500 µm.*

***(D)*** *Phase contrast, high magnification image of a hFT organoid. Red arrows point to internal foldings and invaginations characteristic of FT tissue. Scale bar, 50 µm.*

***(E)*** *Confocal image of immunostaining for proliferation marker KI-67 in whole-mount immunostained hFT organoid. Representative of 2-3 biological replicates (n=2 patients). Scale bar, 50 µm. (C-E are supplementary to Figure 1 B-C)*

***(F)*** *Validation of the inhibitory effect of XAV-939 and PRI-724's on WβS using the TOPFlash assay. Error bars represent mean ± SD for 2 biological replicates. Additional data showing this effect is shown later in a different context (Figure S7H). XAV-939 is a tankyrase inhibitor that stabilizes AXIN and the β-catenin Destruction Complex (Huang et al., 2009; Tian et al., 2013). PRI-724 disrupts β-catenin's interaction with its obligate transcriptional co-activator CBP (Okazaki et al., 2019).*

***(G)*** *Quantification of OFE of p.1 organoids after treatment with WβS inhibitors XAV-939 and PRI-724 for 10-12 days. 4,000 cells were plated in 7 µl Matrigel drops on 48-well plates. A second patient replicates is shown in a different context (Figure S8 A-B). (Supplementary to Figure 1G)*

***(H)*** *Testing effect of all reagents in the standard Organoid Medium (OM) on WβS using the TOPFlash assay in HEK293 cells. CHIR99021 is used as positive control here and does not constitute part of the OM medium.*

***(I)*** *Validating the inhibitory effect of Porcupine inhibitors IWP-2 and LGK-974 on WβS using the TOPFlash assay in HEK293 cells. (Supplementary to Figure 1 F-K)*

*In this study, CHIR99021 (3 µM), IWP-2 (3 µM), LGK-974 (2 µM), XAV-939 (5 µM) and PRI-724 (10 µM) were used at the indicated concentrations.*

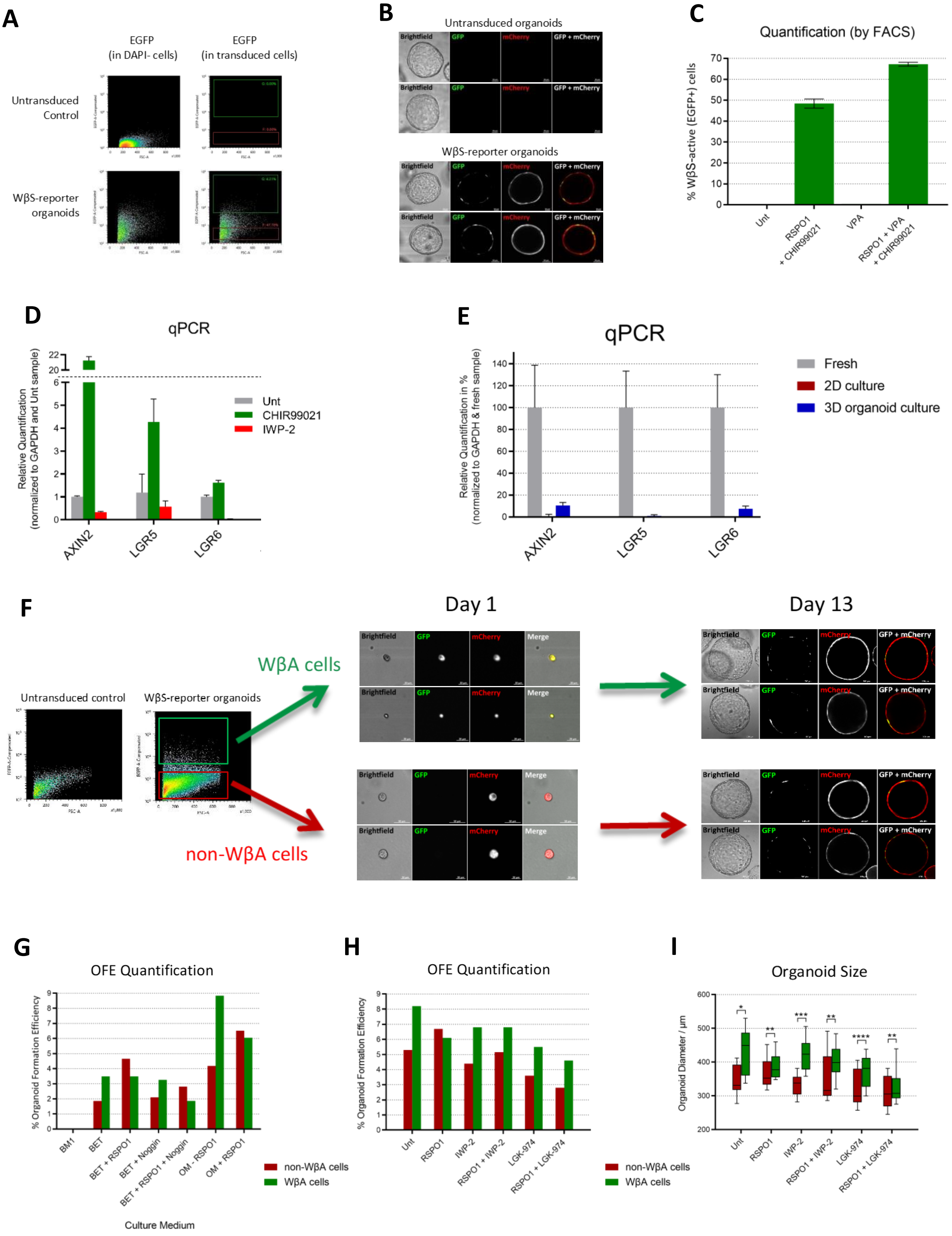
A
EGFP
(in DAPI- cells)
EGFP
(in transduced cells)
Untransduced
Control
WβS-reporter
organoids
B
Untransduced organoids
Brightfield
GFP
mCherry
GFP + mCherry
WβS-reporter organoids
C
Quantification (by FACS)
% WβS-active (EGFP+) cells
Unt
RSPO1
+ CHIR99021
VPA
RSPO1 + VPA
+ CHIR99021
D
qPCR
Relative Quantification
(normalized to GAPDH and Unt sample)
Unt
CHIR99021
IWP-2
AXIN2
LGR5
LGR6
E
qPCR
Relative Quantification in %
(normalized to GAPDH & fresh sample)
Fresh
2D culture
3D organoid culture
AXIN2
LGR5
LGR6
F
Day 1
Day 13
WβA cells
non-WβA cells
Untransduced control
WβS-reporter organoids
Brightfield
GFP
mCherry
Merge
GFP + mCherry
G
OFE Quantification
% Organoid Formation Efficiency
BM1
BET
BET + RSPO1
BET + Noggin
BET + RSPO1 + Noggin
OM - RSPO1
OM + RSPO1
Culture Medium
non-WβA cells
WβA cells
H
OFE Quantification
% Organoid Formation Efficiency
Unt
RSPO1
IWP-2
RSPO1 + IWP-2
LGK-974
RSPO1 + LGK-974
non-WβA cells
WβA cells
I
Organoid Size
Organoid Diameter / µm
Unt
RSPO1
IWP-2
RSPO1 + IWP-2
LGK-974
RSPO1 + LGK-974

***Figure S4: hFT and mOV WβS-reporter organoids.*** *(Supplementary to Figure 2)*

***(A)*** *FACS profile of hFT WβS-reporter organoids. (Supplementary to Figure 2E). Data is representative of at least 10 biological replicates (n = at least 4 patients).*

***(B)*** *Live cell confocal imaging of hFT WβS-reporter organoids after 10-14 days in culture. Images representative of at least 4 patient replicates. Scale bars, 50 μm.*

***(C)*** *FACS analysis of EGFP expression in hFT WβS-reporter organoids treated for 7 days with the conditions shown. Error bars represent mean ± SD for 2-3 technical replicates. Notch activation using Valproic Acid (VPA) potentiates RSPO1 & CHIR99021-induced WβS activation. (Supplementary to Figure 2E)*

***(D)*** *RT-qPCR analysis of the relative mRNA expression of WβS activation markers AXIN2 and LGR5 in hFT organoids treated for 72 hrs with the conditions shown. Error bars represent mean ± 95% confident interval for three technical replicates. AXIN2 and LGR5 were also found to be faithful markers of WβS activation in organoids derived from human intestine, liver, colon, pancreas and stomach (Boonekamp et al., 2021). (Supplementary to Figure 2F)*

***(E)*** *RT-qPCR analysis of the relative mRNA expression of WβS activation markers AXIN2 and LGR5 in fresh hFT cells and patient-matched cultured cells. Error bars represent mean ± 95% confident interval for three technical replicates. WβS activation is more robust in fresh tissue, possibly due to mesenchymal sources of WNT and R-Spondin not present in organoid culture (see Figure 3E).*

***(F)*** *FACS isolation (left panel) and organoid culture (right panel) of mOV WβA or non-WβA cells. Data is representative of 3 biological replicates. Scale bars as indicated. (Supplementary to Figure 2I)*

***(G-I)*** *mOV WβA cells are not enriched in OFE. Shown are quantifications of OFE* ***(G-H)*** *and organoid sizes* ***(I)*** *of WβA or non-WβA cells isolated by FACS from mOV WβS-reporter organoids. 500 cells* ***(G)*** *or 2,000 cells* ***(H)*** *were plated in 50 μl Matrigel drops on 24-well plates. Cells were subjected to the indicated treatments for 7-10 days. For* ***(G)****, BM1 is Basal Medium 1 (DMEM/F12 + HEPES + P/S); BET is BM1 + B27 supplement + EGF + SB431542; OM is complete organoid medium.* ***(I)*** *Box and whisker plot summarizing the sizes of the 10 largest organoids per sample shown in* ***(H)****; centre line, median; box limits, upper and lower quartiles; whisker ends, minimum and maximum values). P-values = *0.007, **n.s., ***<0.0001, ****<0.05 (t-test, two-tailed). While not possessing enhanced OFE, WβA cells here produce slightly larger organoids under Unt and WNT blocking conditions. (G-I are supplementary to Figure 2J)*

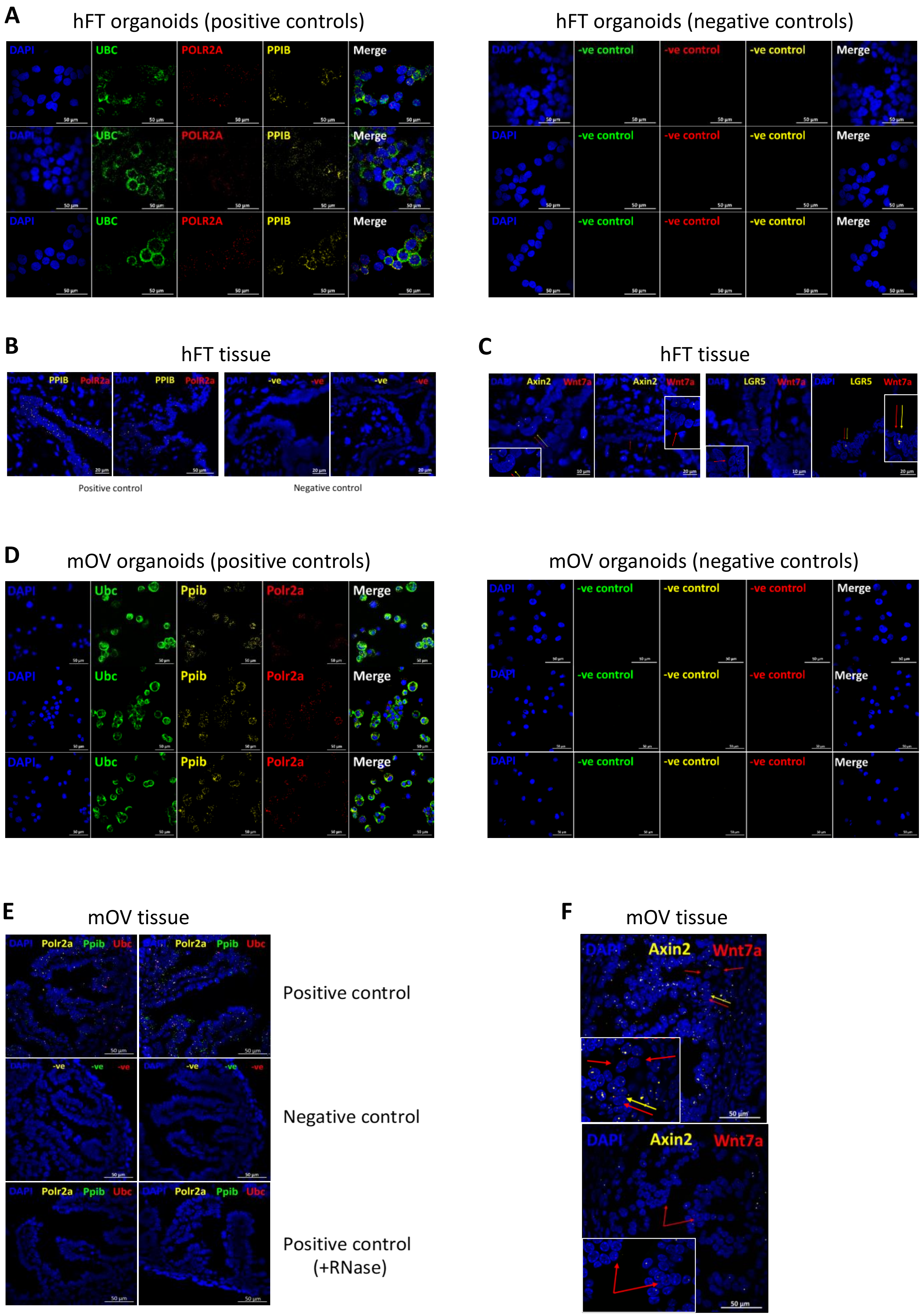
A
hFT organoids (positive controls)
DAPI
UBC
POLR2A
PPIB
Merge
hFT organoids (negative controls)
DAPI
-ve control
-ve control
-ve control
Merge
B
hFT tissue
DAPI PPIB PolR2a
DAPI -ve -ve
Positive control
Negative control
C
hFT tissue
DAPI Axin2 Wnt7a
DAPI LGR5 Wnt7a
D
mOV organoids (positive controls)
DAPI
Ubc
Ppib
Polr2a
Merge
mOV organoids (negative controls)
DAPI
-ve control
-ve control
-ve control
Merge
E
mOV tissue
DAPI Polr2a Ppib Ubc
DAPI -ve -ve -ve
Positive control
Negative control
Positive control
(+RNase)
F
mOV tissue
DAPI Axin2 Wnt7a
50 µm

***Figure S5: RNA FISH staining.*** *(Supplementary to Figure 3 B-C)*

***(A)*** *Confocal images of fluorescent RNAScope FISH staining performed on hFT organoids using positive control (left panel) or negative control (right panel) probes. Scale bars, 50 µm. (Staining controls for Figure 3B)*

***(B-C)*** *Confocal images of fluorescent RNAScope FISH staining performed on fresh frozen sections of hFT tissue derived from a non-ovarian cancer patient. Staining performed using control probes* ***(B)*** *or AXIN2, LGR5 or WNT7A probes* ***(C)****. For* ***(B)****, positive control signal is abolished by RNase treatment (data not shown). Insets in* ***(C)*** *show enlarged parts of punctate RNA pointed to by arrows that are colour-matched to gene names. Scale bars as indicated. (Supplementary to Figure 3B)*

***(D)*** *Confocal images of fluorescent RNAScope FISH staining performed on mOV organoids using positive control (left panel) or negative control (right panel) probes. Scale bars, 50 µm. (Staining controls for Figure 3C)*

***(E-F)*** *Confocal images of fluorescent RNAScope FISH staining performed on fresh frozen sections of mOV tissue derived from C57BL/6 female mice aged 7-12 weeks.* ***(E)*** *Positive control signal is abolished by RNase treatment, confirming punctate dots detected are RNA. Insets in* ***(F)*** *show enlarged parts of punctate RNA pointed to by arrows that are colour-matched to gene names. Scale bars, 50 µm. (Supplementary to Figure 3C)*

*For all control FISH stainings (****A****,* ***B****,* ***D****,* ***E****), UBC/Ubc, PPIB/Ppib and POLR2A/Polr2a probes were used to target ubiquitous / housekeeping genes expressed at high, medium or low levels, respectively. Human and mouse positive control probes target the same genes using human-specific or mouse-specific probes. Negative control probes target bacterial sequences. Organoid samples* ***(A)*** *and* ***(C)*** *were dissociated, cytospinned and fixed prior to RNA FISH staining. -ve is negative control probes. All data representative of at least 2-3 biological replicates. Scale bars as indicated.*

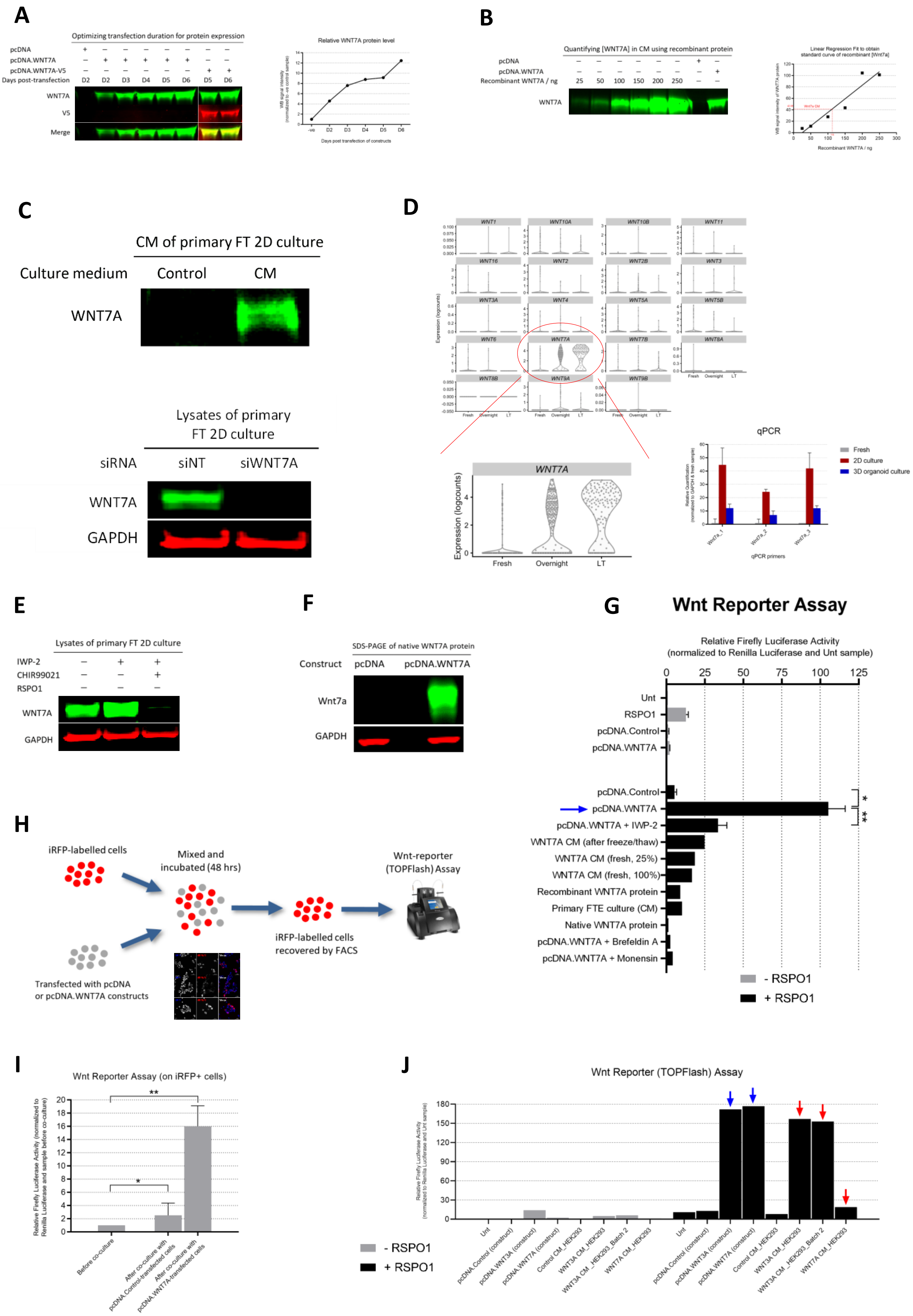
A
Optimizing transfection duration for protein expression
pcDNA
pcDNA.WNT7A
pcDNA.WNT7A-V5
Days post-transfection
D2 D2 D3 D4 D5 D6 D5 D6
WNT7A
V5
Merge
Relative WNT7A protein level
Days post transfection of constructs
B
Quantifying [WNT7A] in CM using recombinant protein
pcDNA
pcDNA.WNT7A
Recombinant WNT7A / ng
25 50 100 150 200 250
WNT7A
Linear Regression Fit to obtain standard curve of recombinant [Wnt7a]
Recombinant WNT7A / ng
C
CM of primary FT 2D culture
Culture medium
Control
CM
WNT7A
Lysates of primary FT 2D culture
siRNA
siNT
siWNT7A
WNT7A
GAPDH
D
WNT1 WNT10A WNT10B WNT11
WNT16 WNT2 WNT2B WNT3
WNT3A WNT4 WNT5A WNT5B
WNT6 WNT7A WNT7B WNT8A
WNT8B WNT9A WNT9B
Expression (logcounts)
Fresh Overnight LT
WNT7A
Expression (logcounts)
Fresh
Overnight
LT
qPCR
Fresh
2D culture
3D organoid culture
qPCR primers
E
Lysates of primary FT 2D culture
IWP-2
CHIR99021
RSPO1
WNT7A
GAPDH
F
SDS-PAGE of native WNT7A protein
Construct
pcDNA
pcDNA.WNT7A
Wnt7a
GAPDH
G
Wnt Reporter Assay
Relative Firefly Luciferase Activity
(normalized to Renilla Luciferase and Unt sample)
0 25 50 75 100 125
Unt
RSPO1
pcDNA.Control
pcDNA.WNT7A
pcDNA.Control
pcDNA.WNT7A
pcDNA.WNT7A + IWP-2
WNT7A CM (after freeze/thaw)
WNT7A CM (fresh, 25%)
WNT7A CM (fresh, 100%)
Recombinant WNT7A protein
Primary FTE culture (CM)
Native WNT7A protein
pcDNA.WNT7A + Brefeldin A
pcDNA.WNT7A + Monensin
- RSPO1
+ RSPO1
H
iRFP-labelled cells
Mixed and incubated (48 hrs)
Wnt-reporter (TOPFlash) Assay
iRFP-labelled cells recovered by FACS
Transfected with pcDNA or pcDNA.WNT7A constructs
I
Wnt Reporter Assay (on iRFP+ cells)
Relative Firefly Luciferase Activity (normalized to Renilla Luciferase and sample before co-culture)
0 2 4 6 8 10 12 14 16 18 20
Before co-culture
After co-culture with pcDNA.Control-transfected cells
After co-culture with pcDNA.WNT7A-transfected cells
J
Wnt Reporter (TOPFlash) Assay
Relative Firefly Luciferase Activity (normalized to Renilla Luciferase and Unt sample)
0 30 60 90 120 150 180
- RSPO1
+ RSPO1
Unt
pcDNA.Control (construct)
pcDNA.WNT3A (construct)
pcDNA.WNT7A (construct)
Control CM_HEK293
WNT3A CM_HEK293
WNT3A CM _HEK293_Batch 2
WNT7A CM_HEK293

***Figure S6: Expression and validation of WNT7A protein reagents.*** *(Supplementary to Figure 3)*

***(A)*** *Western blot of denatured conditioned medium harvested from HEK293 cells that were transfected with the indicated WNT7A constructs for the durations shown. Left panel shows WNT7A immunoblot. V5 signal confirms specificity of the WNT7A antibody. Right panel is a quantification of relative band intensities.*

***(B)*** *Quantification of [WNT7A] in CM harvested on day 3 (see* ***A****) using recombinant WNT7A protein (R&D). Left panel shows immunoblot; right panel is a quantification of band intensity to generate a calibration curve for estimating [WNT7A] in CM.* ***(A-B)*** *are representative of three biological replicates.*

***(C-D)*** *Primary 2D-cultured FT cells strongly overexpress WNT7A protein* ***(C)*** *and mRNA* ***(D)****.* ***(C)*** *Western blot for WNT7A in the CM (upper panel) and lysates (lower panel) of primary 2D-cultured FT cells. For upper panel, CM was harvested after 4-day medium incubation with 1-week old primary FT culture. Control here is the same medium prior to incubation with cells. Upper and lower panel are for different patients. WNT7A knockdown in lower panel confirms specificity of the WNT7A antibody used (also confirmed in* ***A****).* ***(D)*** *Upper panel shows violin plots for the single cell expression profile of the WNT family of ligands in fresh or 2D-cultured FT cells (overnight or LT; long-term, 2 or 6 days). Each dot represents one cell. Bottom panel (left) is an enlargement of the circled plot above. Bottom panel (right) is an RT-qPCR analysis showing relative WNT7A mRNA levels in fresh or cultured (2D or organoid culture) FT cells in a patient-matched sample, using 3 distinct WNT7A qPCR primers. Error bars represent mean ± 95% confident interval for three technical replicates.*

***(E)*** *WNT7A protein detectable in lysates of primary 2D-cultured FT cells is almost completely abolished upon WβS pathway activation, hinting that WβS activation and WNT7A expression may be mutually exclusive cell states. Data in this figure are for the same patient as (****C****, lower panel).*

***(F)*** *Isolation of native WNT7A protein using non-denaturing non-ionic detergent (Passive Lysis Buffer, Promega) from lysates of HEK293 cells that were reverse transfected with the shown constructs for 48 hrs. The blot shown here is after SDS-PAGE, confirming presence of WNT7A in lysate extract.*

***(G)*** *WNT7A protein reagents are not functional. Shown are TOPFlash readings in HEK293 cells. Blue arrow points to induction of WβS by the WNT7A construct. Error bars represent mean ± SEM for at least 3 biological replicates (9 biological replicates for the sample with blue arrow). P-values = *0.0005, **0.009 (t-test, two-tailed). Some samples are normalized to internal controls not shown for simplicity.*

***(H-I)*** *WNT7A has a short-range signal.* ***(H)*** *Set up of co-culture assay.* ***(I)*** *Quantification of TOPFlash signal induced in iRFP+ cells before or after co-culture with control or WNT7A construct. Error bars represent mean ± SD for 2 biological replicates. P-values = * n.s.; **<0.05 (t-test, two-tailed).*

***(J)*** *TOPFlash assay to test activity of HEK293-secreted WNT7A and WNT3A proteins. Blue arrows point to robust TOPFlash signal induced by WNT3A and WNT7A constructs. While WNT7A CM derived from the same construct backbone is not functional, WNT3A CM shows robust activity (red arrows).*

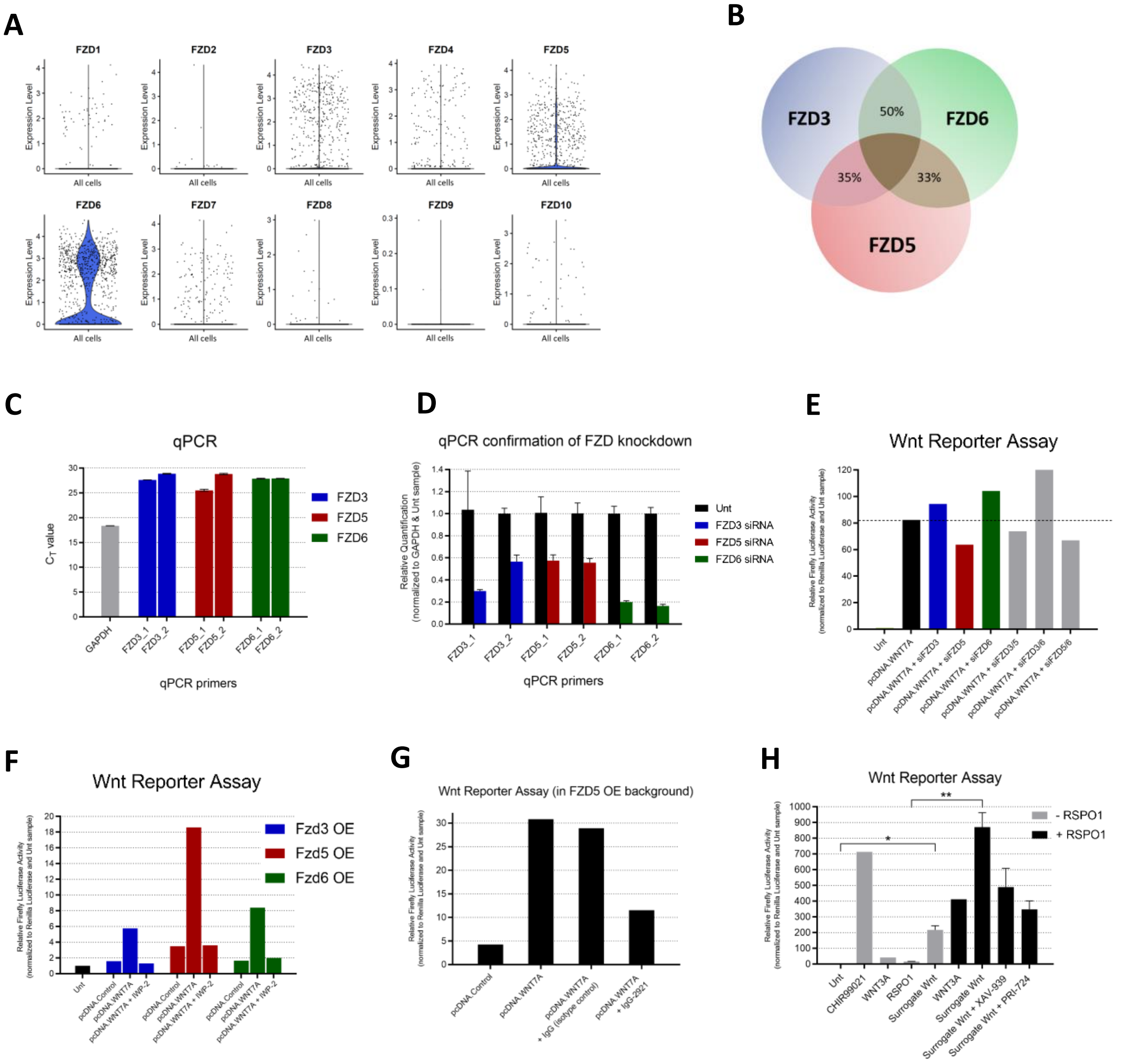
A
FZD1
FZD2
FZD3
FZD4
FZD5
FZD6
FZD7
FZD8
FZD9
FZD10
Expression Level
All cells
B
FZD3
FZD6
FZD5
50%
35%
33%
C
qPCR
$C_T$ value
GAPDH
FZD3_1
FZD3_2
FZD5_1
FZD5_2
FZD6_1
FZD6_2
qPCR primers
FZD3
FZD5
FZD6
D
qPCR confirmation of FZD knockdown
Relative Quantification (normalized to GAPDH & Unt sample)
Unt
FZD3 siRNA
FZD5 siRNA
FZD6 siRNA
qPCR primers
E
Wnt Reporter Assay
Relative Firefly Luciferase Activity (normalized to Renilla Luciferase and Unt sample)
Unt
pcDNA.WNT7A
pcDNA.WNT7A + siFZD3
pcDNA.WNT7A + siFZD5
pcDNA.WNT7A + siFZD6
pcDNA.WNT7A + siFZD3/5
pcDNA.WNT7A + siFZD3/6
pcDNA.WNT7A + siFZD5/6
F
Wnt Reporter Assay
Fzd3 OE
Fzd5 OE
Fzd6 OE
pcDNA.Control
pcDNA.WNT7A
pcDNA.WNT7A + IWP-2
G
Wnt Reporter Assay (in FZD5 OE background)
pcDNA.Control
pcDNA.WNT7A
pcDNA.WNT7A + IgG (isotype control)
pcDNA.WNT7A + IgG-2921
H
Wnt Reporter Assay
- RSPO1
+ RSPO1
Unt
CHIR99021
WNT3A
RSPO1
Surrogate Wnt
Surrogate Wnt + XAV-939
Surrogate Wnt + PRI-724

***Figure S7: FZD5 is a receptor for WNT7A.*** *(Supplementary to Figure 4)*

***(A)*** *Violin plots showing the single cell expression profile of the FZD family of receptors. Fresh EpCAM+ CD45- cells were FACS-isolated from hFT tissue (Hu et al., 2020) and subjected to SMART-seq2 protocol (Picelli et al., 2014). Each dot represents one cell. (Supplementary to Figure 4A)*

***(B)*** *Protein sequence similarity between hFT-expressed FZDs, obtained by pairwise sequence alignments of RefSeq sequences.*

***(C)*** *Relative expression levels of FZDs 3/5/6 in HEK293 cells estimated by $C_T$ values obtained from an RT-qPCR analysis, using two independent qPCR primers for each FZD. GAPDH $C_T$ value is shown for comparison. Error bars represent mean ± SD for three technical replicates. Data is representative of two biological replicates.*

***(D)*** *RT-qPCR analysis of the relative mRNA expression of FZDs 3/5/6 in HEK293 cells after siRNA knockdown for 72 hrs. Error bars represent mean ± 95% confident interval for three technical replicates.*

***(E-F)*** *TOPFlash assay showing the effect of gene knockdown **(E)** and overexpression **(F)** of FZDs 3/5/6 on WNT7A-induced WβS in HEK293 cells. Indicated constructs (200 ng) and siRNAs (40 nM) were transfected for 72 hrs. All samples contained RSPO1 (24 hr treatment), except Unt sample in **(E)**. Data in **(C-E)** are from the same experiment. IWP-2 treatment in **(F)** confirms this signal is due to WNT7A OE.*

***(G)*** *TOPFlash assay showing the effect of anti-FZD5 antibody IgG-2921 on WβS in a background of WNT7A and Fzd5 overexpression in HEK293 cells. Constructs were transfected for 72 hrs. Antibody treatments were done for 24 hrs. Given the Fzd5 OE construct encodes mouse Fzd5, this also shows that IgG-2921 is active against mouse FZD5 protein. (Supplementary to Figure 4 C-D and F-G)*

***(H)*** *TOPFlash assay validating the activity of Surrogate Wnt (Janda et al., 2017). All treatments performed for 24 hrs in HEK293 cells. Surrogate Wnt-induced TOPFlash signal reduces with XAV-939 and PRI-724 treatment, confirming that Surrogate Wnt functions through the canonical WβS machinery. Error bars represent mean ± SD for 2 biological replicates. P-values = *<0.05, **0.0056 (t-test, two-tailed). (Supplementary to Figure 4 C-E and Figure S8)*

*Surrogate Wnt is a synthetic bivalent peptide containing an antibody epitope that binds the extracellular FZD cysteine-rich domain (CRD). Surrogate Wnt also contains a linker sequence and an LRP5/6-binding motif (Janda et al., 2017). The FZD-CRD epitope in Surrogate Wnt is the single chain variable fragment of OMP-18R5, while the LRP5/6-binding motif is the C-terminal portion of DKK1 (Janda et al., 2017). Overall, Surrogate Wnt selectively hetero-dimerizes FZDs with LRP6 co-receptor extracellularly, mimicking the action of endogenous WNTs.*

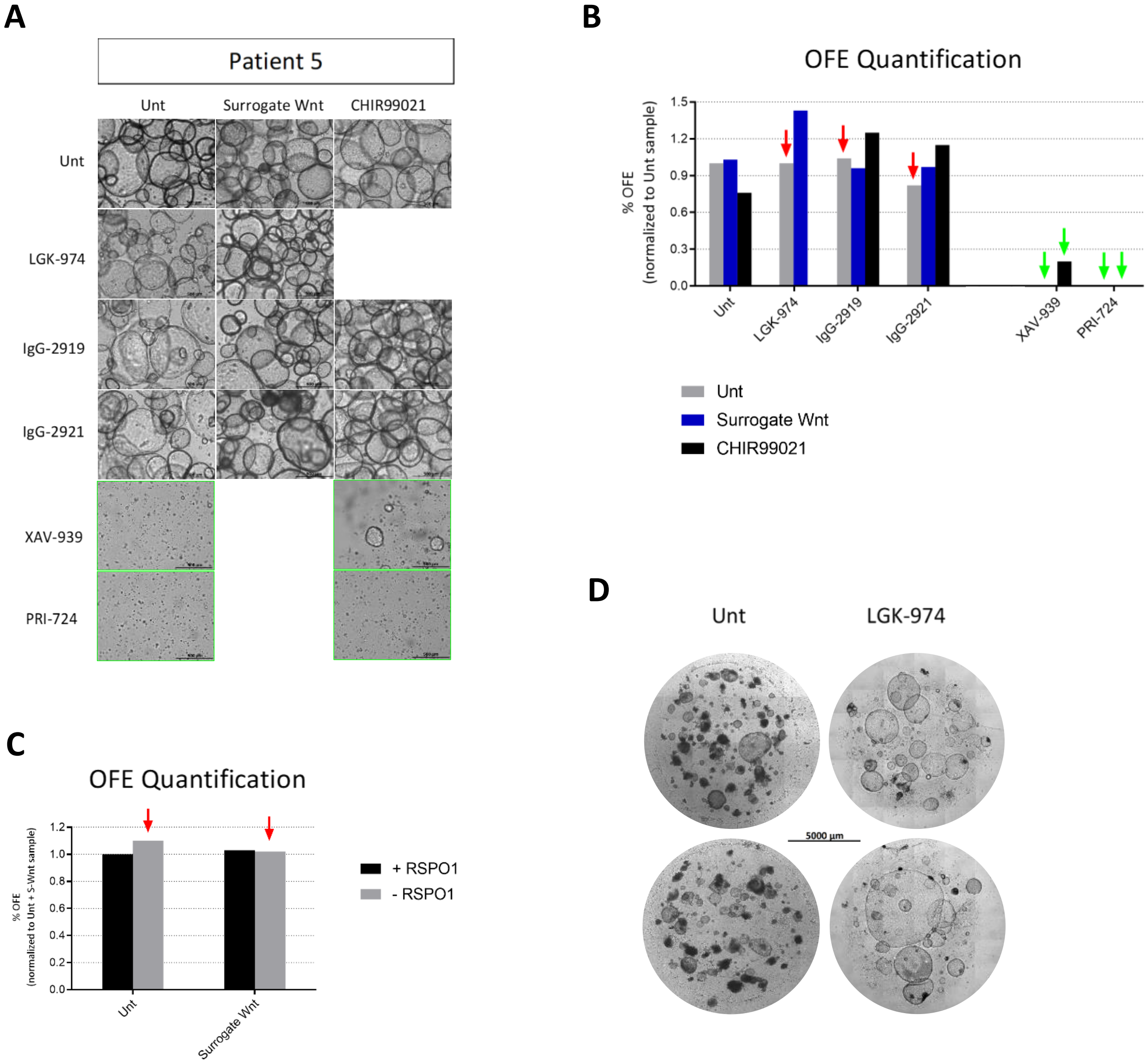
A
Patient 5
Unt
Surrogate Wnt
CHIR99021
Unt
LGK-974
IgG-2919
IgG-2921
XAV-939
PRI-724
B
OFE Quantification
% OFE
(normalized to Unt sample)
1.5
1.2
0.9
0.6
0.3
0.0
Unt
LGK-974
IgG-2919
IgG-2921
XAV-939
PRI-724
Unt
Surrogate Wnt
CHIR99021
C
OFE Quantification
% OFE
(normalized to Unt + S-Wnt sample)
1.2
1.0
0.8
0.6
0.4
0.2
0.0
Unt
Surrogate Wnt
+ RSPO1
- RSPO1
D
Unt
LGK-974
5000 µm

***Figure S8: Patient 5 organoids are resistant to inhibition of the WNT7A-FZD5 signaling axis.*** *(Supplementary to Figure 4 C-E)*

***(A)*** *Representative brightfield images of Patient 5 p.1 organoids treated for 10-13 days under the conditions shown. 3-4K cells were plated in 7 µl Matrigel drops. All samples contained RSPO1. Scale bars, 500 µm. Although resistant to blocking endogenous WNTs and FZD5, Patient 5 organoids are sensitive to downstream WβS inhibitors (images with green outline) as seen in other patients (see Figure S3G). (Supplementary to Figure 4C)*

***(B)*** *Quantification of OFE for the samples shown in* ***(A)****. Red arrows point to treatments that abolish organoid regeneration in all other tested patients. Green arrows point to treatments with downstream WβS inhibitors. (Supplementary to Figure 4D)*

***(C)*** *Quantification of OFE in Surrogate Wnt-treated organoids upon RSPO1 withdrawal. Red arrows point to treatments that abolish OFE in other patients. (Supplementary to Figure 4E)*

*For (****A-C****), Patient 5 organoids were tested in parallel with Patient 3 organoids on the same day using the same inhibitor: culture medium mixes. Patient 3 organoids showed sensitivity to all inhibitors of the WNT7A-FZD5 signaling axis (Figure 4 C-D).*

***(D)*** *Brightfield whole-well images of Patient 5 organoids after growing for 4 passages under the conditions shown, allowing organoids to propagate for 3 weeks without medium change to replenish stem cell factors. This normally leads to differentiation or organoid death (Unt, left panel). Patient 5 organoids expanded under selective WNT-blocked (LGK-974) conditions and continued robust growth across passages, indicating growth advantage and ectopic independence from stem cell niche requirements. Targeted DNA sequencing showed Patient 5 organoids harbour no TP53 mutations.*

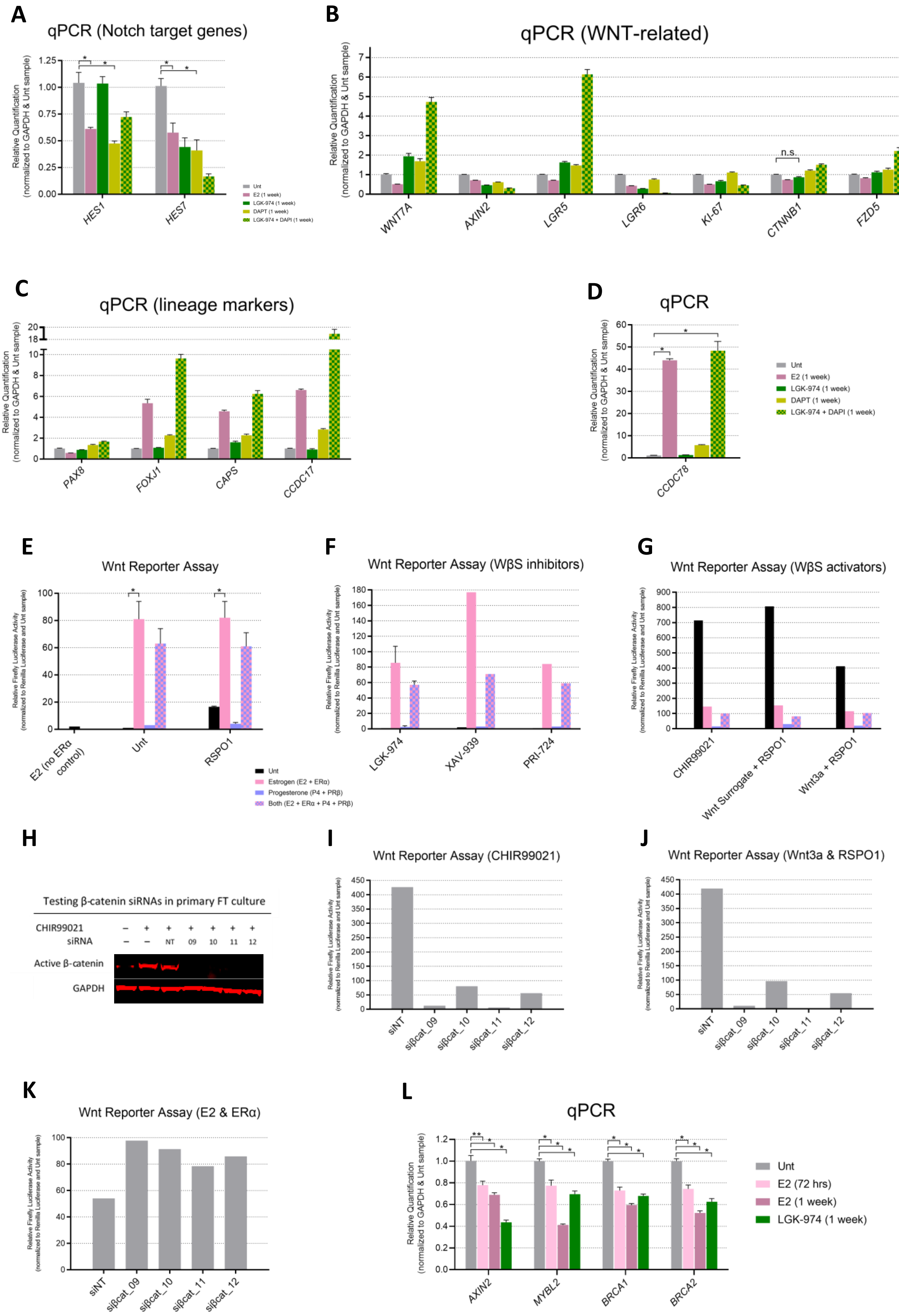
A
qPCR (Notch target genes)
Relative Quantification (normalized to GAPDH & Unt sample)
HES1
HES7
Unt
E2 (1 week)
LGK-974 (1 week)
DAPT (1 week)
LGK-974 + DAPI (1 week)
B
qPCR (WNT-related)
Relative Quantification (normalized to GAPDH & Unt sample)
n.s.
WNT7A
AXIN2
LGR5
LGR6
KI-67
CTNNB1
FZD5
C
qPCR (lineage markers)
Relative Quantification (normalized to GAPDH & Unt sample)
PAX8
FOXJ1
CAPS
CCDC17
D
qPCR
Relative Quantification (normalized to GAPDH & Unt sample)
CCDC78
Unt
E2 (1 week)
LGK-974 (1 week)
DAPT (1 week)
LGK-974 + DAPI (1 week)
E
Wnt Reporter Assay
Relative Firefly Luciferase Activity (normalized to Renilla Luciferase and Unt sample)
E2 (no ERα control)
Unt
RSPO1
Unt
Estrogen (E2 + ERα)
Progesterone (P4 + PRβ)
Both (E2 + ERα + P4 + PRβ)
F
Wnt Reporter Assay (WβS inhibitors)
Relative Firefly Luciferase Activity (normalized to Renilla Luciferase and Unt sample)
LGK-974
XAV-939
PRI-724
G
Wnt Reporter Assay (WβS activators)
Relative Firefly Luciferase Activity (normalized to Renilla Luciferase and Unt sample)
CHIR99021
Wnt Surrogate + RSPO1
Wnt3a + RSPO1
H
Testing β-catenin siRNAs in primary FT culture
CHIR99021
siRNA
NT 09 10 11 12
Active β-catenin
GAPDH
I
Wnt Reporter Assay (CHIR99021)
Relative Firefly Luciferase Activity (normalized to Renilla Luciferase and Unt sample)
siNT
siβcat_09
siβcat_10
siβcat_11
siβcat_12
J
Wnt Reporter Assay (Wnt3a & RSPO1)
Relative Firefly Luciferase Activity (normalized to Renilla Luciferase and Unt sample)
siNT
siβcat_09
siβcat_10
siβcat_11
siβcat_12
K
Wnt Reporter Assay (E2 & ERα)
Relative Firefly Luciferase Activity (normalized to Renilla Luciferase and Unt sample)
siNT
siβcat_09
siβcat_10
siβcat_11
siβcat_12
L
qPCR
Relative Quantification (normalized to GAPDH & Unt sample)
AXIN2
MYBL2
BRCA1
BRCA2
Unt
E2 (72 hrs)
E2 (1 week)
LGK-974 (1 week)

***Figure S9: Estrogen's molecular influences on WNT7A, WβS and differentiation in hFT and HEK293 cells*** *(Supplementary to Figure 5)*

***(A-D)*** *Estrogen induces a ciliogenesis differentiation programme in hFT organoids through inhibition of WβS and Notch signaling. RT-qPCR analysis of the relative mRNA expression of:* ***(A)*** *Notch target genes;* ***(B)*** *WNT7A, AXIN2, LGR6 and other WNT-related genes shown;* ***(C-D)*** *secretory and ciliated cell markers. For* ***(A-D)****, data are for the same samples and treatments were administered to expanded organoids 7-9 days after organoid passage. Estrogen (17β-estradiol, 100 nM), LGK-974 (2 µM) or DAPT (10 µM) were introduced at the indicated concentrations for a period of 1 week. Error bars represent mean ± SEM 3-12 technical replicates (n = 2 patients). Asterisks denote statistical significance calculated using unpaired student's t-test, as follows:* ***(A)*** **P < 0. 0.0072;* ***(B)*** *For each gene, statistical pairwise comparison between Unt and any other sample; P < 0.003, except for the shown comparison (n.s.; not significant);* ***(C)*** *For each gene, statistical pairwise comparison between Unt vs E2, Unt vs DAPT or Unt vs LGK-974 + DAPT; P < 0.00073;* ***(D)*** **P < 0.0001. Additional 3-6 technical replicates for the effect of estrogen on the indicated genes are shown in Figure 5 in a different context.* ***(B)*** *is supplementary to Figure 5F.* ***(C)*** *is supplementary to Figure 5G.* ***(D)*** *is supplementary to Figure 5H.*

***(E-G)*** *TOPFlash assay showing the effect of estrogen or/and progesterone on WβS in cells treated for 24 hrs under the conditions indicated. Error bars represent mean ± SEM for 2 biological replicates. *P-value < 0.05 (t-test, two-tailed).*

***(H-J)*** *β-catenin siRNAs effectively abolish β-catenin protein level* ***(H)*** *and activity* ***(I-J)****.* ***(H)*** *Western blot for active β-catenin (dephosphorylated at Ser37 and Thr41) using lysates from 2D-cultured primary FT cells that were transfected with four independent β-catenin siRNAs for 72 hrs.* ***(I-J)*** *TOPFlash assay showing the effect of β-catenin siRNA's on WβS induced by CHIR99021* ***(I)*** *and Wnt3a+RSPO1* ***(J)****.*

***(K)*** *TOPFlash assay showing estrogen's activation of WβS in a background of β-catenin siRNA knockdown. Data in* ***(I-K)*** *were normalized to Unt controls (not shown) and obtained from the same experiment using the same β-catenin siRNA-containing transfection mixes. For all Wnt Reporter / TOPFlash assays in this figure (****E-G*** *and* ***I-K****), HEK293 cells were used. Estrogen treatment refers to administering 17β-estradiol / E2 (100 nM, 24 hrs) on cells transfected with ERα construct (200 ng) for 48 hrs. Progesterone treatment refers to P4 treatment (1 µM, 24 hrs) on cells transfected with PRβ construct (200 ng) for 48 hrs. For the estrogen + progesterone condition, these treatments / constructs were introduced simultaneously.*

***(L)*** *RT-qPCR analysis of the relative mRNA expression of the indicated genes in hFT organoids that were treated as indicated. Error bars represent mean ± SEM for 3-9 technical replicates (n=2 patients). * P < 0.004 and ** P = 0.02 (t-test, unpaired).*

**A**

| Patient | # WβA cells | # non-WβA cells | Total cell no. (per patient) |
|---|---|---|---|
| 11579 | 110 | 270 | 380 |
| 15149 | 176 | 170 | 346 |
| 11600 | 156 | 139 | 295 |
| Total cell no. (per cell type / state) | 442 | 579 | 1,021 (Total cell no.) |

**B**

Patient 11597, WβA vs non-WβA cells, WβS-related GSEA pathways

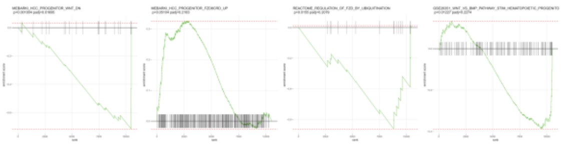

**C**

Patient 15149, WβA vs non-WβA cells, WβS-related GSEA pathways

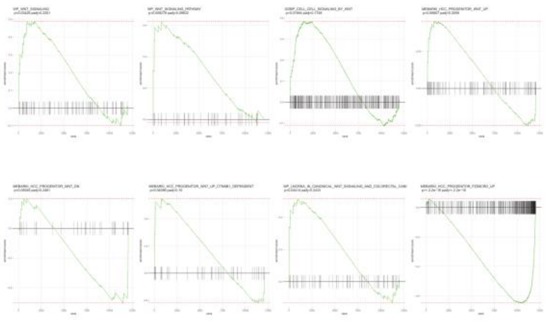

**D**

Patient 11600, WβA vs non-WβA cells, WβS-related GSEA pathways

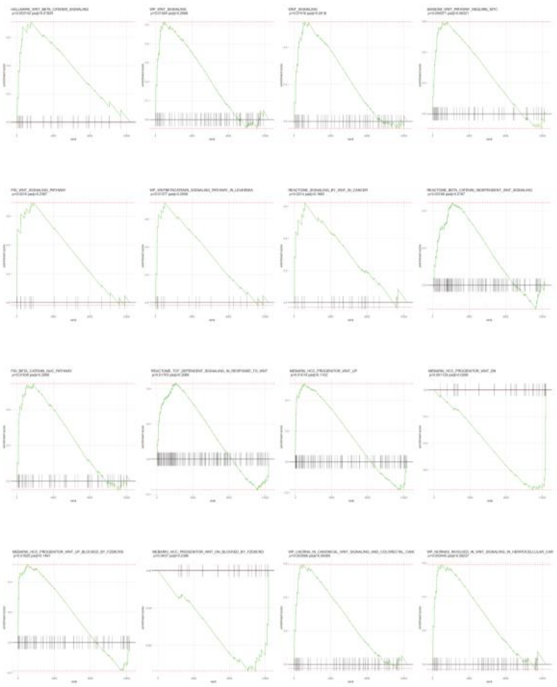

**E**

Patient 11597, WβA vs non-WβA cells, ECM-related GSEA pathways

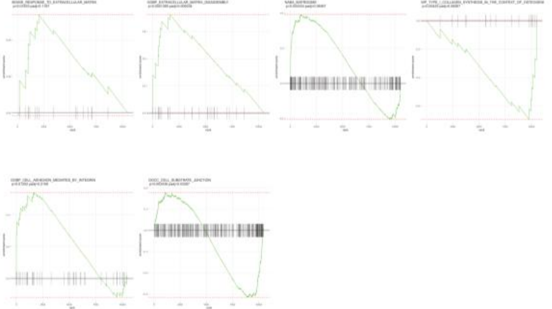

**F**

Patient 15149, WβA vs non-WβA cells, ECM-related GSEA pathways

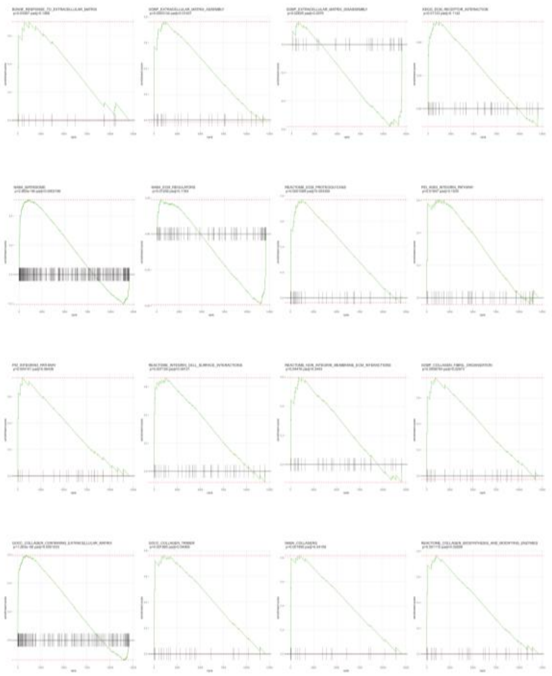

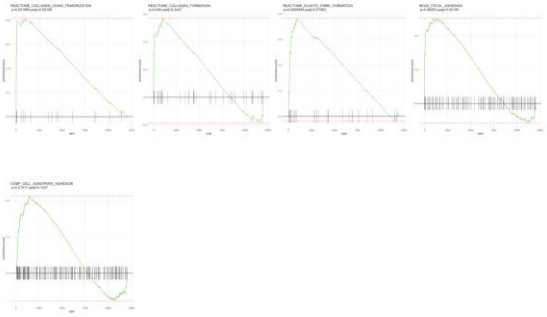

**G**

Patient 11600, WβA vs non-WβA cells, ECM-related GSEA pathways

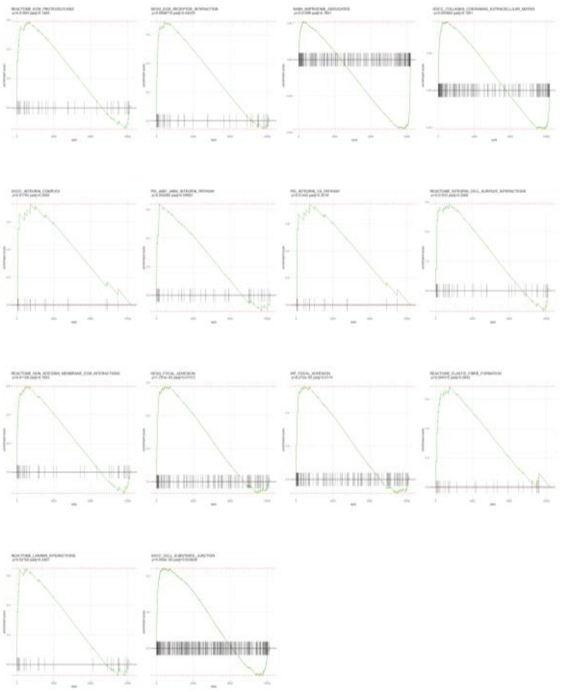

***Figure S10: Further single cell transcriptomic characterization of WβA cells.***
*(Supplementary to Figure 6)*

***(A)*** *Table showing the number of cells per patient and per cell type (GFP+ WβA cells vs GFP- non-WβA cells) that were FACS-isolated from patient-derived hFT WβS-reporter organoids and processed using the SMART-seq2 protocol for scRNA-seq.*

***(B-F)*** *GSEA enrichment plots from intra-patient comparisons of the single cell transcriptomes of WβA vs non-WβA cells from* ***(A)****, showing enrichment of WβS-related* ***(B-D)*** *and ECM-related* ***(E-G)*** *pathways in patients 11597* ***(B, E)****, 15149* ***(C, F)*** *and 11600* ***(D, G)****.*

### LIST OF REFERENCES


Boonekamp, K. E., Heo, I., Artegiani, B., Asra, P., van Son, G., de Ligt, J., & Clevers, H. (2021). Identification of novel human Wnt target genes using adult endodermal tissue-derived organoids. *Developmental Biology*. https://doi.org/10.1016/j.ydbio.2021.01.009

Hu, Z., Artibani, M., Alsaadi, A., Wietek, N., Morotti, M., Shi, T., … Ahmed, A. A. (2020). The Repertoire of Serous Ovarian Cancer Non-genetic Heterogeneity Revealed by Single-Cell Sequencing of Normal Fallopian Tube Epithelial Cells. *Cancer Cell*, *37*(2), 226-242.e7. https://doi.org/10.1016/j.ccell.2020.01.003

Huang, S. M. A., Mishina, Y. M., Liu, S., Cheung, A., Stegmeier, F., Michaud, G. A., … Cong, F. (2009). Tankyrase inhibition stabilizes axin and antagonizes Wnt signalling. *Nature*, *461*(7264), 614–620. https://doi.org/10.1038/nature08356

Janda, C. Y., Dang, L. T., You, C., Chang, J., Lau, W. De, Zhong, Z. A., … Garcia, K. C. (2017). Surrogate Wnt agonists that phenocopy canonical Wnt and β-catenin signalling. *Nature*, *545*(7653), 234–237. https://doi.org/10.1038/nature22306

Levanon, K., Ng, V., Piao, H. Y., Zhang, Y., Chang, M. C., Roh, M. H., … Drapkin, R. (2010). Primary ex vivo cultures of human fallopian tube epithelium as a model for serous ovarian carcinogenesis. *Oncogene*, *29*(8), 1103–1113. https://doi.org/10.1038/onc.2009.402

Lin, H., Li, H., Cho, H. J., Bian, S., Roh, H. J., Lee, M. K., … Kim, D. D. (2007). Air-Liquid Interface (ALI) culture of human bronchial epithelial cell monolayers as an in vitro model for airway drug transport studies. *Journal of Pharmaceutical Sciences*, *96*(2), 341–350. https://doi.org/10.1002/jps.20803

Okazaki, H., Sato, S., Koyama, K., Morizumi, S., Abe, S., Azuma, M., … Nishioka, Y. (2019). The novel inhibitor PRI-724 for Wnt/β-catenin/CBP signaling ameliorates bleomycin-induced pulmonary fibrosis in mice. *Experimental Lung Research*, *45*(7), 188–199. https://doi.org/10.1080/01902148.2019.1638466

Pezzulo, A. A., Starner, T. D., Scheetz, T. E., Traver, G. L., Tilley, A. E., Harvey, B. G., … Zabner, J. (2011). The air-liquid interface and use of primary cell cultures are important to recapitulate the transcriptional profile of in vivo airway epithelia. *American Journal of Physiology - Lung Cellular and Molecular Physiology*, *300*(1), 25–31. https://doi.org/10.1152/ajplung.00256.2010

Tian, X. H., Hou, W. J., Fang, Y., Fan, J., Tong, H., Bai, S. L., … Li, Y. (2013). XAV939, a tankyrase 1 inhibitior, promotes cell apoptosis in neuroblastoma cell lines by inhibiting Wnt/β-catenin signaling pathway. *Journal of Experimental and Clinical Cancer Research*, *32*(1). https://doi.org/10.1186/1756-9966-32-100
